# Generative epigenetic landscapes map the topology and topography of cell fates

**DOI:** 10.1101/2025.06.09.658705

**Authors:** Victoria Mochulska, Paul François

**Affiliations:** Department of Physics, McGill University, Montréal, QC, Canada; Département de Biochimie et Medecine Moléculaire, Université de Montréal, Montréal, QC, Canada; Mila - Quebec Artificial Intelligence Institute

## Abstract

Epigenetic landscapes were proposed by Waddington as the central concept to describe cell fate dynamics in a locally low-dimensional space. In modern landscape models, attractors represent cell types, and stochastic jumps and bifurcations drive cellular decisions, allowing for quantitative and predictive descriptions. However, given a biological problem of interest, we still lack tools to infer and build possible Waddington landscapes systematically. In this study, we propose a generative model for deriving epigenetic landscapes compatible with data. To build the landscapes, we combine gradient and rotational vector fields composed of locally weighted elements that encode ‘valleys’ of the Waddington landscape, resulting in interpretable models. We optimize landscapes through computational evolution and illustrate our approach with two developmental examples: metazoan segmentation and neuromesoderm differentiation. In both cases, we obtain ensembles of solutions that reveal both known and novel landscapes in terms of topology and bifurcations. Conversely, topographic features appear strongly constrained by dynamical data, which suggests that our approach can generically derive interpretable and predictive epigenetic landscapes.

## Introduction

During development, cells gradually assume a specialized identity through a process called differentiation. Cellular phenotypes spontaneously change and go through a sequence of well-defined states. Traditionally, cell states have been characterized phenotypically, e.g., by their morphology and function [1], but with the advancement of high-throughput data acquisition, cell states are now defined molecularly, by patterns of gene expression (e.g., abundance of mRNAs and synthesized proteins [2, 3]). Theoretical studies have confirmed that such data allow for the definition of unambiguous cell states [4, 5].

Because of this, cellular differentiation can be abstracted as a dynamics in a high-dimensional ‘gene’ space [6], thus raising the question of the underlying rules governing such dynamics. Remarkably, *cellular decisions* appear to be (locally) tree-like and binary [1], a fact first formalized by Waddington, who proposed the concept of *epigenetic landscape*. In this view, the cell is envisioned as a ball rolling downhill, ‘canalized’ into branching valleys leading to distinct fates. Multiple underlying weights ‘shape’ the landscape and could potentially change based on signalling [7]. Waddington’s analogy can now be more rigorously tied to data and modern mathematical frameworks, such as gene network descriptions [8, 9]. However, complex, emergent landscapes are difficult to capture with descriptions focused only on a few genes [10], suggesting that more abstract descriptions are needed. An alternative strategy is to coarse-grain complex, high-dimensional gene expressions to represent cellular dynamics in a space of possible cell states, leading to the development of the so-called *landscape models* [11–16]. Such landscape descriptions further enable rigorous mathematical analysis and classifications of decisions, leveraging dynamical systems and catastrophe theory: for instance, binary decisions can be classified into either the binary choice or the heteroclinic flip [12, 17]. It is, however, unclear how to systematically build such models for a given biological problem. Practically, the current strategy is to assume or infer a particular decision structure, then connect polynomial normal forms to fit data, such as the statistics of cell states as a function of time [14, 16]. However, such descriptions involve explicit or implicit choices (e.g., on the nature of bifurcations), and it would be valuable to have a broader sense of the family of landscapes and bifurcations consistent with a given problem. Machine learning techniques can help, e.g., by learning families of potentials from a projection of high-dimensional gene expression data [18], or by reconstructing flow in high-dimensional spaces [3]. Such methods come with other issues: they typically require a large amount of data, are not transparent, and might not allow for full landscape inference from canalized trajectories characteristic of development.

Here, we propose a general, versatile, and intuitive formalism to systematically construct landscape models with minimal assumptions. We achieve an unconstrained landscape topology by using simple ‘building blocks’, referred to as modules, which can be added and optimized. In current landscape models [13, 14], the focus is on decisions: each building block corresponds to a developmental choice, written with polynomial normal forms, so the *topology* of decisions is imposed. We propose a dual approach, focused on cell states: our building blocks encode valleys in the epigenetic landscape, written with Gaussian potentials, so that the *topography* of the landscape is optimized. Importantly, in our proposed formalism, the topology of the landscape *emerges* from the topography, and we show that very similar topographies can give rise to different topologies.

Our manuscript is organized as follows. We first introduce the modular formalism for constructing landscapes and demonstrate its duality with previous approaches [12] by recovering standard biological decision-making processes, and further extend our framework to other types of biological dynamics and bifurcations. We then introduce the optimization process allowing us to build landscapes, and proceed to revisit two problems involving cell fate decisions: a landscape ‘screen’ of animal segmentation coupled to growth, and fitting the dynamics of neuromesodermal differentiation. For both cases, we show how our approach extends the previously obtained results, we identify common features and differences between possible solutions and explore how additional constraints can modulate the results. Our work shows how landscape approaches can be used to efficiently explore the space of possible biological dynamics and generate new predictions. Our algorithm for landscape construction and optimization, **Evoscape**, is available on Github: https://github.com/victoria-mochulska/evoscape.

## Results

### Optimizing landscapes with interpretable modules

We consider cellular differentiation in a two-dimensional phenotypic space, where the coordinate 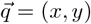 describes the cell state. Motion in this space corresponds to changes in cell state and is given by a flow field: 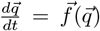. To construct a general flow field 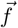, we include both a gradient component, representing the downhill motion, and a rotational component. In 2D, these components can be written using two scalar potential functions, which we define as the gradient potential 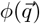 and the rotational potential 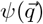. Additionally, we include a global confining potential term *ϕ*_0_:

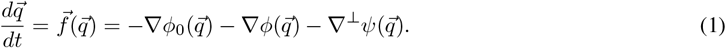

Here, the skew gradient operator ∇^⊥^ is the usual gradient combined with a rotation of 90 degrees. For the gradient potential *ϕ*, the valleys (local minima) are attractive and the hills (local maxima) are repellent. Meanwhile, for the rotational potential *ψ*, the valleys give local clockwise (left) rotation and the hills counterclockwise (right) rotation (Fig. 1).

**Figure 1:**
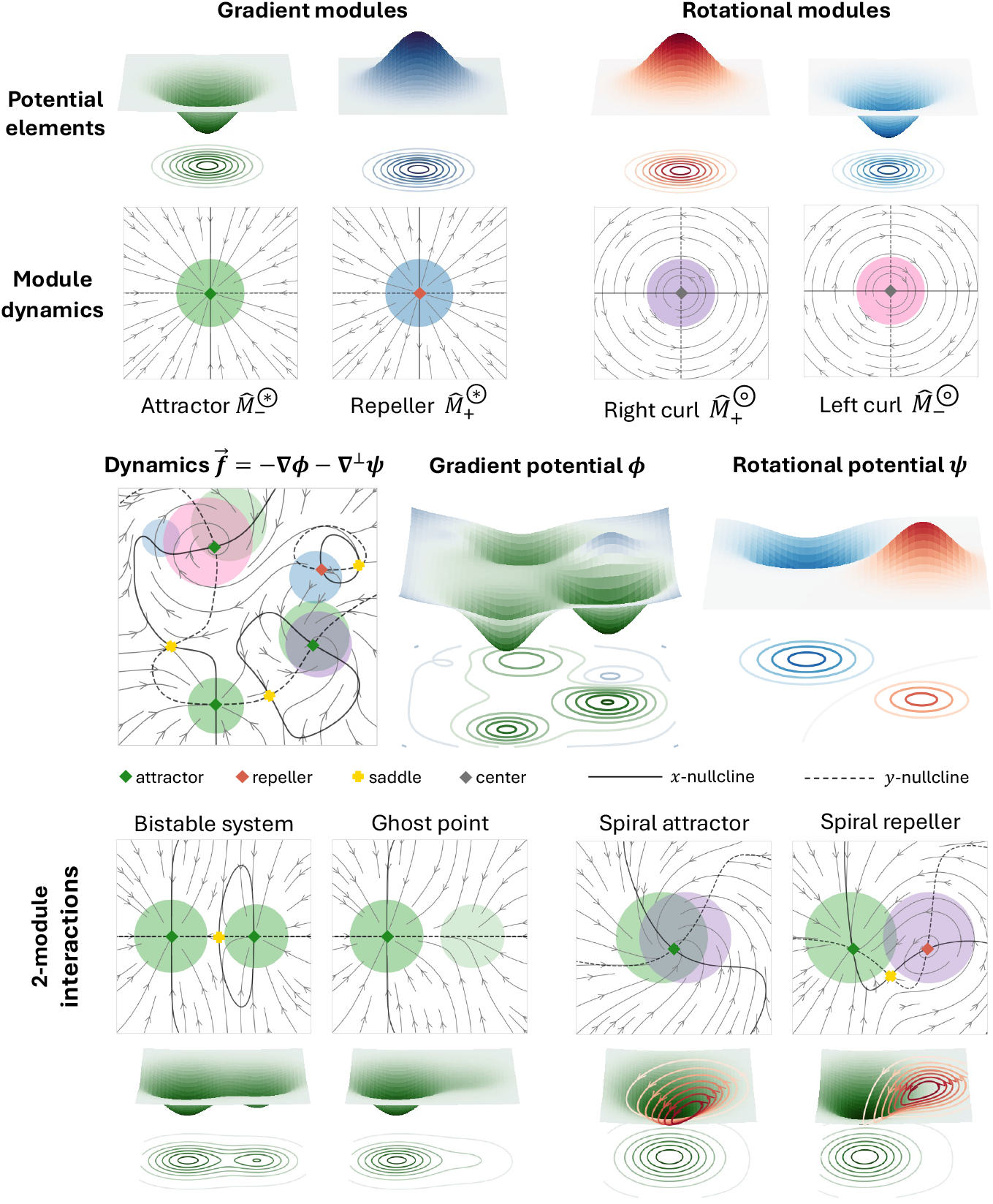
Modular landscape construction. Two classes of local dynamics modules are defined: gradient and rotational. Each module contributes a Gaussian element to the corresponding scalar potential. In vector field plots, modules are visualized as circles (radius: full width at half maximum of the Gaussian, opacity: module strength). Nullclines are shown as black lines: solid for *x*, dashed for *y*. Multiple modules are combined into a landscape, described by the dynamics 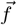 or, equivalently, the gradient potential *ϕ* and rotational potential *ψ*. Examples illustrate local interactions. For spiral interactions, the rotational potential is shown as contours on the surface of the gradient potential.

An important aspect of cell state dynamics is stochasticity, with gene expression subject to both intrinsic and extrinsic noise [19, 20]. We can incorporate noise as a stochastic perturbation to the underlying deterministic dynamical system. Specifically, we use Langevin dynamics with additive, uncorrelated Gaussian noise:

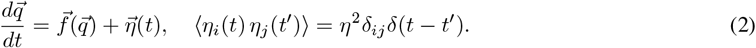

The *landscape* refers to the deterministic structure 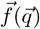. Cellular decisions – transitions between attractors – can occur through deterministic flows or stochastic fluctuations.

### Module definition

To construct the landscape, we loosely follow the original analogy of Waddington, where cell states correspond to valleys. We thus combine local features such as hills and valleys in both scalar potentials, using radial functions:

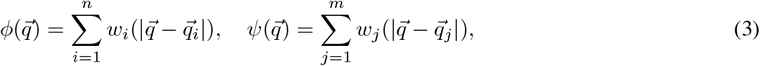

where 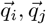 are the locations of landscape features. The functions *w*_*i*_ should be smooth and vanishing at infinity, 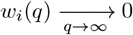, to be able to control the shape of the landscape locally. We choose Gaussian functions as the basis:

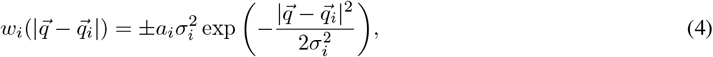

where a minus sign is used for valleys and a plus sign for hills. The amplitude of the Gaussian 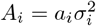 is chosen for convenience in parametrizing the flow 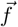, as shown below. Taking the gradients in Eq. 1, the vector field 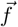 becomes a sum of local dynamics with the confining field:

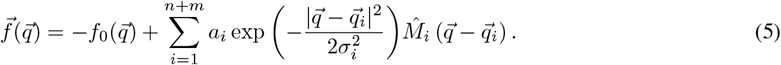

We refer to each element of the sum as a dynamics module (Fig. 1). A module comprises a local spatial weight, here Gaussian, and the linear dynamics given by the matrix 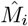. Each module is characterized by its parameters: location 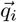 strength *a*_*i*_, size *σ*_*i*_, as well as one of the four possible behaviors 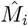 attractor 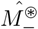, repeller 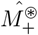 left rotator 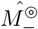, and right rotator 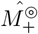 (see Supplement, Section S1.2).

On its own or remote from other modules, each module creates a fixed point of a corresponding type: an attractor or repeller for gradient modules, and a center for rotational modules. For an isolated module, the fixed point is located at 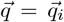 and has eigenvalues *λ*_1_, *λ*_2_, |*λ*_1_| = |*λ*_2_| = *a*_*i*_. The interactions of two closely spaced modules are also intuitive (Fig. 1). In a bistable system, a saddle forms naturally between attractors, while weak attractor modules create ghost points, proposed to drive directed transient dynamics [21]. An overlap of a gradient module with a rotational one creates a spiral. Combining multiple modules allows us to build up the complexity of the landscape and its bifurcations while controlling each local feature independently, as we illustrate in the following subsections.

### Landscape model captures bifurcations ubiquitous in biology

We aim to fit cellular decision dynamics as generically as possible. As a sanity check, we first show that our formalism can easily recover known models of cellular decisions. Binary cell state decisions are ubiquitous in development. They can be modelled as a gradient-like landscape with three attractors: the initial pluripotent state, and the two alternative, more differentiated fates. Two distinct decision structures are possible: the double cusp (binary choice) landscape and the heteroclinic flip landscape [12, 22, 14]. In the double cusp landscape, the origin attractor can undergo a saddle-node bifurcation with either of the saddles leading to fate attractors. In contrast to the local saddle-node, a global flip bifurcation reroutes cells between the two possible fates in the heteroclinic flip landscape [12].

We use three attractor modules, initially of the same strength and size. As shown in Fig. 2A-B (middle panels), we can recover both the double cusp and the heteroclinic flip landscapes by adjusting the relative positions of the modules. Another way to obtain bifurcations of a landscape is to fix the module locations 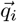 while varying the weight parameters *a*_*i*_ and *σ*_*i*_. The landscape’s attractors - corresponding to defined phenotypes - are located approximately at locations 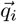 in the phenotypic space. Holding 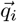 constant with respect to time or external signals thus allows us to fix the identity of cell states. Here, we use this approach to maintain stable attractor identities and control their basins of attraction and bifurcations through *a*_*i*_ and *σ*_*i*_. Details on parameterization are available in the Supplement, Section S1.3.

**Figure 2:**
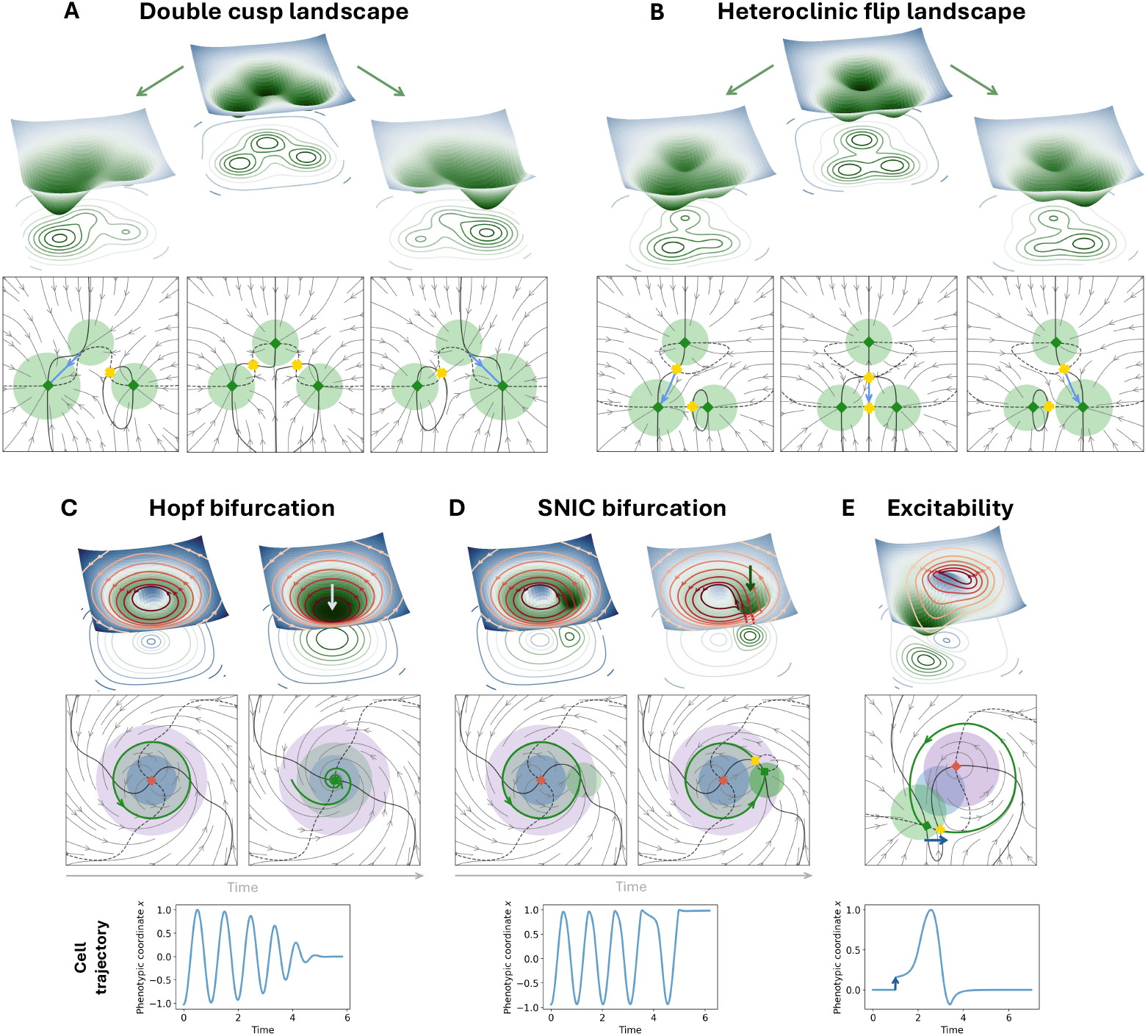
Constructed landscapes for biological dynamics. **A-B)** Gradient landscapes for binary cell fate decisions. **A)** Double cusp landscape: the central attractor is connected by saddles to two peripheral attractors. Starting from a symmetric configuration (middle), the central attractor can bifurcate with either saddle, forcing cells to move to one of the fate attractors (left and right panels, respectively). This is achieved by changing the size of the corresponding fate module. The escape trajectory is shown in blue. **B)** Heteroclinic flip landscape: the escape route from the central attractor can flip between the fate attractors when their sizes are changed. In the middle panel, the escape trajectory (blue) lands exactly on the saddle between fates, creating a heteroclinic connection. **C-D)** Landscapes with non-gradient dynamics. **C)** A limit cycle oscillator is created by combining an attractor, a repeller, and a rotation. The 3D plot shows the gradient potential and the contour lines of the rotation potential. Hopf bifurcation: decreasing the amplitude of the repeller shrinks the limit cycle into a spiral point. **D)** SNIC bifurcation: a smaller attractor module is added close to the limit cycle; increasing its amplitude slows down the oscillation and eventually creates a saddle-node pair on the cycle. **E)** An excitable system: when a cell escapes the attractor in the blue arrow direction (by perturbation or stochastically), it makes a long excursion before returning to the attractor. The loop is created using a repeller and a rotation.

To obtain more complex dynamics and bifurcations, we incorporate non-gradient (rotational) modules. Oscillations are widespread in development, driving differentiation of multiple structures from arthropod segments [23, 24] and vertebrae [25] to fish scales [26]. The simplest limit cycle oscillator is a circular valley overlaid with a rotation and can be built with three modules (Fig. 2C): a repeller, a broader attractor, and a rotational flow. We then construct examples of bifurcations that have been proposed to create or destroy the limit cycle in biological oscillators [27–29]. Hopf bifurcation is obtained by shrinking the repeller. SNIC (saddle-node in cycle) is obtained by adding a small attractor module close to the limit cycle trajectory and gradually increasing its strength (Fig. 2D). Analytic derivations of normal forms for the Hopf and SNIC bifurcations are presented in the Supplement, S2.2-S2.3. Homoclinic bifurcation can be obtained similarly to SNIC by adding an attractor module, but placing it further from the limit cycle. Notice that past the bifurcation, one can get other biologically relevant behaviors, such as Type I excitability (Fig. 2E) [30].

### Evolutionary optimization

We use our modular landscape formalism to fit biologically relevant dynamics from the ground up. For fitting, we use an evolutionary algorithm (Fig. 3), mimicking biology, but also taking advantage of the modularity of our landscape by dynamically adding or removing modules. The set of modules and their parameters then acts as a genotype that can be randomly mutated. The complete evolutionary procedure is described in the Supplement. Briefly, we define mutation operators: module addition, module deletion, and parameter modification (for location or weight parameters). A population of landscapes is set up, then mutated and selected over a number of generations. Selection is performed according to a fitness function, which can be based on the dynamics produced by the landscape or the outcomes of these dynamics regarding cell fate. For the fitness calculation, we simulate the trajectories of cells in each landscape, starting from an initial state. Depending on the biological system of interest, the dynamics outcomes to consider are the obtained cell fates, number of populated cell states, cell distribution across attractors, the spatial pattern of cell fates, etc. More details on the optimized functions are given in the subsequent sections.

**Figure 3:**
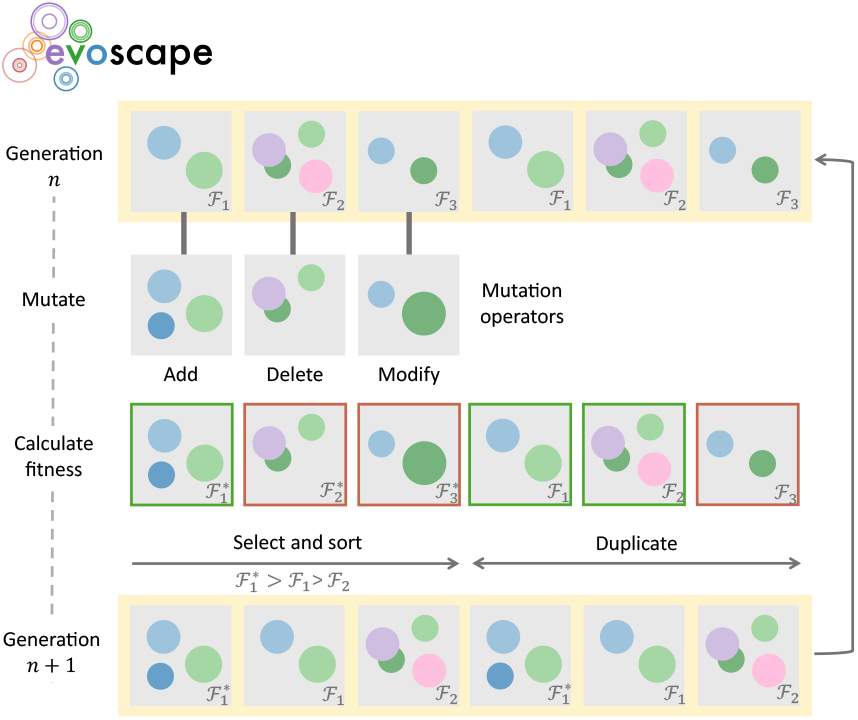
Evolutionary algorithm implemented in Evoscape. In each iteration of the algorithm, half of the landscapes are randomly mutated, and the other half are carried over from the previous generation. After fitness evaluation, the best half (green outlines) is selected and duplicated to form the next generation.

### A phenotypic landscape view of metazoan segmentation

As a first example, we proceed with the optimization of a model of metazoan segmentation, such as somitogenesis in vertebrates [31], Fig. 4A, left. The development of vertebrae begins with segmenting a tissue called the presomitic mesoderm (PSM) into blocks, or somites. The formation of somites is periodic and sequential, with new somites being added one by one from anterior to posterior, and occurs as the embryonic tail is growing. PSM cells exhibit oscillations of gene expression, which stop when cells form a future somite (with physical boundaries forming and separating somites). Within a somite, cells can assume one of two final states: rostral (anterior part of a somite) or caudal (posterior part) [32]. Each somite is patterned along the anteroposterior direction, so that markers for the anterior and posterior parts of the somite form stripes of genetic expression [25, 33], Fig. 4A. Because of the coupling between embryonic growth and patterning, this phenomenon can be described in the following way from the cell standpoint: cells are first injected into the PSM in the very posterior part of the tail where they oscillate, then, as they get more anterior relative to the growing tail, the oscillation slows down and stops. Cells are then integrated into a future somite where they express either posterior or anterior fate markers, presumably based on the phase of the oscillation when it stops, Fig. 4A, middle. See [31] for more details and models of this process. Previous explorations of gene networks driving stripe formation coupled with embryonic growth have used in silico evolution of gene network models [34] or direct enumeration [35]. Remarkably, the best gene networks implicate a genetic oscillator upstream of a bistable system [34], suggesting that a genetic oscillator is the most parsimonious solution for this problem.

**Figure 4:**
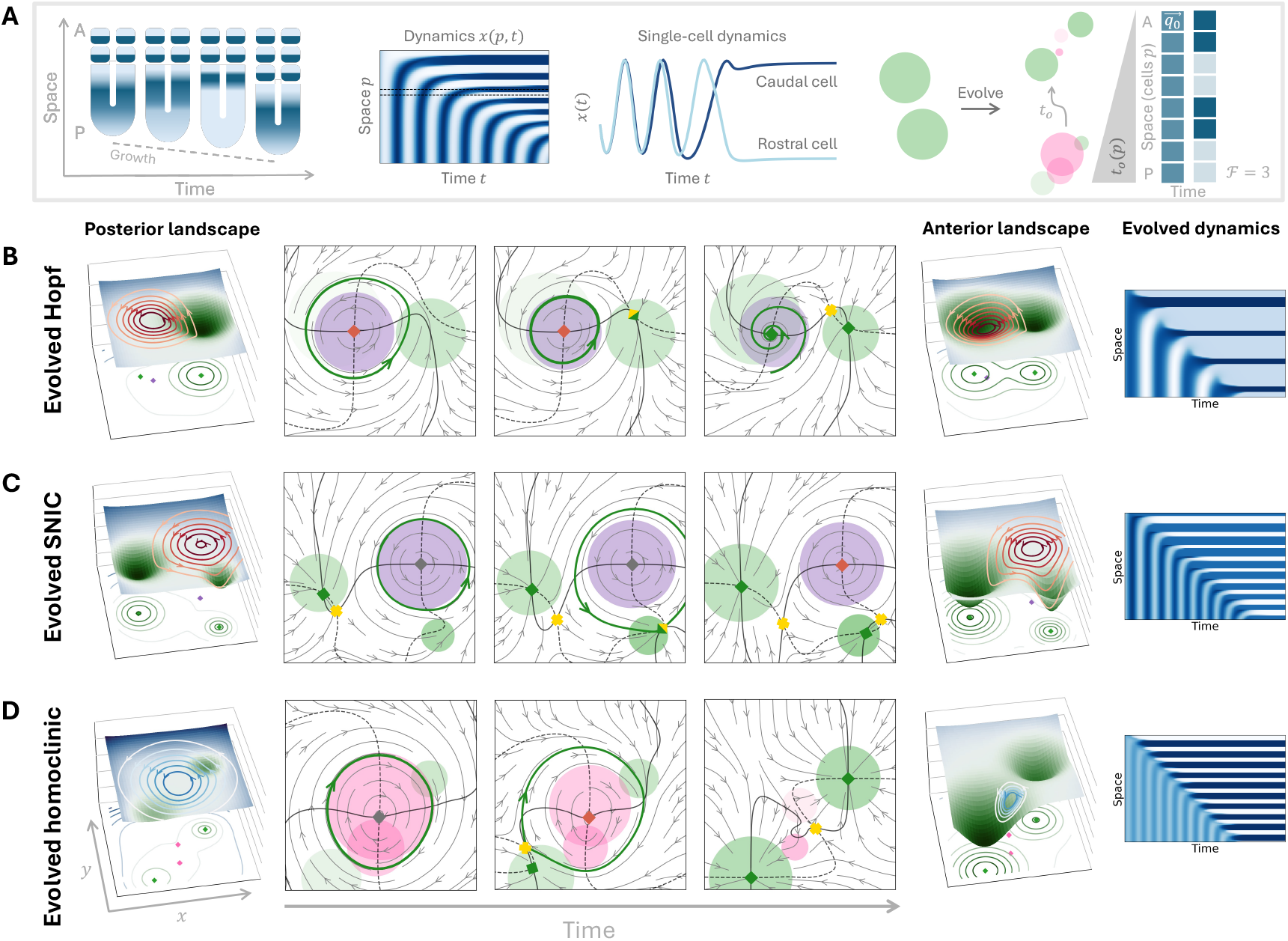
Evolving landscapes for vertebrate segmentation. **A)** Phenomenology of somitogenesis and the optimization setup. During somitogenesis, new segments are formed from anterior (A) to posterior (P) while the tissue elongates. Gene expression dynamics in the tissue can be visualized in a kymograph, showing a transition from oscillations to a stable striped pattern. Dynamics of two representative cells (dashed horizontal lines in the kymograph) are shown as a function of time. Oscillations stabilize at discrete levels associated with cell fate (rostral/front of somite or caudal/back of somite). Based on these two fates, landscapes are initialized with two attractor modules and module addition is allowed. Two landscape configurations are parameterized: initial and final (modules shown with circles; radius ∼*σ*_*i*_ and opacity ∼ *a*_*i*_. A line of cells in physical space is simulated, where the landscape changes over time with an A-P timing gradient. In the phenotypic space, cells start at the same initial condition 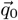. The final pattern is used to calculate the landscape’s fitness ℱ: the number of segment boundaries. For each of the evolved examples **(B-D)**, we show the flow at three chosen timepoints, as well as the gradient potential and the rotation contours for the first and last timepoint, and the kymograph produced for the line of cells (landscape variable *x* shown).

We use a setup similar to [34], but for landscape (instead of gene network) evolution. We simulate a line of cells in the PSM where each cell (position *p*) has the same landscape (genotype) and the same initial condition 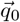. For this problem, the simulation is deterministic, with noise set to zero (*η* = 0). All module parameters *a*_*i*_, *σ*_*i*_ change sigmoidally at time *t*_0_(*p*) linear in *p*. This models an anteroposterior morphogen gradient combined with axis elongation [31, 32], and is similar to the 1D primary waves proposed in previous models [36, 31]. We define fitness as the number of formed segments – stripes with alternating fates in the final pattern [34]. For our minimalistic fitness function, we do not need to specify the exact location of the fate attractors, the width of the segments, or the proportion of rostral to caudal within a segment. The cells just need to end up in alternating “rostral” and “caudal” regions of the landscape to produce a segmented phenotype.

We initialize the landscape with two randomly generated attractor modules since these are necessary to produce the two fates. During evolution, we allow modules of any type to be added, and optimize the module locations 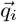 and two values for each parameter *a*_*i*_, *σ*_*i*_ - the initial and final configurations of the landscape. Details of the fitness function, hyperparameters and parameter priors used are provided in the Supplement.

We repeated the evolutionary procedure several times for a fixed number of generations and then analyzed the best evolved landscape in each population. Typically, after a few generations, a bistable system is established, producing one stripe. Subsequently, the landscape evolves a transition from an oscillator to a bistable system (Fig. 4 B-D) and a segmented phenotype. This qualitatively fits the results previously obtained with explicit gene network models where, similarly, a bistable system, then an upstream oscillator sequentially evolved, giving rise to a ‘Clock and Switch’ system [34, 31].

In evolved models of gene networks, the transition from the oscillating to stable states typically occurs through a Hopf bifurcation followed by a saddle-node bifurcation [37]. Remarkably, our modular approach gives rise to many more bifurcation scenarios, with three typical cases illustrated in Fig. 4 B-D (see also Supplementary movies 1-2). For each example, we show the landscape at an early developmental time, corresponding to posterior cells (P in Fig. 4A), transitioning towards the landscape at later developmental times, corresponding to anterior cells (A in Fig. 4A). The rightmost column shows the kymographs recapitulating the dynamics. While we recover the Hopf + saddle-node case (Fig. 4B), our simulated evolution discovers alternative transitions. For instance, Fig. 4C shows a transition occurring through a SNIC bifurcation, where the increasing weight of the bottom-right final attractor deforms the limit cycle until a saddle-node bifurcation occurs on the cycle. This solution is very similar to a previously proposed geometric model of vertebrate somitogenesis [27]. We also obtain a new scenario, Fig. 4D, via a homoclinic bifurcation based on the previous appearance of an attractor outside the cycle (bottom left attractor). This possibility was suggested in [31] and corresponds to a so-called switch-clock-switch mechanism.

All the above examples were obtained within the same number of evolutionary generations and under the influence of the same fitness function, but produce a different number of stripes and different bifurcations, corresponding to distinct evolutionary trajectories. Such variability suggests that specific landscapes might appear in response to different evolutionary constraints. Thus, we also ran simulations to evolve patterns with 5, 10 and 15 segment boundaries, and compared their dynamics and landscapes (see Fig. S3 and technical details in Supplement). Patterns with fewer segments correspond to oscillations with longer periods, as expected from the Clock and Wavefront theory [31], and this is typically achieved through larger oscillation amplitudes (thus taking more time to complete a cycle) combined with SNIC bifurcations. Another interesting feature is observed in simulations with many segments: the proportion of cells reaching rostral/anterior (A) vs. caudal/posterior (P) fates tends to 1 (compared to simulations with fewer segments). This is intuitive since maximum fitness corresponds to a perfect spatial alternation of both states. Consistent with this observation, we find that for such landscapes the bifurcations creating the A/P fates happen at nearly the same value of the control parameter and are therefore temporally synchronized, Fig. S4. This suggests that specific selective pressures can lead to unexpected symmetries, reflected in both the final pattern and the bifurcation diagrams. We conclude that our landscape exploration model offers a rich dynamic repertoire, going beyond previous works focused on more classical gene regulatory network descriptions.

### Application to neuromesoderm differentiation reveals two distinct scenarios

#### Fitting landscapes to cell type proportions data

Our approach can also be applied to experimental data to infer plausible landscapes. We consider data on differentiation from stem cells to neural and mesodermal fates [14], Fig. 5. In their study, Sáez et al. exposed mouse embryonic stem cells to combinations of morphogens at different times to push the system toward distinct outcomes. Six cell states were then defined: epiblast (Epi), transitory (Tr), caudal epiblast (CE), anterior neural (AN), posterior neural (PN), and mesoderm (M), and the proportions of cells in each of those states were calculated based on a Gaussian mixture clustering algorithm. The authors proposed a landscape model for the data, where a binary choice landscape between AN, Epi and CE was ‘stitched’ to a heteroclinic flip landscape connecting CE to PN and M [14].

**Figure 5:**
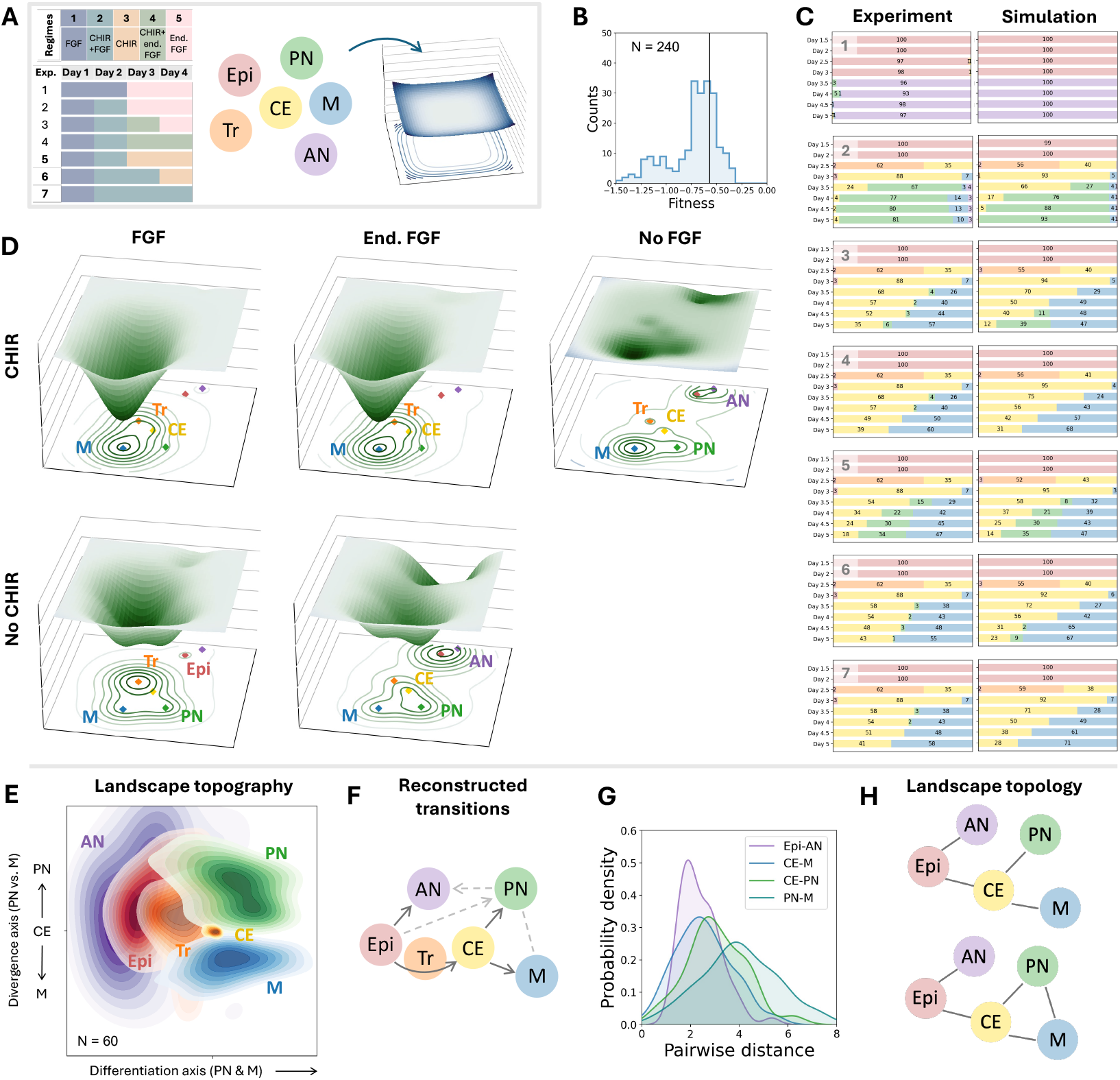
Evolving landscapes for neuromesoderm differentiation based on cell proportion data. **A)** Optimization setup. The training dataset consists of 7 experiments with 5 signal combinations (regimes). Initial training was done on experiments 5-7. Landscapes are initialized with a fixed number of attractor modules, one module per observed cell state. Module parameters are optimized for each signal combination. The fitness is based on the total KL divergence between the simulated and experimental cell state distributions over all timepoints. **B)** Achieved fitness distribution over evolutionary runs for experiments 1-7. The vertical line indicates the fitness threshold used to select successful simulations. **C)** Example of simulated cell state proportions in an optimized landscape, compared to data from 7 experiments. In each experiment, rows represent timepoints and colors indicate cell states, as defined in **(A). D)** Visualization of the evolved landscape example from **(C)**. The potential *ϕ* is shown for the five signalling regimes optimized. **E)** Probability distributions of optimized module locations 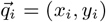 in the selected landscapes. The coordinate systems were aligned and rotated to account for equivariance. The *x, y* coordinates of CE module were fixed in the new coordinate system and a small random spread was added for visualization purposes. **F)** Cell differentiation tree inferred from the simulations. Dark edges show transitions present in all landscapes, light edges are transitions seen in some landscapes. Arrows indicate the direction of transitions. **G)** Probability distributions for pairwise distances between modules. **H)** Two landscape topologies obtained from the ensemble of evolved landscapes. Graphs show two binary decisions - first at Epi and second at CE. Edges indicate realizable saddle connections between attractors.

We wondered if and what different landscapes were possible, and, more generally, how data on cell proportions constrains a possible landscape. We thus specified six attractors, corresponding to the six cell states, and let our evolutionary algorithm optimize their positions and shapes to match the data (see fitness definitions in the Supplement, Eqs. S49-S50. Morphogen inputs were assumed to change the attractor parameters (strength and size), as illustrated in Fig. 2. Specifically, these parameters were modelled to be piecewise constant in simulation time, depending on the morphogen combination currently applied (Fig. 5A), which allows for nonlinear effects. Cell trajectories were simulated in the dynamic landscape, with noise magnitude fixed as a hyperparameter.

The full landscape parameterization in this setting comprises 72 parameters; to reduce the number of parameters optimized at once and speed up fitness evaluations, we performed an iterative evolutionary optimization (see Supplement for procedure and parameter values). We first noted that out of five morphogen combinations applied, three regimes were the most distinct (FGF, CHIR, and CHIR+FGF), and two more were obtained by replacing the FGF signal with endogenous FGF. We therefore started by optimizing for a subset of three experiments describing the posterior neural/mesoderm decision, with five cell states and three morphogen regimes (N = 1200 runs). We then performed a full optimization for seven experiments, with most parameters fixed from the first step (N = 240 runs). Each run was performed for a population of *P* = 200 landscapes and 300 generations. After 100 generations, a fitness plateau was typically observed, although with a variable plateau height. Fig. 5B illustrates the fitness distribution obtained for 240 runs. We then selected the best 25% (N=60) of landscapes for further analysis. Visually, those simulations present cell state proportions in excellent agreement with experiments (see Fig. 5C-D for an example). We provide in Supplementary Data 1 a visual study of each of those best 60 landscapes, including the proportion of cell states as a function of time and a description of the landscape corresponding to the final decision (see results below). Typical examples of cellular dynamics recapitulating the experiments are provided in Supplementary movies 3-6.

#### Landscape optimization discovers a constrained landscape topography

Next, we analyzed the ensemble of obtained landscapes, starting with the topography of the attractors considered. Since we did not constrain the attractors’ locations, direct landscape comparison is difficult. Furthermore, landscapes are subject to translational, rotational, and reflectional invariance. We therefore perform a simple standardization of the coordinate system (see Supplement): landscapes are aligned by the location of the caudal epiblast (CE) module, and the epiblast (Epi), posterior neural (PN) and mesoderm (M) modules are used to define the *x*- and *y*-axes. Based on these definitions – and further confirmed by the geometries observed in this coordinate system – we interpret the *x*-axis as the PN-M differentiation axis and the *y*-axis as the PN/M divergence axis. Remarkably, after such alignment, we obtain very well-defined localizations of the modules, shown in Fig. 5E. In particular, the Epi, Tr, and CE states lie approximately on a line, which is further aligned with the typical PN-M differentiation direction, effectively recapitulating a developmental time axis. Such an arrangement is not a necessity, and we see rare exceptions where trajectories can go backwards in the 2D plane, closer to the initial Epi attractor (see Fig. S5; of note, such ‘backward’ motions towards an early attractor have been observed in Waddington-like machine learning dynamics [38]). Nevertheless, our simulations suggest that simple knowledge of the proportion of cell states as a function of time statistically constrains the landscape topography (see Fig. S6 for the emergence of the constrained landscape topography as a function of fitness threshold and data provided).

At the cell trajectory level, we recover the sequences Epi →AN and Epi →Tr →CE →PN & M. Fig. 5F represents typical cell differentiation trajectories, with solid lines indicating transitions observed in all simulations, and dashed lines indicating transitions only in some simulations. We notice that the transitions found by our optimization largely recapitulate previous descriptions with landscape structure assumed a priori [14], despite fitting only cell state proportions as a function of time. That said, we observe two types of rare transitions that were not previously described: in some simulations, we observe transitions from Epi to PN, or from PN to AN (Fig. S7). Fig. 5G summarizes the distribution of pairwise distances between modules, in particular, we find that pairwise distances are more constrained for modules connected by a differentiation route (CE to M, CE to PN, Epi to AN) than for those that are not (PN to M).

From our simulations, we also obtain a set of interpretable, high-level parameters characterizing modules. In particular, we can compute their size and depth in the potential space. We can then compare the distributions of parameters for different modules in each of the signalling regimes (Fig. S8). For instance, overall, we see that module sizes in CHIR+FGF/CHIR+endogenous FGF signalling are typically inversely correlated with module sizes in the absence of either morphogen, suggesting that morphogens act in a synergistic way to change the landscape.

#### Two solutions for neuro-mesoderm proportioning

Next, we considered the topology of our best 60 landscapes, and manually classified decisions based on the nomenclature developed in [12] (see Supplement and Supplementary Data 1). For this, we mapped the optimized landscape configurations to regions of the bifurcation space, with the method illustrated in Fig. S9 and results summarized in Fig. 5H. For the initial decision between Epi, AN and CE, we always obtain a binary choice (double cusp landscape with all-or-nothing dynamics), matching the solution proposed in [14] (Fig. S10). Meanwhile, for the final decision between CE, PN and M, we typically obtained two possible landscapes. In Fig. 6 we show four examples of evolved landscapes, two examples per topology. The first possibility corresponds to a double cusp landscape, with one saddle between CE and PN and the other between CE and M. The second is a heteroclinic flip landscape, similar to [14]. This topology contains a saddle between PM and M, as we illustrate in the ‘Decision structure’ graphs in Fig. 6A. In these graphs, each edge indicates a saddle connection realizable for some parameter values. Although at a given time both topologies contain up to two saddles, the heteroclinic flip decision structure forms a closed triangle because the connection from CE can flip (CE to PN or CE to M).

**Figure 6:**
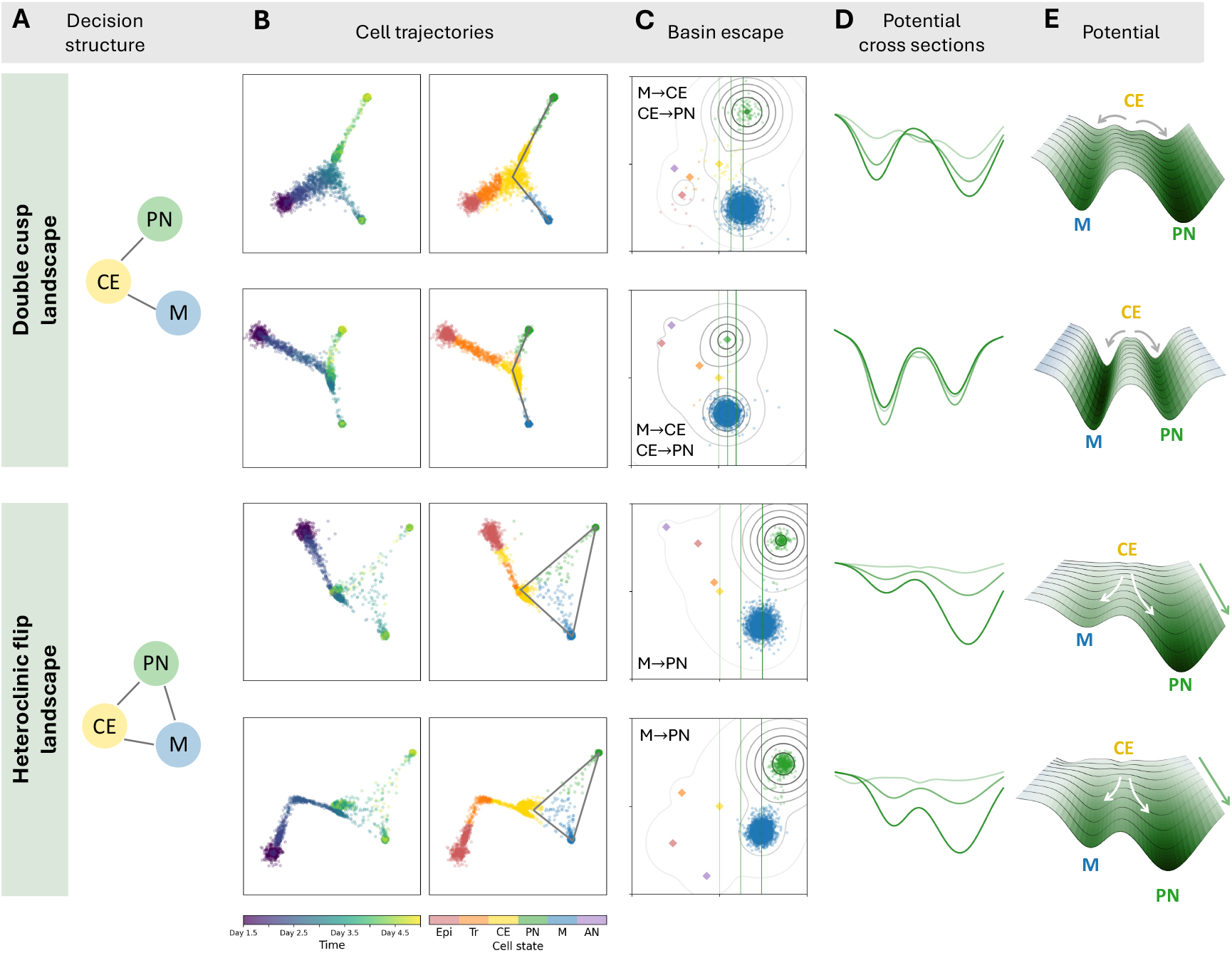
Two solutions for the posterior neural/mesoderm decision. **A)** Decision structures are shown as graphs where edges indicate a realizable saddle connection. For columns **(B-E)**, two evolved examples are shown for each topology. **B)** Cell trajectories starting from the initial Epi state (simulation for experiment 5). **C)** Basin escape numerical experiments used to confirm the decision structure; shown here is an M state initialization under CHIR signalling. Scatter plots of cell trajectories are overlaid on grey potential contours. Diamonds indicate module locations. **D)** Potential in CHIR signalling at cross sections marked by vertical green lines in panel **(C). E)** The region of landscape between the first and last vertical lines in **(C)** is shown as a potential surface plot.

Both landscapes appear consistent with the data: we saw no clear differences between these scenarios in the quality of the fit of proportions (See Supplementary Data 1). In most optimized landscapes of both topologies, we also notice that the CE attractor is very shallow (see Fig. 6), which matches the model proposed in [14]. The difference between the two landscapes is more apparent from simulated cellular trajectories in 2D. First, for the double cusp landscape, cells typically stay canalized along the manifolds connecting CE to either PN or M, while in the heteroclinic flip landscape, trajectories are typically more diffuse within the triangle formed by the three attractors, Fig. 6B. Second, the decision structure can be reconstructed from cellular trajectories, providing a criterion for distinguishing between the two solutions. For this, we ran simulations with increased noise to detect escapes from the fate attractors PN and M (Fig. 6C). We find a direct PN-M transition in the heteroclinic flip landscape, which is absent in the double cusp landscape. In the latter topology, the only route from PN to M passes through CE (dedifferentiation). Notably, realizing such a path requires a much higher noise magnitude than used for the experiment simulations, to enable escape from a committed fate attractor.

To further illustrate the origin of the difference between those scenarios, we plotted cross-sections of the potential as the cell moves towards differentiation (lines along the differentiation axis). We see clear topographical differences: for the double cusp landscape, the local minima of CE, M and PN align along the transverse direction (divergence axis). This explains why the trajectories are typically canalized into well-separated valleys. Conversely, for the heteroclinic flip, the CE attractor is uphill of the ridge (saddle) separating M and PN, with a flat intermediate region and a relatively smaller distance between the fate attractors. We also notice that, compared to the double cusp landscape, the optimized heteroclinic flip has a more ‘downhill’ structure (compare relative levels of cross-sections in Fig. 6D), thus realizing the classical Waddington picture of a binary cell fate decision (Fig. 6E, see also pictures of 3D printed models in Fig. S11).

#### Building up constraints in the landscape

To ensure the generality of our results and further illustrate how specific assumptions can be implemented in our framework, we also ran simulations with additional hypotheses: a *temporal regularization* scheme and a *linear combination* scheme. For temporal regularization, we only keep attractors observed in a given signalling regime, and set the parameters of other attractors to 0. At each timepoint, this reduces the landscape to its essential components, and the lower number of parameters allows to streamline the optimization procedure. In particular, we can skip optimizing for the subset of three experiments and proceed directly with seven experiments. For the linear combination landscape, the effects of two morphogens are combined linearly into the strength and size of the attractors (see technical details and implementations in the SI). Typical results of these regularized simulations are shown in Figs. S12 and S13 and are consistent with the more general results described above, with the caveat that the optimization of linear landscapes had a lower convergence rate. With both approaches, we recover the two landscape topologies (double cusp and heteroclinic flip). The linear assumption allows for continuous modulation of the flow to explicitly observe bifurcations at intermediate signal levels and further connect the evolved landscapes to the corresponding catastrophe diagrams, Fig. S9.

## Discussion

We propose a framework for constructing landscape descriptions of cellular differentiation trajectories. Our formalism derives from Waddington’s intuition of cell states as valleys, which we explicitly construct through the modular definition of local attractors, with additional modules to account for local repellers and rotational components. Such modular interactions are in line with the idea that cellular attractors are biologically defined by co-expressed genes interacting with specific enhancers, similar to previous systems biology works [27, 39]. We show that we can easily engineer well-known landscapes, e.g., for binary decisions or more complex limit cycle bifurcations. Critical points such as saddles emerge from the interactions of modules, and bifurcations typically occur when modulating relative landscape weights, which thus act as effective control parameters.

We applied this formalism to systematically derive families of landscapes using an evolutionary algorithm to mimic biology. We first illustrated our approach with the problem of segmentation coupled to growth. We observed the spontaneous evolution of a segmentation clock, combined with limit cycle bifurcations. While our landscape model recovers a clock and switch system previously obtained in a gene-network based formalism, it also appears more versatile, with a broader variety of scenarios spontaneously evolving (SNIC, homoclinic bifurcations). Our framework thus allows us to explore if multiple solutions - multiple fitness maxima - are possible and evolvable. Evolutionary optimization of landscapes could also help to study the sequence of mutations changing the landscapes: identifying which landscape mutations provide the largest gain in fitness and looking for recurrent patterns of mutations (see examples for explicit models of gene networks in [34, 40]). This could further inspire insights into the evolution or plasticity of complex mechanisms, either evolved independently or widely diverged, such as segmentation controlled by oscillators in metazoans [41, 42]. As an example, we observe here that evolutionary pressure for more segments leads to more symmetries in the landscape and in the bifurcation diagram (Fig. S4), hinting at a possible scenario for the evolution of symmetries in biology [43].

Next, we used our approach to fit experimental data, based on the cell state proportions at different developmental times and under various morphogenetic controls. For this problem, the only constraints are the number of attractor states, while all other parameters (locations, depths, etc.) remain free. Strikingly, comparing many successful simulations, we saw rather constrained *topographies* (i.e., relative location of attractors), but variable *topologies* (i.e., attractor connections), self-consistent with and justifying our formalism based on the definition of modules corresponding with cellular states.

We obtained two types of landscapes, distinguished by the bifurcations involved in the final neuro-mesoderm proportioning decision. For this system, one type of landscape has already been proposed, combining two binary decisions: a binary choice and a heteroclinic flip [14]. The other solution we found relies on two binary choices, with differential attractor depths depending on the signalling condition. These findings are robust to specific assumptions about the landscape (i.e., unconstrained nonlinear response, minimal components at each timepoint, or linear signal dependency). While our results show that there are many quantitatively different ways (i.e., parameter sets) to obtain similar dynamics and fate outcomes, the statistically constrained topographies and the emergence of only two topologies indicate that the dataset we fit – proportions of cell states as a function of time – constrains high-level properties of the underlying landscapes. These findings confirm that predictive and actionable geometric models can be built from datasets of relatively modest size as long as they are well designed [44, 11, 13, 45].

Having identified families of landscapes with distinct topologies, we also showed that distinguishing between solutions requires a more detailed analysis of time-resolved cellular trajectories, which are more canalized in the binary choice scenario. Such trajectory information could be incorporated in the form of intermediate states and/or transitions between states (vs. the proportion of states). Transitions between states could be inferred, for example, through pre-identification of plausible routes [16] or computation of Wasserstein distance [46], and used to bias the topography of attractors. This could further facilitate optimization for more complex problems with a large number of cellular states.

Looking ahead, our framework could naturally be extended to account for cell-cell interactions for spatial problems. The position of a given cell in its landscape could either modulate the landscape parameters of a neighboring cell [17], or the positions of cells could be coupled within a shared landscape. On the computational side, our framework could be modified to include differential programming for automatic gradient descent in parameter space [47] once the number of modules is fixed.

It is now standard in developmental biology to draw 2D embeddings of differentiation dynamics in latent space and to interpret the resulting trajectories. Our goal here is to infer explicit geometries amenable to theoretical studies [12]. Such landscape descriptions solve well-known issues resulting from insufficiently constrained representations (inferred flows leading to nowhere, jumps in latent space… [48]). We restrict our landscape model to a 2D space, but it is unclear if such dimensional reduction is always possible. More complex problems may require a tailored approach to obtain 2D projections, or even moving to higher dimensions, where analogous local modules could be defined (e.g., 3D Gaussian attractors, see also a recent 5D treatment of the neuromesoderm differentiation problem in [49]). An alternative and complementary approach could be to use Gaussian modules to model probability distributions, taking the log probability as potential, akin to what is done in diffusion/flow matching models [50], but such formalisms do not explicitly account for biological features like well-defined attractors or non-gradient dynamics such as oscillations. Lastly, high-dimensional gene expression states could be recovered from the phenotypic coordinates and predicted by training an explicit decoder from the 2D latent space in which we simulate the dynamics, as done in [51, 52].

## Methods

For landscape simulations, stochastic cellular trajectories were integrated using the Euler-Maruyama method with a fixed noise magnitude. Cell states were assigned from cell coordinates using a Gaussian mixture model. Fitness evaluations during evolutionary optimization were parallelized with Python’s multiprocessing. Landscape topography statistics were analyzed using kernel density estimation in Seaborn.

Details on general model construction, optimization, and parameters are provided in the Supplement. The codes used for this work are available in the GitHub repository https://github.com/victoria-mochulska/evoscape.

## Aknowledgement

We thank Eric Siggia, Madhav Mani, and members of the François group for useful discussions. This research is supported by an NSERC Discovery Grant, a CIHR Program Grant and a Fonds Courtois Grant to PF, and a doctoral research grant from the Fonds de recherche du Québec (FRQ) #328292 to VM.

## Notes

https://github.com/victoria-mochulska/evoscape

### Competing Interest Statement

The authors have declared no competing interest.

### Summary of Updates

Updated section Phenotypic landscape view of metazoan segmentation; Added section Building up constraints in the landscape; Updated SI: added sections S5.3 and S6.7; Added supplementary figures S3-S5, S9, S12, S13.

