## Supplementary Information for "Generative epigenetic landscapes map the topology and topography of cell fates"

September 19, 2025

### Supplementary Methods

#### S1 General parameterized landscape model

##### S1.1 Theoretical background

In the dynamical systems view, the gene expression of a cell follows the dynamics in an $n$-dimensional *gene space*, given by a smooth vector field $\vec{F}$:

$$\dot{\vec{Q}} = \vec{F}_\theta(\vec{Q}), \tag{S1}$$

where $\vec{Q}$ is the cell's coordinate in this space; $\vec{Q}, \vec{F}_\theta \in \mathbb{R}^n$, and $\theta$ indicates a parameter dependency. Each cell state is an attractor, or stable fixed point, of the dynamics: $\vec{F}_\theta(\vec{Q}) = 0$, and cellular decisions correspond to transitions between different basins of attraction.

Without knowledge of $\vec{F}_\theta$, the first assumption, proposed in [1], is to consider the class of *Morse-Smale systems*. These are among the simplest and most common dynamical systems [2] and are suitable for describing biological dynamics due to their structural stability. *Structural stability* means that the qualitative form of the system remains unchanged under small perturbations of parameters [3], making it a natural assumption for robust developmental processes. Changes in qualitative behavior can then occur only at the bifurcation set in the parameter space, with codimension-1 bifurcations usually considered.

Morse-Smale systems are defined by several conditions [4, 5]. Here, we only outline the practical consequences of these conditions for modelling. First, Morse-Smale systems can contain attractors, repellers, saddles, stable and unstable limit cycles, but no neutrally stable states (with zero eigenvalues) or lines of fixed points. Second, the unstable manifold of one saddle cannot coincide with the stable manifold of another. Thus, such dynamical systems have a simple structure: attractors are connected through the unstable manifolds of saddle points, and a trajectory escaping a basin of attraction will follow the unstable manifold of the saddle towards a different attractor [6].

The second simplification involves dimensionality reduction, initially motivated by the notions of phenotype, canalization, and geometric descriptions [7–9]. A more formal mathematical result exists for a subclass without limit cycles and featuring binary decisions, in which one attractor is directly connected through saddles to two other attractors. In this case, Rand et al. show that the topology of the routes between attractors is dimension-independent and can be embedded in two dimensions, with bifurcations of these routes also represented in 2D [1]. This statement allows to consider cell fate decisions as locally two-dimensional and embed them in the *phenotypic space* $\vec{q}$, $\vec{q} \in \mathbb{R}^2$, with local mappings of the gene space $\vec{Q}$ to the phenotypic space $\vec{q}$. Still, we are interested in a broader class of biological systems, including non-gradient dynamics. We consider these in a 2D phenotypic space under a self-consistent assumption.

Therefore, for our landscape model, we consider a dynamical system in the phenotypic coordinate $\vec{q}$:

$$\dot{\vec{q}} = \vec{f}_\theta(\vec{q}), \quad \vec{q} = (x, y). \tag{S2}$$

##### S1.2 Details of modular construction

We represent the flow field $\vec{f}_\theta$ as

$$\dot{\vec{q}} = -\nabla\phi_0(\vec{q}) - \nabla\phi(\vec{q}) - \nabla^\perp\psi(\vec{q}). \tag{S3}$$

The use of the gradient (curl-free) and rotational (divergence-free) flows is motivated by the Helmholtz-Hodge decomposition [10, 11]. The field decomposed is assumed to be smooth and decaying for $q \to \infty$. The skew gradient operator $\nabla^\perp$ used for the rotation component is

$$\nabla^\perp = \begin{pmatrix} 0 & -1 \\ 1 & 0 \end{pmatrix} \nabla = (-\frac{\partial}{\partial y}, \frac{\partial}{\partial x}). \tag{S4}$$

We also introduce a confining term $\vec{f}_0 = -\nabla\phi_0(\vec{q})$ that is non-vanishing for $q \to \infty$. It is globally attractive and keeps the trajectories within a relevant region of the phenotypic space by preventing escapes to infinity. We use an order-4 confining potential:

$$\phi_0 = A_0 \frac{(x - x_0)^4 + (y - y_0)^4}{4}, \tag{S5}$$

where $A_0 \ll 1$ so that the dynamics in the relevant region is defined by $\phi$ and $\psi$. The confining term is usually centered at the origin: $(x_0, y_0) = (0, 0)$.

The scalar fields $\phi$, $\psi$ are constructed using modules – radial functions that represent local features of the landscape:

$$\phi = \sum_{i=1}^{n} \pm w_i(|\vec{q} - \vec{q}_i|), \quad \psi = \sum_{j=1}^{m} \pm w_j(|\vec{q} - \vec{q}_j|), \tag{S6}$$

where $n$ is the number of gradient modules, $m$ – number of rotational modules, $\vec{q}_i$, $\vec{q}_j$ are the locations of modules, and $w_i(q) > 0$, $w_i(q) \xrightarrow[q \to +\infty]{} 0$. The combination of two types of potential with two signs $(+1, -1)$ gives four types of modules.

Consider the gradient of one such module:

$$\nabla w_i(|\vec{q} - \vec{q}_i|) = w_i'(|\vec{q} - \vec{q}_i|) \frac{\vec{q} - \vec{q}_i}{|\vec{q} - \vec{q}_i|}, \tag{S7}$$

while for the rotation potential $\psi$,

$$\nabla^{\perp} w_j(|\vec{q}-\vec{q}_j|) = w_j'(|\vec{q}-\vec{q}_j|)\begin{pmatrix}0 & -1\\ 1 & 0\end{pmatrix}\frac{\vec{q}-\vec{q}_i}{|\vec{q}-\vec{q}_i|}. \tag{S8}$$

A module can thus be interpreted as linearized dynamics near a fixed point

$$\dot{\vec{q}} \propto \hat{M}\left(\vec{q}-\vec{q}_i\right), \tag{S9}$$

with a decaying nonlinear spatial weight $\pm\frac{w_i'(|\vec{q}-\vec{q}_i|)}{|\vec{q}-\vec{q}_i|}$.

The matrix $\hat{M}$ defines the type of module: attractor $\hat{M}_-^{\circledast}$, repeller $\hat{M}_+^{\circledast}$, left (clockwise) rotator $\hat{M}_-^{\circledcirc}$, or right (counterclockwise) rotator $\hat{M}_+^{\circledcirc}$, with

$$\hat{M}_-^{\circledast} = \begin{pmatrix}-1 & 0\\ 0 & -1\end{pmatrix} \qquad \hat{M}_+^{\circledast} = \begin{pmatrix}1 & 0\\ 0 & 1\end{pmatrix} \tag{S10}$$

$$\tag{S11}$$

$$\hat{M}_-^{\circledcirc} = \begin{pmatrix}0 & 1\\ -1 & 0\end{pmatrix} \qquad \hat{M}_+^{\circledcirc} = \begin{pmatrix}0 & -1\\ 1 & 0\end{pmatrix} \tag{S12}$$

We choose Gaussian weights $w_i$:

$$w_i(|\vec{q}-\vec{q}_i|) = a_i\sigma_i^2 \exp\left(-\frac{|\vec{q}-\vec{q}_i|^2}{2\sigma_i^2}\right), \tag{S13}$$

so that

$$\frac{w_i'(|\vec{q}-\vec{q}_i|)}{|\vec{q}-\vec{q}_i|} = -a_i \exp\left(-\frac{|\vec{q}-\vec{q}_i|^2}{2\sigma_i^2}\right). \tag{S14}$$

Thus, we express the dynamics as a sum:

$$\vec{f}(\vec{q}) = -A_0\begin{pmatrix}(x-x_0)^3\\ (y-y_0)^3\end{pmatrix} + \sum_{i=1}^{n}\exp\left(-\frac{|\vec{q}-\vec{q}_i|^2}{2\sigma_i^2}\right)\begin{pmatrix}\pm a_i & 0\\ 0 & \pm a_i\end{pmatrix}(\vec{q}-\vec{q}_i)+ \tag{S15}$$

$$+\sum_{j=1}^{m}\exp\left(-\frac{|\vec{q}-\vec{q}_j|^2}{2\sigma_j^2}\right)\begin{pmatrix}0 & \mp a_j\\ \pm a_j & 0\end{pmatrix}(\vec{q}-\vec{q}_j), \tag{S16}$$

where the first summation is over gradient modules and the second over rotational modules.

#### S1.3 Dynamic landscape: time or signal dependency

At a given timepoint or configuration, the constructed landscape is described by the set of parameters $\theta(t) = \{(\vec{q}_i, a_i(t), \sigma_i(t))\},\ i = \overline{1, N}$, where $N = n + m$ is the number of modules and $\vec{q}_i \in \mathbb{R}^2$. To be able to model cell state transitions, we introduce the dependency of the parameters on the external signal (morphogens) or time. First, we choose the **module locations** $\vec{q}_i$ to be **static** and independent of morphogens since these correspond to cell state identities - particular locations in gene expression space. We do not expect these identities to change drastically with external signals, rather, a given state can become (un)populated with cells over time or change its availability due to bifurcation. The bifurcations, in turn, are controlled by shape parameters $a_i$ and $\sigma_i$. The width of the Gaussian $\sigma_i$ is most directly related to the size of the basin of attraction, while changing $a_i$ allows a module to effectively appear or disappear (delta-like shapes with small $\sigma$ and high $a$ are undesired, so only $a_i$ is allowed to reach 0). The **shape parameters** $a_i$ and $\sigma_i$ are therefore **dynamic**.

Concretely, we model the dynamic parameters as functions of simulation time (and implicitly the signal/morphogen):

$$a_i(t) = g(t, \vec{a}_i), \quad \sigma_i(t) = g(t, \vec{\sigma}_i), \tag{S17}$$

where $\vec{a}_i, \vec{\sigma}_i \in \mathbb{R}^K$ are parameter vectors. The full parameterization of the landscape with $N$ modules becomes $\Theta = \{(\vec{q}_i, \vec{a}_i, \vec{\sigma}_i)\}$, $i = \overline{1, N}$. The dimensionality $K$ and the form of $g$ are chosen based on the biological context and signalling dynamics. In particular, consider the following cases:

#### Static landscape

If there are no changes in the landscape, $g \equiv const$, $K = 1$, so that $a_i(t) = a_i^0$ and $\sigma_i(t) = \sigma_i^0$.

#### Sigmoidal landscape

For a landscape that smoothly changes from one configuration (initial) to another (final), $g$ is sigmoidal in time and $K = 2$. The parameter vectors describe the initial and final values, the transition time $t_0$ is the center of the sigmoid, and a timescale parameter $\tau$ determines the width of the sigmoid:

$$a_i(t) = a_i^1 + \frac{a_i^2 - a_i^1}{2}\left(1 + \tanh\frac{t - t_0}{2\tau}\right), \tag{S18}$$

and analogously for $\sigma_i(t)$.

#### Piecewise landscape

If there are $K$ configurations of the landscape, and the changes are assumed to be much faster than the duration between transition times, then $g$ is a piecewise constant function:

$$a_i(t) = \begin{cases} a_i^1, & t < t_1 \\ a_i^2, & t_1 \le t < t_2 \\ \vdots & \\ a_i^K, & t \ge t_{K-1} \end{cases} \tag{S19}$$

The transition times $\vec{t_0} = (t_1, t_2, \ldots, t_K)$ are fixed by the experiment or problem setup.

##### S1.4 Numerical implementation

To simulate trajectories in the landscape, we use the Euler-Maruyama method for numerical SDEs. The integration step is implemented as follows:

$$\begin{aligned}
\text{Evaluate parameters:} \quad & a_i(t) = g(t, \vec{a}_i), \quad \sigma_i(t) = g(t, \vec{\sigma}_i) \quad \text{for } i \in \overline{1, N}; \\
\text{Evaluate derivatives:} \quad & \dot{\vec{q}}(t) = \vec{f}\left(\vec{q}(t),\, \{\vec{q}_i, a_i(t), \sigma_i(t)\}\right); \\
\text{Update coordinates:} \quad & \vec{q}(t+\delta t) = \vec{q}(t) + \dot{\vec{q}}(t)\,\delta t + \eta \mathcal{N}(0,1)\sqrt{\delta t}; \\
\text{Update time:} \quad & t \to t + \delta t.
\end{aligned} \tag{S20}$$

Here, $N$ is the number of modules in the landscape, $\eta$ is the noise magnitude, and $\mathcal{N}(0,1)$ denotes a standard normal random variable. The integration step was chosen such that $\Delta T/\delta t = n_{\delta t}$, where $\Delta T$ is the data or observation timestep, and $n_{\delta t} \in [50, 200]$.

The scripts used for simulations and optimization, as well as notebooks for visualizations and analysis, are available at `https://github.com/victoria-mochulska/evoscape`.

#### S2 Constructed bifurcations

Here, we study some simple module combinations analytically to illustrate the landscape's dynamical repertoire. We show that the proposed landscape construction results in Morse-Smale systems and that the relevant bifurcations can be obtained, involving either $a_i$ or $\sigma_i$ as control parameters.

##### S2.1 Two attractor modules

Consider two attractor modules far away from any other modules and without the influence of the global potential, $A_0 << 1$. The coordinate system is chosen so that $\vec{q}_1 = (-l, 0)$ and $\vec{q}_2 = (l, 0)$. We have the equation

$$\begin{pmatrix} \dot{x} \\ \dot{y} \end{pmatrix} = \exp\left(-\frac{(x+l)^2 + y^2}{2\sigma_1^2}\right)\begin{pmatrix} -a_1(x+l) \\ -a_1 y \end{pmatrix} + \exp\left(-\frac{(x-l)^2 + y^2}{2\sigma_2^2}\right)\begin{pmatrix} -a_2(x-l) \\ -a_2 y \end{pmatrix} \tag{S21}$$

The two modules can overlap to create one stable fixed point or, if they are far apart, they can form two stable fixed points and a saddle between them. To find the location of the fixed points, first assume the weight of the left module at a fixed point is $p$ times the weight of the right module, $0 < p < +\infty$:

$$e^{\left(-\frac{(x+l)^2+y^2}{2\sigma_1^2}\right)} = p e^{\left(-\frac{(x-l)^2+y^2}{2\sigma_2^2}\right)}. \tag{S22}$$

This gives the system of equations:

$$p a_1(x+l) + a_2(x-l) = 0; \tag{S23}$$

$$p a_1 y + a_2 y = 0, \tag{S24}$$

with the solution $x^* = \frac{a_2 - p a_1}{a_2 + p a_1} l$, $y^* = 0$. Substituting these coordinates into Eq. S22 and denoting $\alpha = a_2/a_1$, $\kappa = \sigma_2/\sigma_1$, gives the self-consistency relation:

$$\ln p = \frac{2l^2}{\sigma_2^2}\frac{p^2 - \alpha^2\kappa^2}{(p+\alpha)^2} = F(p), \tag{S25}$$

which can have one or three solutions $p$ depending on the values of $\alpha$, $\kappa$ and $l$. For the graphical solution $\ln p = F(p)$, it is useful to note that $F(p) \to \text{const}$ as $p \to +\infty$. In the three solutions case, we can approximate two solutions as $p = 0$ (stable fixed point at the right module $\vec{q}_2$) and $p = +\infty$ (stable fixed point at the left module $\vec{q}_1$).

As expected, changing the width and/or strength of a module is a simple way to obtain saddle-node bifurcations in the landscape.

#### S2.2 Hopf bifurcation

To work with modules in polar coordinates $\vec{r} = (r, \theta)$, we can use

$$\dot{r} = \frac{x\dot{x} + y\dot{y}}{r}, \quad \dot{\theta} = \frac{x\dot{y} - y\dot{x}}{r^2}. \tag{S26}$$

Then, the dynamics of a gradient module located at the origin is simply

$$\dot{r} = \pm a_i r \exp\left(-\frac{r^2}{2\sigma_i^2}\right) \tag{S27}$$

$$\dot{\theta} = 0 \tag{S28}$$

while for a rotational module

$$\dot{r} = 0 \tag{S29}$$

$$\dot{\theta} = \pm a_i \exp\left(-\frac{r^2}{2\sigma_i^2}\right) \tag{S30}$$

Consider now the normal form of the Hopf bifurcation in polar coordinates:

$$\dot{r} = (\mu - r^2)r \tag{S31}$$

$$\dot{\theta} = \omega \tag{S32}$$

where a limit cycle exists for $\mu > 0$, with the radius $R = \sqrt{\mu}$.

A minimal limit cycle oscillator landscape is constructed with two gradient modules and a third rotational module. The rotational module provides the angular dynamics, with $a_3 = \omega$ and $\sigma_3 = \sigma_\omega \gg 0$:

$$\dot{\theta} = \omega \exp\left(-\frac{r^2}{2\sigma_\omega^2}\right) \tag{S33}$$

For the radial part, we need a 'Mexican hat' landscape, which can be created by combining an attractor module with a repeller module. In the simplest case, they are aligned at $x = 0,\ y = 0$:

$$\dot{r} = -a_1 r \exp\left(-\frac{r^2}{2\sigma_1^2}\right) + a_2 r \exp\left(-\frac{r^2}{2\sigma_2^2}\right) = \dot{r}_c(r) \tag{S34}$$

Around $r=0$, we approximate the weights $\exp \varepsilon \approx 1+\varepsilon$ to get

$$\dot{r} = \left( (a_2 - a_1) - \left( \frac{a_2}{2\sigma_2^2} - \frac{a_1}{2\sigma_1^2} \right) r^2 \right) r, \tag{S35}$$

which can be rescaled to get the Hopf bifurcation normal form with

$$R = \sigma_2 \sqrt{\frac{2(a_2 - a_1)}{a_2 - \frac{\sigma_2^2}{\sigma_1^2} a_1}}. \tag{S36}$$

For a supercritical Hopf bifurcation, we need $\sigma_2 < \sigma_1$, so the limit cycle exists for $a_2 > a_1$, when the center of the landscape is destabilized.

#### S2.3 SNIC bifurcation

To obtain a saddle-node in cycle (SNIC) bifurcation, we add a small attractor module close to the limit cycle in the oscillator landscape given by Eqs. S33-S34. Suppose for concreteness that the oscillator landscape has a limit cycle at $r = R = 1$, the new module is located at $\vec{q}_4 = (x_0, 0) = (1, 0)$, and $\sigma_4 < 1$. Then, using Eqs. S26, we have:

$$\dot{r} = \dot{r}_c(r) + a_4 \left( x_0 \cos\theta - r \right) \cdot \exp\left( -\frac{|\vec{r} - \vec{r}_4|^2}{2\sigma_4^2} \right) \tag{S37}$$

$$\dot{\theta} = \omega - a_4 \frac{x_0 \sin\theta}{r} \exp\left( -\frac{|\vec{r} - \vec{r}_4|^2}{2\sigma_4^2} \right) \tag{S38}$$

Here, $|\vec{r} - \vec{r}_4|^2 = r^2 + x_0^2 - 2x_0 r \cos\theta$, and we take the size of the rotational module $\sigma_\omega \gg 1$.

Consider the saddle-node point at the bifurcation $(r^*, \theta^*)$. The new saddle-node pair appears initially at this point, and then the two fixed points move further apart after the bifurcation. The added attractor module will slightly distort the shape of the limit cycle from the unit circle. Assuming that this distortion is small, we approximate the location of the saddle-node point as $(r^*, \theta^*) \approx (1, \theta^*)$, with $\theta^*$ small (close to the location of the new module). Then, with $r = 1$, $x_0 = 1$, we have for the angular dynamics:

$$\dot{\theta} = \omega - a_4 \sin\theta \exp\left( -\frac{1 - \cos\theta}{\sigma_4^2} \right) \approx \omega - a_4 \theta \left( 1 - \frac{\theta^2}{2\sigma_4^2} \right). \tag{S39}$$

Finding the minimum of RHS, we have

$$\theta^* = \sqrt{\frac{2}{3}} \sigma_4, \tag{S40}$$

which finally leads to the normal form of a saddle-node bifurcation for the angular variable:

$$\dot{\tilde{\theta}} = a_4 \left( \left( \frac{\omega}{a_4 \theta^*} - \frac{2}{3} \right) + \tilde{\theta}^2 \right) = a_4 \left( \mu + \tilde{\theta}^2 \right), \tag{S41}$$

where $\tilde{\theta} = \frac{\theta - \theta^*}{\theta^*}$. With a fixed $\sigma_4$, a saddle-node pair in the cycle is created for $a_4 > \frac{3\omega}{2\theta^*}$. Thus, we see that the SNIC bifurcation happens when the strength of the added attractor module cancels the rotational flow around the cycle.

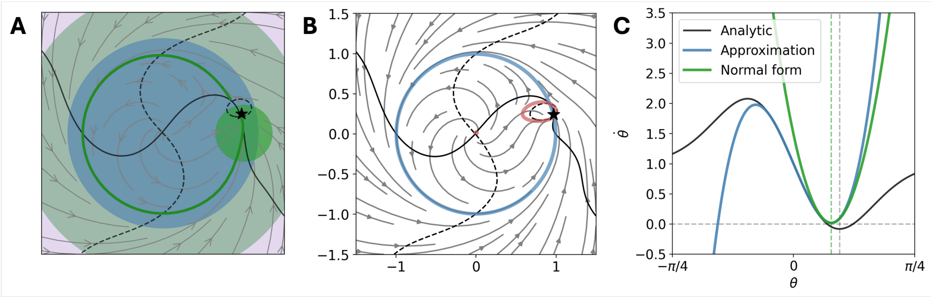


Figure S1: **SNIC bifurcation. A)** Minimal construction with modules: an oscillator landscape with an added small attractor module (bright green). Two fixed points appear on the cycle (nullcline crossings). The star indicates the location of the analytically calculated saddle-node point $(r^*, \theta^*)$. **B)** Approximating the limit cycle with a circle (no radial distortion). The red line is the $\theta$-nullcline. **C)** Angular dynamics close to the bifurcation $\dot{\theta}(r = 1, \theta)$. The original analytic expression Eq. S38, the approximation Eq. S39, and the derived normal form Eq. S41 are shown.

### S3 Cell state assignment

The state of a given cell is described by its landscape (phenotypic) coordinate $\vec{q}(t)$. For many applications of the landscape model, it is useful to discretize cell states (i.e, for comparisons with clustered experimental data or visualizations). For this, we supplement our modular construction with a Gaussian mixture model (GMM) to assign each cell to one of the attractor modules. For each cell, we calculate probabilities $p_i$ for every module $i = \overline{1, N}$:

$$p_i(\vec{q}, t) = \frac{1}{\sigma_i(t)^2} \exp\left(-\frac{|\vec{q} - \vec{q}_i|^2}{2\sigma_i(t)^2}\right) \tag{S42}$$

and assign the state based on the maximum probability:

$$s(\vec{q}, t) = \arg\max_i \; p_i(\vec{q}, t) \tag{S43}$$

Note that the state at coordinate $\vec{q}$ is time-dependent, which is expected since the basins of attraction of different modules can shrink or expand. Our procedure approximates the basins of attraction, but, contrary to a strictly basin-based assignment, allows to have cells assigned to transitory states [12]. In this case, a module does not create an attractor fixed point, but rather a transient region of a landscape (with potentially slower flow) is assigned to a state.

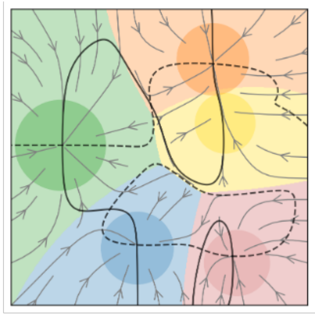

Figure S2: Example of cell state assignment: colored regions indicate coordinates assigned to each of the modules (circles). The yellow state here is an example of a transitory (non-attractor) state, as the yellow module does not create a fixed point.

### S4 Evolutionary optimization algorithm

For landscape optimization, we use computational evolution [13, 14]. This algorithm simulates natural selection based on a fitness function. Firstly, a fitness function is constructed for a specific biological system. This is essentially a task that we intend the landscape to perform, and it can be based on phenomenology or experimental data from the biological system. The fitness evaluation involves integrating trajectories in the landscape and can use readouts such as cell fate, cell state proportions, and temporal patterns.

The optimization is started with a set (population) of $P$ randomly initialized landscapes, with population size $P$ chosen to be even. In each iteration, half of the landscapes are randomly mutated using one of the mutation operators. The probabilities of applying each mutation are set as hyperparameters. After fitness evaluation, the less fit half of the landscapes are discarded, and the other half is duplicated, proceeding to the next iteration.

As mutation operators, we use addition, deletion, and modification, applied with relative probabilities $p_{add}$, $p_{drop}$, and $p_{mod}$ respectively, $p_{add} + p_{drop} + p_{mod} = 1$. The probability values used in applications are provided in Tables S1 and S4. The mutation operators are implemented as follows:

- addition: a new module is generated, randomly choosing its type (potential or rotational), sign (attractor/repeller, clockwise/counterclockwise), and sampling its parameters $\vec{q}_i, \vec{a}_i, \vec{\sigma}_i$ from prior distributions. We use uniform prior distributions;
- deletion: one of the existing modules is chosen randomly and removed. This mutation is only available if there is currently more than one module;
- modification: one of the existing modules is chosen randomly, then one of its parameters $\vec{q}_i, \vec{a}_i, \vec{\sigma}_i$ is chosen randomly and resampled. For vector parameters, only one randomly chosen element is mutated. This mutation cannot modify the type and sign of the module.

The fitness function can also include penalties, i.e., to constrain the number of modules, ensure stability or the transient nature of some states, etc.

### S5 Modelling vertebrate segmentation dynamics

#### S5.1 Simulation

For this problem, we simulate a line of cells in physical space, where a propagating front (i.e., a moving morphogen gradient) drives changes of the epigenetic landscape. Each cell is characterized by its physical coordinate (position in the tissue) $p$, $p = 1, \ldots, n_{cells}$, and landscape coordinate (phenotype) $\vec{q}_p(t)$. The landscape is the same for all cells and changes in time sigmoidally, according to Eq. S18. The transition time $t_0$ is a function of cell position:

$$t_0(p) = \Delta t_0 * p \tag{S44}$$

so that $p = 1$ is the most anterior cell (earliest transition) and $p = n_{cells}$ is the most posterior cell (latest transition). All cells start with the same initial landscape coordinate $\vec{q}_0$ and move in the phenotypic space according to equation S3 for $0 < t < t_f$.

In this 2D landscape, we designate the $x$-coordinate as the output (it can be loosely associated with gene expression levels such as rostro-caudal polarity markers in somites). Cell state dynamics can be visualized as a kymograph: a heatmap of the output as a function of time and space $x_p(t)$. The resulting pattern is evaluated using the spatial profile at the last timepoint $x_p(t_f)$.

#### S5.2 Fitness function

The fitness function has a patterning term and a stability term:

$$\mathcal{F} = \mathcal{F}_{stripe} - \lambda \mathcal{F}_{stab}, \tag{S45}$$

where $\mathcal{F}_{stripe}$ is equal to the number of boundaries in the final pattern, defined using two thresholds $x_{low}$ and $x_{high}$:

$$\mathcal{F}_{stripe} = \sum_{p=1}^{n_{cells}-1} \left(x_p(t_f) < x_{low} \text{ and } x_{p+1}(t_f) > x_{high}\right) \text{ or } \left(x_p(t_f) > x_{high} \text{ and } x_{p+1}(t_f) < x_{low}\right) \tag{S46}$$

and the stability term penalizes changes in the pattern in the last $n_{st}$ timesteps:

$$\mathcal{F}_{stab} = \sum_{p=1}^{n_{cells}} \sum_{n=1}^{n_{st}} \left(x_p(t_f - n\Delta t) - x_p(t_f)\right)^2. \tag{S47}$$

The parameters used for segmentation landscapes and evolutionary optimization are given in the table below. The computation time for one optimization (all generations in one population) is around 2 minutes.

Table S1: Parameters used for segmentation landscape evolution

| Parameter | Notation | Value or prior |
|---|---|---|
| Simulation parameters | | |
| Number of cells | $n_{cells}$ | 50 |
| Front propagation | $\Delta t_0$ | 1 |
| Initial condition | $\vec{q}_0$ | (0, 0) |
| Fate thresholds | $x_{low}; x_{high}$ | -1; +1 |
| Simulation duration | $T$ | 50 |
| Stability penalty weight | $\lambda$ | 0.1 |
| Stability time | $n_{st}$ | 5 |
| Confining potential strength | $A_0$ | 0.005 |
| Noise magnitude | $\eta$ | 0 |
| Evolutionary parameters | | |
| Population size | $P$ | 100 |
| Number of generations | $G$ | 30 |
| Probability of adding | $p_{add}$ | 0.15 |
| Probability of deletion | $p_{drop}$ | 0.15 |
| Prob. of parameter mutation | $p_{mod}$ | 0.7 |
| Evolved parameters | | |
| Module location | $\vec{q}_i = (x_i, y_i)$ | $x_i, y_i \in [-2, 2]$ |
| Module size | $\vec{\sigma}_i = (\sigma_i^1, \sigma_i^2)$ | $\sigma_i^k \in [0.2, 1.5]$ |
| Module strength | $\vec{a}_i = (a_i^1, a_i^2)$ | $a_i^k \in [0, 4]$ |
| Timescale | $\tau$ | $\tau \in (0, 2)$ or $\tau = 5$ |

#### S5.3 Evolution with a target number of segments

The fitness function Eq. S45 increases monotonically with the number of boundaries formed. For the evolution of a specific number of segments, we modify the fitness function using the target number of boundaries $n_b$:

$$\mathcal{F} = -|\mathcal{F}_{stripe} - n_b| - \lambda \mathcal{F}_{stab}. \tag{S48}$$

We provide the parameters used for this set of simulations in the table below; the other parameters are the same as in Table S1.

Table S2: Parameters used for evolution with a fixed segment count

| Parameter | Notation | Value or prior |
|---|---|---|
| Number of boundaries | $n_b$ | 5, 10 or 15 |
| Front propagation | $\Delta t_0$ | 0.5 |
| Confining potential strength | $A_0$ | 0.05 |
| Number of generations | $G$ | 50 |
| Timescale | $\tau$ | $\tau = 5$ |

For each value of $n_b$, we ran 30 evolutionary optimizations. We then kept only those landscapes that formed exactly $n_b$ boundaries and exhibited stable oscillations at time $t_0 - 25$ relative to the transition time $t_0$ (no damped oscillations or long transient dynamics from the initial condition). The results of these simulations are presented in Fig. S3. The oscillator characteristics (period and amplitude) were calculated at time $t_0 - 25$.

To analyze the temporal synchrony of evolved bifurcations (Fig. S4), we identified the timings of two bifurcations in each landscape. The first ($t_1$) is the limit cycle bifurcation (SNIC or homoclinic), involving also one of the fates A or P (either the fate attractor is created by the SNIC, or the saddle of the homoclinic trajectory leads to this attractor). The other bifurcation ($t_2$) is the saddle-node creating the opposing fate attractor (A or P). We then quantify the delay between bifurcations $|t_2 - t_1|$ in evolved landscapes with 5 and 15 boundaries. The samples where the opposing attractor is preexisting ($|t_2 - t_1| = \infty$) were excluded.

### S6 Fitting the neuromesoderm differentiation dataset

#### S6.1 Dataset and landscape setup

We fit data from an *in vitro* system of mouse embryonic stem cells developed by Sáez et al., initial experimental series in [12]. In this work, a flow-cytometry assay was used to measure the expression of marker proteins in single cells at several timepoints. The data were clustered using a Gaussian mixture model to quantify the proportions of cell states – epiblast (Epi), transitioning (Tr), caudal epiblast (CE), mesoderm (M), posterior neural (PN), anterior neural (AN), and unclassified (UT) – at each timepoint.

Starting from the epiblast state, cells were directed to distinct outcomes by combinatorial modulation of FGF and WNT signalling. WNT signalling was controlled by adding activator CHIRON99021 (CHIR) with baseline inhibition of endogenous WNT by LGK974. FGF signalling was controlled by exogenous FGF and FGF inhibitor PD0325901 (PD). In total, 5 combinations were applied:

1. FGF
2. FGF + CHIR
3. CHIR (with PD)
4. CHIR + endogenous FGF
5. Endogenous FGF

We interpret these signalling combinations as regimes (configurations), $k = \overline{1,5}$, of a landscape that changes as a piecewise function of time, Eq. S19. The difference between experimental conditions is in the transition times $t_k$ from one regime to another. We use 7 experiments for evolutionary optimization:

| Exp. | Description | Regimes $k$ | Transition times $t_k$ |
|---|---|---|---|
| 1 | No CHIR | $1 \to 5$ | $t_1 = t_2 = t_3 = t_4 =$ D3 |
| 2 | CHIR 2-3 | $1 \to 2 \to 5$ | $t_1 =$ D2, $t_2 = t_3 = t_4 =$ D3 |
| 3 | CHIR 2-4 | $1 \to 2 \to 4 \to 5$ | $t_1 =$ D2, $t_2 = t_3 =$ D3, $t_4 =$ D4 |
| 4 | CHIR 2-5 | $1 \to 2 \to 4$ | $t_1 =$ D2, $t_2 = t_3 =$ D3 |
| 5 | CHIR 2-5 FGF 0-3 | $1 \to 2 \to 3$ | $t_1 =$ D2, $t_2 =$ D3 |
| 6 | CHIR 2-5 FGF 0-4 | $1 \to 2 \to 3$ | $t_1 =$ D2, $t_2 =$ D4 |
| 7 | CHIR 2-5 FGF 0-5 | $1 \to 2$ | $t_1 =$ D2 |

Table S3: Description of experiments and their parametrization with $k$ and $t_k$, also presented as bar plots in Fig. 5A of the main text.

Based on the cell states defined in [12], we use a fixed number ($S = 6$) of attractor modules, each with pre-assigned cell identity (Epi, Tr, CE, M, PN, AN). Mutations modify the locations and parameters of these modules, without adding new attractors or other module types ($p_{add} = p_{drop} = 0$).

For fitting, we use the proportions of only classified cells $P^{i,j}_{exp}(s)$, rescaled so that $\sum_{s=1}^{S} P^{i,j}_{exp}(s) = 1$. Here, $i$ denotes the index of experiment and $j$ the timepoint. The definition of the fitness function is given below.

#### S6.2 Simulation

The simulation time runs from Day 1.5 until Day 5. We convert data time to simulation time using auxiliary units, $\Delta T = 0.5$ (day) $= 2$ (a.u.), with the initial timepoint Day 1.5 $= 0$. The transition times $t_k$ are then converted to simulation time, and the simulation for one experiment spans the simulation time $0 < t < 7\Delta T$.

We simulate trajectories of $n_{cells}$ cells in the landscape with noisy dynamics and fixed noise magnitude $\eta$. The initial condition for all experiments is the Epi state, so we initialize cells using the coordinates of Epi module and some random noise, $\vec{q}_0 = \vec{q}_{Epi} + \mathcal{N}(0, \eta)$. Trajectories are then integrated for each experiment $i$; at simulation timesteps corresponding to data timepoints $j$ ($t = j\Delta T$), cell states $s(\vec{q}, t)$ are calculated for all cells (Eq. S42-S43), and cell state counts are normalized to obtain proportions $P^{i,j}(s)$, $\sum_{s=1}^{S} P^{i,j}(s) = 1$.

#### S6.3 Fitness function

The fitness function in this case evaluates the agreement between simulation and experiment and is based on the Kullback–Leibler (KL) divergence. For discrete observations (cell states) $s = \overline{1, S}$, the KL divergence of an observed (simulation) distribution $P$ from a target (experiment) distribution $P_{exp}$ is

$$D_{KL}(P_{exp}||P) = \sum_{s=1}^{S} P_{exp}(s) \log \frac{P_{exp}(s)}{P(s)}. \tag{S49}$$

For the fitness function, we sum over timepoints of experiment and over experiments:

$$\mathcal{F} = \sum_{i=1}^{N_{exp}} \sum_{j=1}^{n_t} D_{KL}(P^{i,j}_{exp}||P^{i,j}). \tag{S50}$$

Optionally, we can include penalties to control for cell state stability, based on biological knowledge:

$$\overline{\mathcal{F}} = \mathcal{F} - \lambda \mathcal{F}_{stab}. \tag{S51}$$

For example, if a state $i$ is known to be stable at time $t_s$, we verify its stability in the landscape by running a long deterministic trajectory starting from the module location $\vec{q}_i$: $(\vec{q}_i, t_s) \to \vec{q}(t), t \in [t_s, t_s + T]$, with landscape parameters fixed at time $t_s$. We then check the cell state at the end of this trajectory using Eq. S43. For a stable attractor, we should obtain $s(\vec{q}(t_s + T), t_s + T) = i$. Conversely, if a state is known to be transitory, the attractor never stabilizes and will be escaped, so we should obtain $s(\vec{q}(t_s + T), t_s + T) \neq i$.

The penalty term is then:

$$\mathcal{F}_{stab} = \sum_{i \in \text{stable}} \sum_{t_s} \left( s(\vec{q}(t_s + T), t_s + T) \neq i \right) + \sum_{j \in \text{transient}} \sum_{t_s} \left( s(\vec{q}(t_s + T), t_s + T) = j \right). \tag{S52}$$

For instance, one can enforce that the differentiated (committed) states occupied by cells at the end of experiment are stable ($t_s$ is then the final experimental timepoint). Cell states observed only briefly (i.e., at a single timepoint $t_s$) can be imposed to be transient.

#### S6.4 Iterative optimization

The parameter values for each of the configurations $a_i^k$, $\sigma_i^k$ are fitted independently, without assuming a linear dependency on the signals. For the dataset of $N_{exp} = 7$ experiments with $S = 6$ cell states (6 attractor modules) and $K = 5$ applied morphogen combinations, the number of parameters is 72 (2 coordinates and 5 levels of $a_i$ and $\sigma_i$ for every module). We use a population of size $P = 200$, with 100 mutations per generation. To further reduce the number of parameters compared to the number of mutations, we split the optimization into several steps, based on time courses and the specifics of the dataset.

First, we selected 3 experiments (experiments 5-7) for 'Stage 1' optimization. In these experiments, three morphogen regimes are applied (FGF, CHIR, and FGF+CHIR), $K = 3$, and 5 cell states are observed (all except for AN). We found that these experiments offer a good representation of different cell state proportions as well as the main external signals. Furthermore, the initial 4 timepoints (Day 1.5 - Day 3) are common between experiments 2-7. These constitute 'Stage 0' optimization. Finally, all 7 experiments are fitted in 'Stage 2'. For each of the optimization steps, we define a fitness function (Eqs. S49-S50) using the corresponding number of experiments $N_{exp}$, number of timepoints $n_s$, applied signals (configurations) $K$, and observed cell states $S$. These numbers and the number of parameters fitted at each stage $n_{fit}$ are given in Table S4.

Table S4: Stages of iterative optimization

| | $N_{exp}$ | Experiments | $n_t$ | $S$ | States | $K$ | $n_{fit}$ |
|---|---|---|---|---|---|---|---|
| Stage 0 | 1 | Exp. 2-7 D1.5-D3 | 4 | 3 | Epi, Tr, CE | 3 | 24 |
| Stage 1 | 3 | Exp. 5-7 | 8 | 5 | Epi, Tr, CE, M, PN | 3 | 40 |
| Stage 2 | 7 | Exp 1-7 | 8 | 6 | Epi, Tr, CE, M, PN, AN | 5 | 48 |

In Stage 0, the observed cell states are Epi, Tr, CE, and a small proportion of M at timepoint 4 (Day 3). We initially set $P_{CE} = 1$, $P_M = 0$ for timepoint 4 and optimize landscapes with three modules. While this pre-optimization is not a necessary step, we observed that it significantly increases the success rate for capturing the initial timepoints, in particular for the briefly transient state Tr, which is only observed at timepoint 3 (Day 2.5). This exemplifies how time courses can be used to sequentially add attractors to the landscape, as opposed to adding binary decisions.

For Stage 1, we use the pre-optimized landscapes with Epi, Tr, and CE modules as the initial condition, while also adding two new randomly generated modules for states PN and M. The Stage 0-Stage 1 procedure was run $N = 1200$ times to obtain a distribution of landscape parameters. We then used a fitness threshold to select 240 best landscapes (Fig. S6) that proceeded to Stage 2. The parameter vectors $\vec{a}_i$ and $\vec{\sigma}_i$ of all modules were expanded from 3 elements to 5 to account for new signalling combinations. We initialize the endogenous FGF regime with parameter values from the FGF regime, and analogously, the CHIR+endogenous FGF with the CHIR+FGF values. We also add another randomly generated attractor module to each landscape for the AN state. In Stage 2, we fix the locations of the Epi, Tr, CE, and M modules and their parameters in the FGF and CHIR+FGF regimes, while keeping all parameters and locations free for the AN and PN modules.

The computation time for one evolutionary optimization in a population of 200 landscapes is under 1 minute for Stage 0, around 7 minutes for Stage 1, and 20 minutes for Stage 2. The best landscape from each optimization is one sample in the evolutionary ensemble that we analyze.

Table S5: Parameters used for neuromesoderm landscape evolution

| **Parameter** | **Notation** | **Value or prior** |
|---|---|---|
| Simulation parameters | | |
| Number of cells - stage 0-1 | $n_{cells}$ | 300 |
| Number of cells - stage 2 | $n_{cells}$ | 500 |
| Timepoint duration | $\Delta T$ | 2 |
| Confining potential strength | $A_0$ | 0.005 |
| Noise magnitude | $\eta$ | 0.2 |
| Evolutionary parameters | | |
| Population size | $P$ | 200 |
| No. of generations - stage 0 | $G_0$ | 100 |
| No. of generations - stage 1 | $G_1$ | 300 |
| No. of generations - stage 2 | $G_2$ | 300 |
| Prob. of adding/deleting | $p_{add} = p_{drop}$ | 0 |
| Prob. of parameter modification | $p_{mod}$ | 1 |
| Evolved parameters - stages 0-1 | | |
| Module location | $\vec{q}_i = (x_i, y_i)$ | $x_i, y_i \in [-4, 4]$ |
| Module size | $\vec{\sigma}_i \in \mathbb{R}^3$ | $\sigma_i^k \in [0.1, 1.5]$ |
| Module strength | $\vec{a}_i \in \mathbb{R}^3$ | $a_i^k \in [0, 16]$ |
| Evolved parameters - stage 2 | | |
| Module location AN and PN | $\vec{q}_i = (x_i, y_i)$ | $x_i, y_i \in [-5, 5]$ |
| Module size | $\vec{\sigma}_i \in \mathbb{R}^5$ | $\sigma_i^k \in [0.1, 1.5]$ |
| Module strength | $\vec{a}_i \in \mathbb{R}^5$ | $a_i^k \in [0, 16]$ |

| Fixed parameters - stage 2 | |
|---|---|
| Parameter | Notation |
| Module location Epi, Tr, CE, M | $\vec{q}_i = (x_i, y_i)$ |
| Module size Epi, Tr, CE, M - 2 regimes | $\sigma_i^k, k = 2, 3$ |
| Module strength Epi, Tr, CE, M - 2 regimes | $a_i^k, k = 2, 3$ |

#### S6.5 Coordinate system standardization

For landscape comparison and topography analysis, we transform the coordinate system using translation, rotation, and reflection. The distances and angles are preserved under such a transformation.

1. Translation to locate the origin of the coordinate system at CE module: $\vec{q} \to \vec{q} - \vec{q}_{CE}$;

2. Rotation to align the $x$-axis with the average direction (bisector) from origin CE to PN and M: $\vec{q} \to R\,\vec{q}$ with

$$\vec{q}_{avg} = (x_{avg},\, y_{avg}) = \frac{\vec{q}_{PN}}{|\vec{q}_{PN}|} + \frac{\vec{q}_M}{|\vec{q}_M|} \tag{S53}$$

and

$$R = \frac{1}{|\vec{q}_{avg}|} \begin{pmatrix} x_{avg} & y_{avg} \\ -y_{avg} & x_{avg} \end{pmatrix} \tag{S54}$$

This way, the $x$-axis passes between PN and M;

3. If needed, a reflection of the $x$-axis so that $x_{Epi} < 0$ (Epi is to the left of CE);
4. If needed, a reflection of the $y$-axis so that M is in the bottom half-plane, $y_M < 0$ ($y_{PN} > 0$).

#### S6.6 Topology analysis and landscape categorization

The topological analysis of binary decisions is based on the classification developed in [1], Fig. S9. A binary decision includes three attractors, two saddles, and their bifurcation set. In our framework, a binary decision is realized with three attractor modules. The modules implicated in a decision can be identified from streamplots with fixed points or from simulated trajectories (trajectories from a state leading to two other states).

We represent binary decisions using two types of graphs: differentiation tree and global (in parameter space) decision structure. The *differentiation tree* is a directed graph of observed cell state transitions in simulated trajectories. Based on the differentiation tree, we define the central attractor of a binary decision as the initial (less differentiated) state and the fate attractors as the two more differentiated states. The *global decision structure* is an undirected graph where the edges indicate saddle connections (through unstable manifolds) between attractors. For a landscape with fixed $\vec{q}_i$, we include in the decision structure all saddle connections observed at different values of shape parameters $a_i$, $\sigma_i$, rather than only those observed for given parameter values. As in Fig. 5 of the main text, both types of graphs can be extended to include sequences of binary decisions and intermediate states. The differentiation tree can also include non-attractor (transitory) states.

The *double cusp (binary choice)* landscape (Fig. S9 A) is a region of parameter space where two saddle-node bifurcation curves of the center attractor meet in a cusp [1]. The *heteroclinic flip* landscape (Fig. S9 B) is a region of parameter space that contains a flip bifurcation curve [1], where the saddle connection from the central attractor switches between the two fate attractors. Although the differentiation tree can be the same for both topologies (central $\rightarrow$ fate 1 and central $\rightarrow$ fate 2), they differ by the decision structure as shown in Fig. 6 of the main text.

Thus, to classify landscapes, we first identified saddle connections visually (Fig. S9, discrete landscapes). Second, we detect the connections using the simulated *basin escape dynamics*. For the CE-M/PN decision, we initialize 300 cells in the fate attractor M, $\vec{q}_0 = \vec{q}_M + \mathcal{N}(0, 0.2\eta)$. We use parameters for static CHIR signalling ($k = 3$) where both fate attractors are present. We integrate cell trajectories with increased noise magnitude $\eta^* = 6\eta$ or $\eta^* = 8\eta$ for duration $0 < t < 5\Delta T$. The presence of cell trajectories going from M directly to PN is used as a criterion for the heteroclinic flip landscape.

Additionally, by visual inspection, we identified 10 optimized landscapes that did not have a binary decision structure. Rather, they contain one of the fates as a transient state, or mixed states where several modules are merged into one attractor. These landscapes can reproduce experimental proportions as different regions of the mixed attractor are assigned to different states. We consider these *assignment-based* solutions unsuitable since our objective is to find *attractor-based* solutions, consistent with the dynamical systems theory of cell fate decisions.

The resulting landscape categorization for 60 landscapes is provided in the Supplementary Data.

#### S6.7 Landscape regularization

##### Temporal regularization

We explore regularization strategies to reduce the number of parameters and/or impose additional constraints on the landscape. First, landscapes can be temporally regularized by suppressing modules while the corresponding cell state is not observed. Namely, if state $s$ is not observed in regime $k$ ($P^{i,j}_{exp}(s) \leq 0.01$ for all experiments $i$ and timepoints $j$ when signalling is in regime $k$), we set $a^k_s = 0$ and $\sigma^k_s = 0$ (Fig. S12 A). For this dataset, we then need to optimize 16 discrete values for $a$ and $\sigma$ each. The number of shape parameters is thus reduced to 32, and we keep 12 location parameters ($\vec{q_i}$) to be optimized.

Due to the smaller number of parameters, these optimizations were performed with Stage 2 directly after Stage 0: we pre-optimize only for 4 initial timepoints with Epi, Tr, and CE modules, and then evolve for all 7 experiments at once, without fixing any of the parameters. The priors and other optimization parameters are the same as in the main simulations, Table S5, except for the timescale (duration between experimental timepoints) $\Delta T$. We noticed that after module suppression, the velocities in the landscape can be reduced; in our main simulations, 'irrelevant' modules might actually play a role in shaping transient trajectories, thus canalizing and speeding up the landscape. To compensate for the reduced number of modules, we increase $\Delta T$ to 3 a.u.

Results obtained with such regularized models are very similar to the main simulations. In particular, we still find the two bifurcation scenarios for the CE-PN/M decision (Fig. S12). We also find the same differentiation tree, including rare transitions such as PN $\rightarrow$ AN (Fig. S12C, Experiment 2).

##### Linear regularization

Another possibility is to impose a specific form of parameter dependency on the signal/morphogen input in Eq. S17. In the main simulations, we use a piecewise function for the most general description. Alternatively, we can assume a linear dependency on the signals:

$$a_i(t) = a^i_0 + a^i_1 S_1(t) + a^i_2 S_2(t); \tag{S55}$$

$$\sigma_i(t) = \sigma^i_0 + \sigma^i_1 S_1(t) + \sigma^i_2 S_2(t), \tag{S56}$$

where $S_1(t)$ is the level of CHIR and $S_2(t)$ the level of FGF. To keep the resulting parameters within a reasonable range, we also clip them to $[0, a_{max}]$ and $[\sigma_{min}, \sigma_{max}]$, respectively. This prevents very narrow peaks and attractors turning into repellers.

We use the same values of the signals as in [12], so the 5 discrete combinations are translated to signal levels in the following way:

| Regime | $S_1$ | $S_2$ |
|---|---|---|
| FGF (no CHIR) | 0 | 1 |
| FGF + CHIR | 1 | 1 |
| CHIR (with PD) | 1 | 0 |
| CHIR + endogenous FGF | 1 | 0.9 |
| Endogenous FGF (no CHIR) | 0 | 0.9 |

The parameter vectors $\vec{a}_i$ and $\vec{\sigma}_i$ are then reduced to three elements for each module; we thus have 36 shape parameters and 12 location parameters. We also use tighter priors for module locations, based on the knowledge of their relative positions in the main simulations. The parameters used are provided in Table S6. Again, we omit Stage 1 of iterative optimization and optimize first for 4 initial timepoints, then for all experiments at once. Due to a known nonlinearity (memory effect) in the data from Experiment 3 [12], we lower the relative weight of this experiment in the fitness function when optimizing landscapes with the linear assumption. In Eq. S50, the corresponding KL term was multiplied by 0.1.

As expected, solutions are more difficult to find for landscapes with the linear constraint (any change of $\vec{a}_i, \vec{\sigma}_i$ parameter vectors impacts all experiment simulations). We obtain less landscapes with a high fitness and a more heavy-tailed fitness distribution. Optionally, to increase the convergence rate of simulations with the linear constraint, we can restrict module responses to specific signals to positive or negative, based on the data. For instance, cells leave the Epi state when CHIR is added (Exp. 2-7) or FGF level is lowered (Exp. 1), so Epi stability is anticorrelated with CHIR and correlated with FGF. We can thus set $a_1 < 0, \sigma_1 < 0$ and $a_2 > 0, \sigma_2 > 0$ for the Epi module. Similarly, we set $a_1 < 0, \sigma_1 < 0$ and $a_2 < 0, \sigma_2 < 0$ for the PN and AN modules (anticorrelated with both CHIR and FGF).

The successful optimizations of linear landscapes are consistent with our results for the more general, piecewise parameterization (Fig. S13). Again, we recover the two types of final decision (double cusp and heteroclinic flip). With the linear parameterization, we interpolate the landscape for intermediate levels of input signals to observe bifurcations - crossings of the boundaries in the catastrophe diagram (Fig. S9). In particular, for the heteroclinic flip example, the flip bifurcation can be found at CHIR = 1 and FGF $\approx 0.08$. Fo both decision topologies, the bifurcation space trajectories estimated from such continuous modulation are consistent with the mappings inferred from the discrete landscapes (Fig. S9).

Table S6: Parameters used for linearized landscape evolution

| **Parameter** | **Notation** | **Value or prior** |
|---|---|---|
| Simulation parameters | | |
| Number of cells | $n_{cells}$ | 300 |
| Timepoint duration | $\Delta T$ | 3 |
| Confining potential strength | $A_0$ | 0.005 |
| Noise magnitude | $\eta$ | 0.2 |
| Evolutionary parameters | | |
| Population size | $P$ | 200 |
| No. of generations - stage 0 | $G_0$ | 100 |
| No. of generations - all exp. | $G_1$ | 300 |
| Prob. of adding/deleting | $p_{add} = p_{drop}$ | 0 |
| Prob. of parameter modification | $p_{mod}$ | 1 |
| Evolved parameters | | |
| Module location Epi | $\vec{q}_1$ | $x_1 \in [-4, 0], y_1 \in [-3, 3]$ |
| Module location Tr | $\vec{q}_2$ | $x_2 \in [-2, 0], y_2 \in [-2, 2]$ |
| Module location CE | $\vec{q}_3$ | $x_3 \in [-1, 1], y_3 \in [-1, 1]$ |
| Module location PN | $\vec{q}_4$ | $x_4 \in [0, 4], y_4 \in [0, 4]$ |
| Module location M | $\vec{q}_5$ | $x_5 \in [0, 4], y_5 \in [-4, 0]$ |
| Module location AN | $\vec{q}_6$ | $x_6 \in [-5, -1], y_6 \in [-4, 4]$ |
| Minimum module size | $\sigma_{min}$ | 0.2 |
| Maximum module size | $\sigma_{max}$ | 1.2 |
| Module size (no signal) | $\sigma_0^i$ | $[0.2, 1.2]$ |
| Size response | $\sigma_1^i, \sigma_2^i$ | $[-1, 1]$ |
| Maximum module strength | $a_{max}$ | 20 |
| Module strength (no signal) | $a_0^i$ | $[0, 20]$ |
| Strength response | $a_1^i, a_2^i$ | $[-20, 20]$ |

### Supplementary Data

We provide a PDF showing the behavior of the 60 best landscape samples for neuromesoderm differentiation, using the fitness threshold mentioned in the main text to select the samples.

The layout of each page is as follows:

A. Cellular trajectories in a simulation for experiment 5;

B. Left: Basin escape numerical experiment with increased noise magnitude in the CHIR signalling regime. All cells were initialized in M state. Cellular trajectories and a potential contour plot are shown. Right: stream plot of the dynamics in the CHIR signalling regime. Crossings of nullclines indicate fixed points. Diamonds indicate locations of modules;

C. Potential cross sections along green vertical lines in B;

D. Potential surface between the first and last cross section;

E. Comparison of simulated cell state proportions to fitted data over 7 experiments.

Landscapes were categorized based on the topology of the CE - PN/M decision, inferred from the locations of the saddles and the escape experiment. Two binary decision topologies are observed:

- *Heteroclinic flip*: triangular topology with a saddle between PN and M. A subcategory *Heteroclinic flip with overlap* was separated based on significant spatial overlap of the flip landscape with the initial decision (Epi - Tr - CE)
- *Double cusp*: linear topology, with no direct path from M to PN. A subcategory *Flat double cusp* was defined based on the shape of the potential.

Finally, the *Transient/mixed state* category contains landscapes where PN or M acts as a transient state, or where two modules are merged into one attractor. These have assignment-based spatial structure rather than attractor-based.

### Supplementary Movies

In this section, we briefly describe the Supplementary Movies.

**Supplementary Movie 1**: Dynamics of an evolved landscape for somitogenesis, with a *SNIC* bifurcation. Cell trajectories are shown, color-coded by initial condition (oscillation phase). Nullcline crossings indicate fixed points. The attractor module on the left creates the bifurcation.

**Supplementary Movie 2**: Dynamics of an evolved laandscape for somitogenesis, with a *homoclinic* bifurcation. Cell trajectories are shown, color-coded by initial condition (oscillation phase). Nullcline crossings indicate fixed points. The saddle point between the oscillator and the top attractor creates the bifurcation.

**Supplementary Movie 3**: Cell fate dynamics in neuromesoderm differentiation. Simulated trajectories in an evolved landscape with a *heteroclinic flip* topology, for *experiment 2* (FGF $\rightarrow$ CHIR+FGF $\rightarrow$ end. FGF). Cells are color-coded by current state.

**Supplementary Movie 4**: Simulated trajectories in an evolved landscape with a *heteroclinic flip* topology, for *experiment 5* (FGF $\rightarrow$ CHIR+FGF $\rightarrow$ CHIR). Cells are color-coded by current state.

**Supplementary Movie 5**: Simulated trajectories in an evolved landscape with a *double cusp* topology, for *experiment 2* (FGF $\rightarrow$ CHIR+FGF $\rightarrow$ end. FGF). Cells are color-coded by current state.

**Supplementary Movie 6**: Simulated trajectories in an evolved landscape with a *double cusp* topology, for *experiment 5* (FGF $\rightarrow$ CHIR+FGF $\rightarrow$ CHIR). Cells are color-coded by current state.

### Supplementary Figures

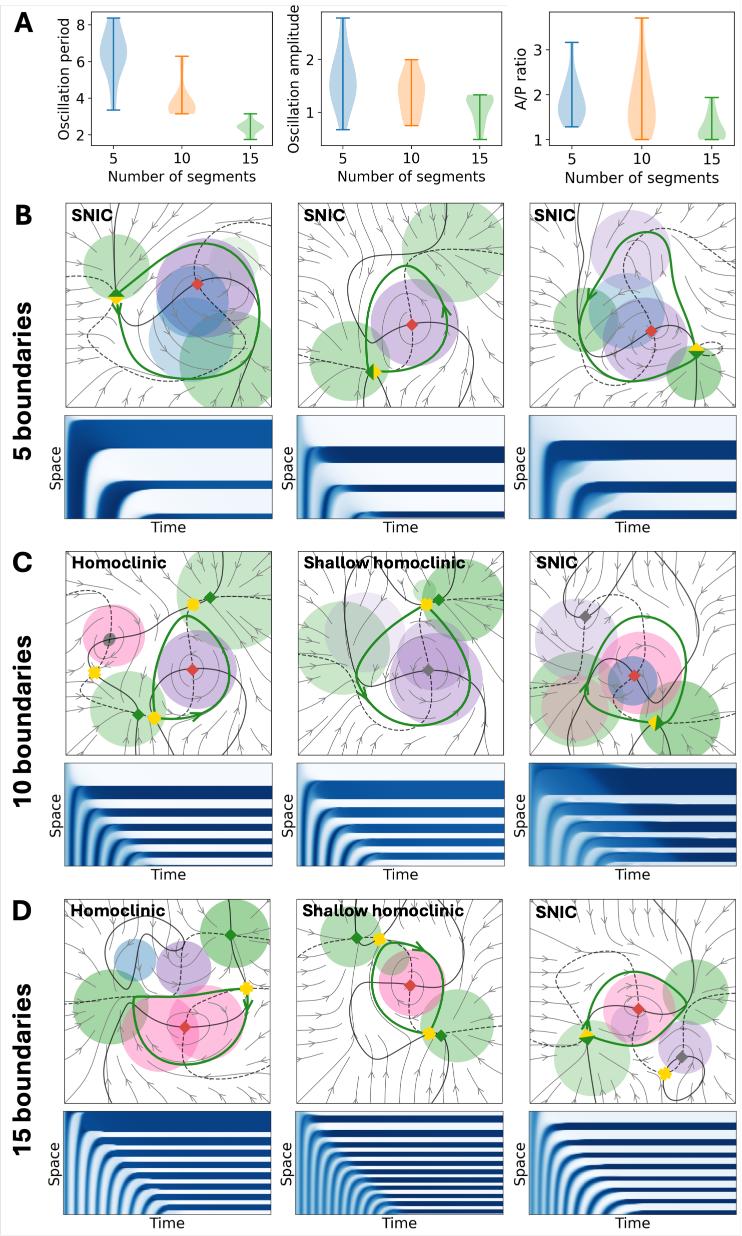


Figure S3: Evolved landscapes with 5, 10, and 15 segment boundaries. **A)** Oscillation period (estimated from the peak of the frequency spectrum), oscillation amplitude ($x$ variable), and the ratio of A/P cells in the resulting pattern. The number of evolved landscapes analyzed is $N = 13$ for 5, $N = 15$ for 10, and $N = 11$ for 15 boundaries. **(B-D)** Representative landscapes, shown close to the limit cycle bifurcation, and their kymographs $x(p, t)$. **B)** Landscapes producing 5 boundaries almost always bifurcate through a SNIC. Slow cycles are created by using large repellers and overlapping the cycle with future attractor modules. **C)** With 10 boundaries, SNIC and homoclinic bifurcations are obtained with approximately equal probabilities. The homoclinic bifurcation can occur with a preexisting attractor outside the cycle, or soon after a saddle-node close to the cycle ('shallow homoclinic'). **D)** With 15 boundaries, we observe a nearly equal mix of SNIC and homoclinic bifurcations (and occasionally, Hopf bifurcations). The bifurcation of the limit cycle becomes more spatially symmetric and temporally synchronized to the saddle-node of the opposing attractor.

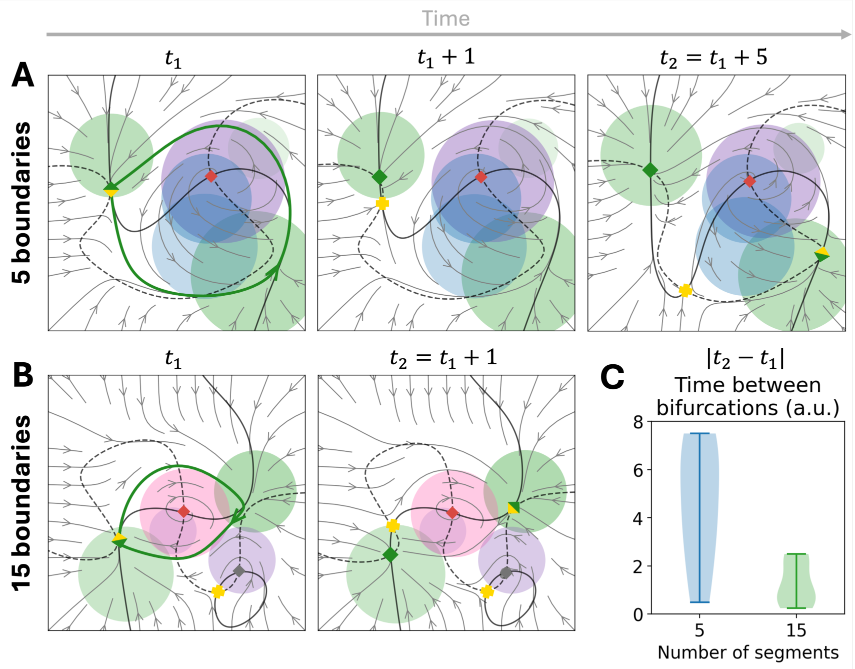


Figure S4: Temporal synchrony in bifurcations of the segmentation landscape. **A)** Example landscape producing 5 boundaries: a SNIC stops the oscillation and creates the anterior state (left attractor) at time $t_1$. After a delay of 5 a.u., a saddle-node creates the posterior state (right attractor). **B)** Example landscape with 15 boundaries: the saddle-node happens 1 a.u. after the SNIC. **C)** Observed delays between bifurcations (SNIC or homoclinic and saddle-node) in landscapes evolved for 5 and 15 boundaries. The absolute time difference is reported; $t_2 < t_1$ if the saddle-node happens earlier than the limit cycle bifurcation. $N = 9$ for 5 and $N = 8$ for 15 boundaries.

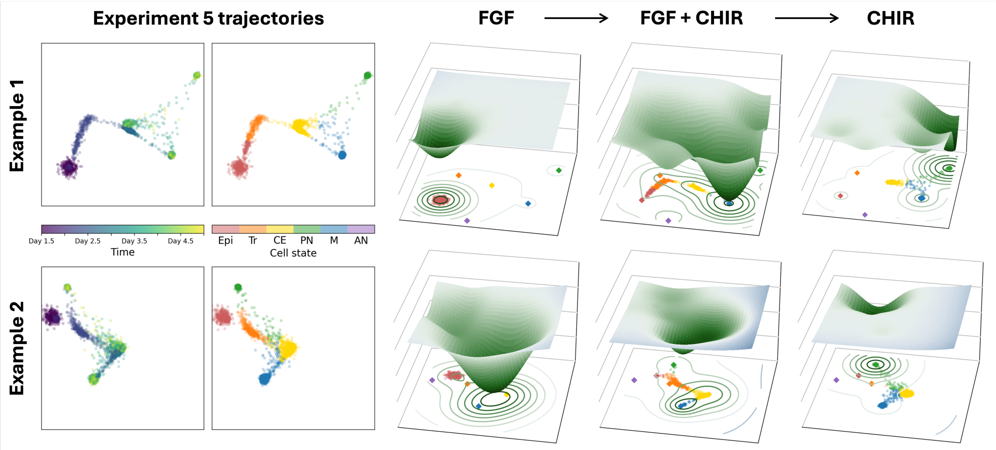


Figure S5: Cellular trajectories and sequential changes of the potential for Experiment 5. With each potential, the corresponding portion of the trajectory is shown. Example 1: a landscape organized along a differentiation axis (left to right). Example 2: a landscape without a clear differentiation axis.

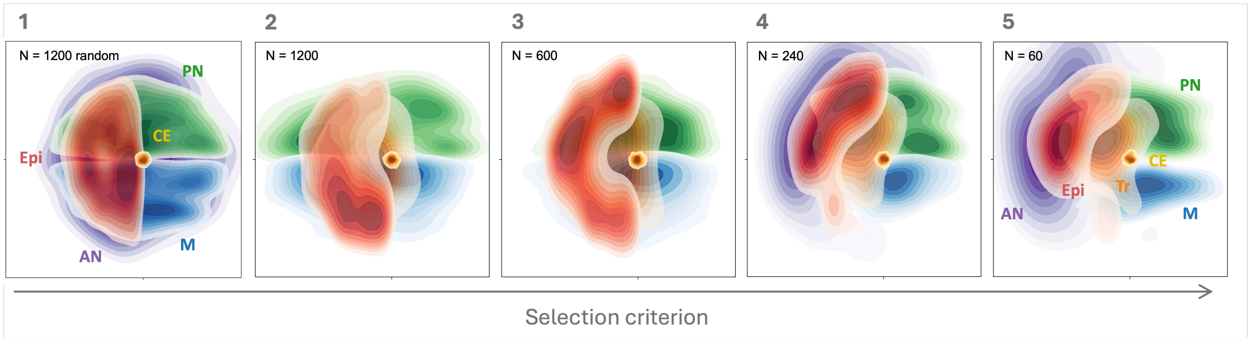


Figure S6: Emergence of the constrained landscape topography. From panel 1 to panel 5, a progressively stricter selection criterion is applied. The distributions of module locations are shown. **1)** No selection: random initialization of module locations in the standardized coordinate system – CE at the origin, Epi in the left half-plane, PN in the top half-plane, M in the bottom half-plane, AN and Tr anywhere in the region. Tr location distribution (full circle) not shown. **2)** All 1200 landscapes evolved in Stage 1 (fitting experiments 5-7). **3)** 600 landscapes selected from Stage 1 using a fitness threshold. **4)** 240 landscapes selected from Stage 1 and evolved in Stage 2 optimization (experiments 1-7). **5)** Best 60 landscapes. This topography is presented in Figure 5E of the main text.

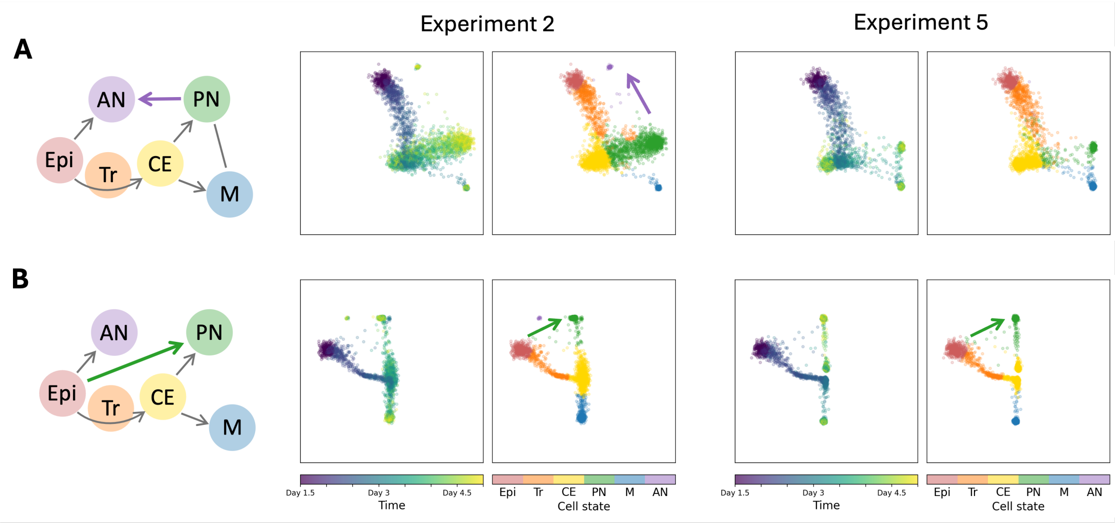


Figure S7: Example of transitions from PN to AN (observed in Experiment 5 simulation) and from Epi to PN (observed in Experiment 2 and Experiment 5 simulations).

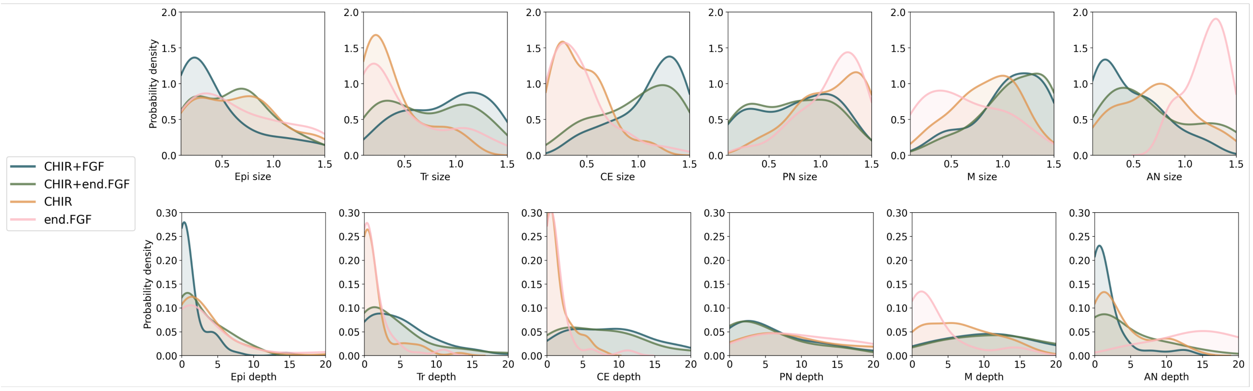


Figure S8: Distributions of sizes and depths of modules in different signalling regimes. Statistics from $N = 60$ landscapes are shown.

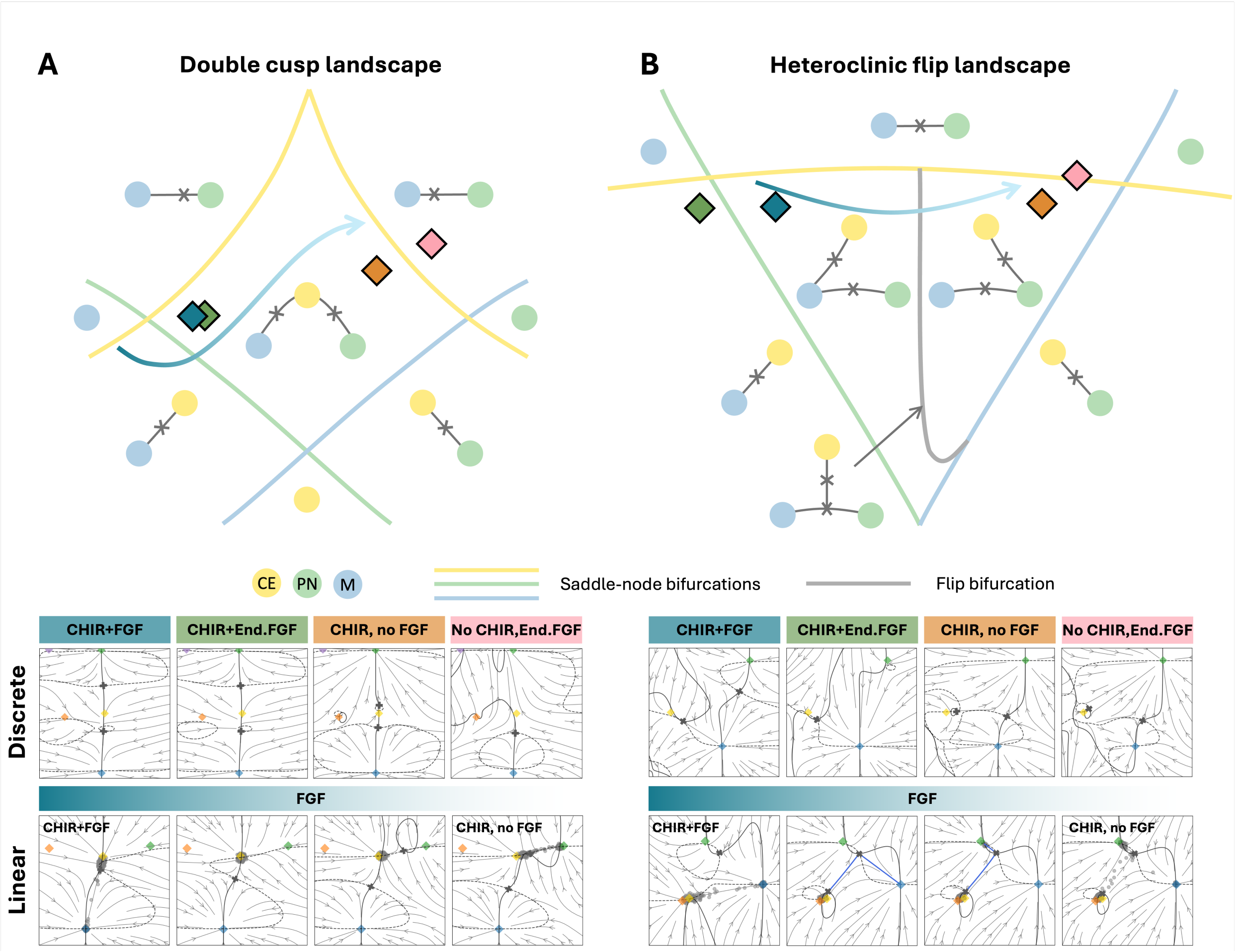


Figure S9: Identification of the decision structure for the CE-PN/M decision from optimized landscapes. For the two landscape types, we show the catastrophe diagram schematic, adapted from [1], and examples of optimized landscapes (bottom). The observed configurations are approximately mapped to regions of the catastrophe diagram (Morse-Smale components). For landscapes with discrete signalling regimes (main simulations), the estimated locations in the bifurcation space are shown with diamond markers. For landscape with linear combinations (supplementary simulations), we show an interpolation for the FGF signal decreasing from exogenous FGF (1) to FGF inhibition (0), with CHIR kept constant (CHIR = 1). The estimated trajectory in the bifurcation space is shown with an arrow.

**A)** Double cusp landscape. Discrete configurations: in CHIR+FGF and CHIR regimes, three attractors form a 1D geometry with two saddles. In CHIR+FGF signalling, CE is close to bifurcation with the M saddle, while in CHIR - close to the PN saddle. Linear landscape: when modulating the level of FGF signalling, we observe how CE moves away from a bifurcation with the M saddle, PN stabilizes through a saddle-node, then CE approaches bifurcation with the PN saddle.

**B)** Heteroclinic flip landscape. Discrete configurations: in regimes with all three attractors present, a saddle connects the PN and M states. CHIR+FGF and CHIR configurations are located on different sides of the flip bifurcation curve, as seen from the escape route of CE towards M or PN. Linear landscape: at low levels of FGF, we see a flip bifurcation where the escape route of CE flips from M to PN. The escape route of the CE saddle before and after the flip is shown in the streamplots in blue.

In both types of landscape, the preferential transition shifts from CE→ M to CE→ PN, as shown with simulated trajectories starting from CE (grey scatter points).

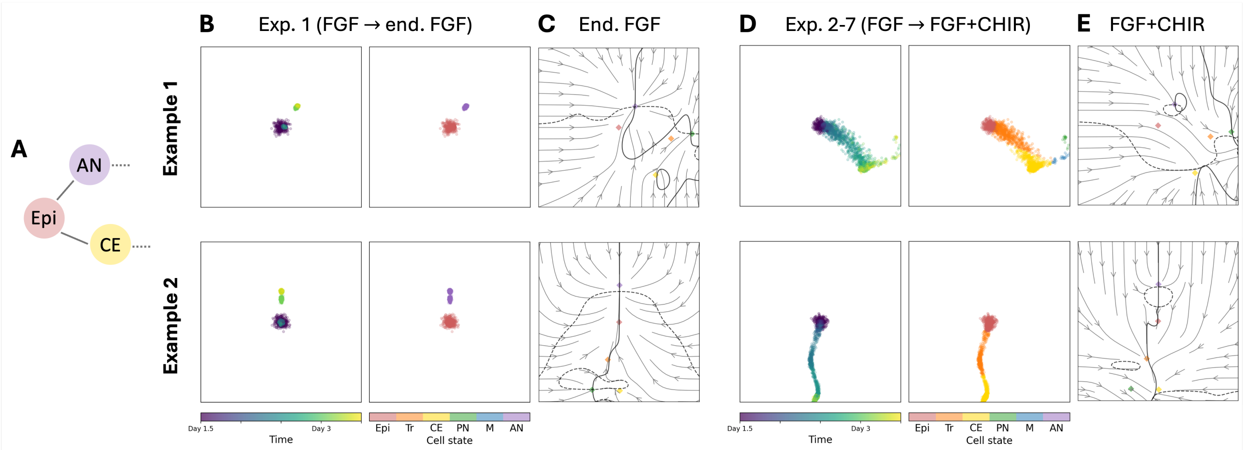


Figure S10: Topology of the initial binary decision of the landscape. **A)** The decision structure contains connections from Epi to AN and from Epi to CE. **B)** Examples of simulated cellular trajectories for Experiment 1 (FGF → endogenous FGF). **C)** Dynamics in the Endogenous FGF regime. The Epi attractor bifurcates towards AN. **D)** Simulated cellular trajectories for the initial part of Experiments 2-7 (FGF → FGF + CHIR). **E)** Dynamics in the FGF+CHIR regime. The Epi attractor bifurcates towards CE, with a non-attractor state Tr along the route.

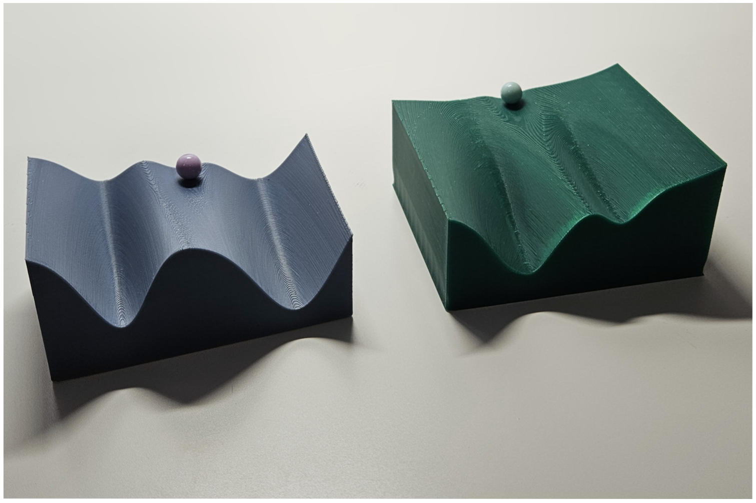

Figure S11: 3D-printed examples of optimized landscapes for the CE-M/PN decision: binary cusp (left) and heteroclinic flip (right). A slice of the potential surface along the differentiation axis is shown. The bead represents a cell in the central (least differentiated) attractor, here CE.

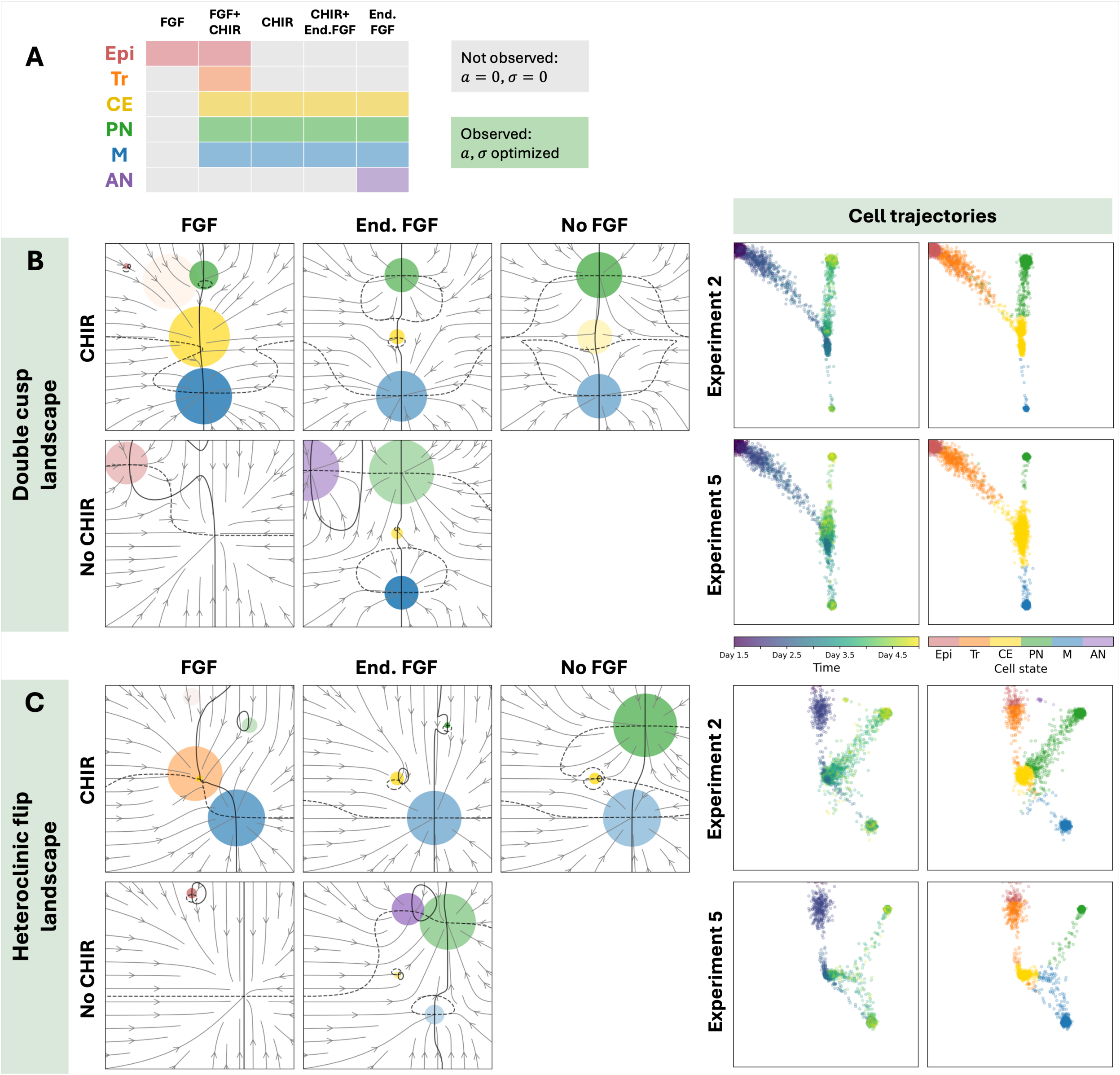


Figure S12: Temporally regularized landscapes. **A)** When a cell state is not observed in the data for a given signal combination, module parameters are set to 0. Other parameters are optimized as in the main simulations. **B)** Evolved double cusp landscape: landscape configurations in different signalling conditions and examples of cellular trajectories. Saddles connect CE to PN and M. **C)** Evolved heteroclinic flip landscape: landscape configurations in different signalling conditions and examples of cellular trajectories. There is a saddle connection between PN and M (e.g, CHIR, no FGF regime).

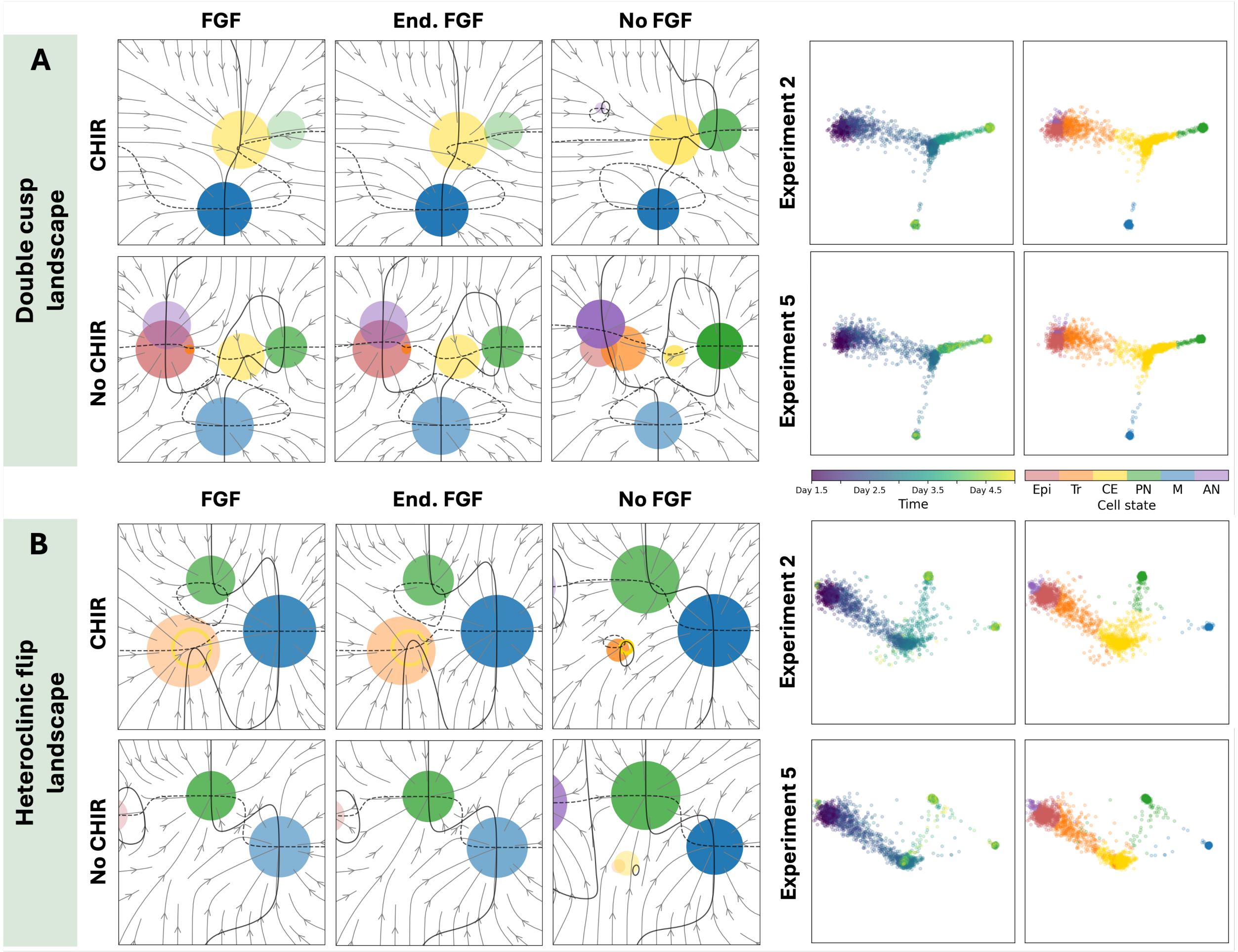


Figure S13: Landscapes with a linear combination of input signals. **A)** Evolved double cusp landscape: landscape configurations in different signalling conditions (with no CHIR, no FGF as prediction) and examples of cellular trajectories. **B)** Evolved heteroclinic flip landscape: landscape configurations in different signalling conditions (no CHIR, no FGF is predicted) and examples of cellular trajectories. The same landscape examples are used in Fig. S9 to illustrate the effect of continuous signal modulation and obtain bifurcations.
