## Supplementary Data 1 for "Generative epigenetic landscapes map the topology and topography of cell fates"

Epi Tr CE PN M AN

In this document, we present 60 evolved landscape samples for neuromesoderm differentiation (data from Sáez et al., 2022).

The layout of each page is as follows:

A. Cellular trajectories in a simulation for experiment 5;
B. Left: Basin escape numerical experiment with increased noise magnitude in the CHIR signalling regime. All cells were initialized in M state. Cellular trajectories and a potential contour plot are shown. Right: stream plot of the dynamics in the CHIR signalling regime. Crossings of nullclines indicate fixed points. Diamonds indicate locations of modules;
C. Potential cross sections along green vertical lines in B;
D. Potential surface between the first and last cross section;
E. Comparison of simulated cell state proportions to fitted data over 7 experiments.

Landscapes were categorized based on the topology of the CE - PN/M decision, inferred from the locations of the saddles and the escape experiment. Two binary decision topologies are observed:
**Heteroclinic flip**: triangular topology with a saddle between PN and M. A subcategory *Heteroclinic flip with overlap* was separated based on significant spatial overlap of the flip landscape with the initial decision (Epi - Tr - CE)
**Double cusp**: linear topology, with no direct path from M to PN. A subcategory *Flat double cusp* was defined based on the shape of the potential.
Finally, the *Transient/mixed state* category contains landscapes where PN or M acts as a transient state, or where two modules are merged into one attractor. These have assignment-based spatial structure rather than attractor-based.

### Heteroclinic flip landscapes

### 00 — Timecode: 20250530-162508

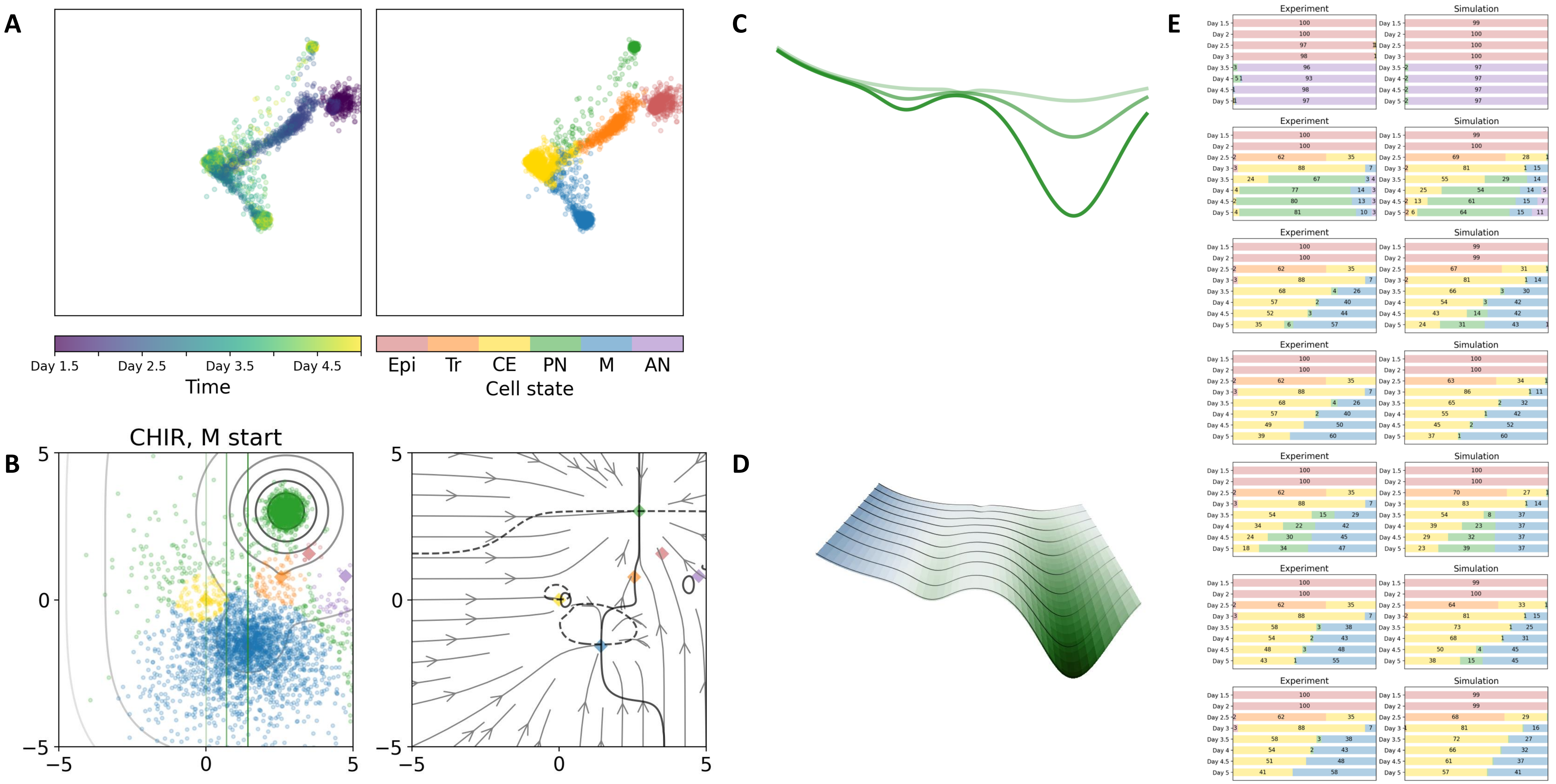

### 02 — Timecode: 20250530-183729

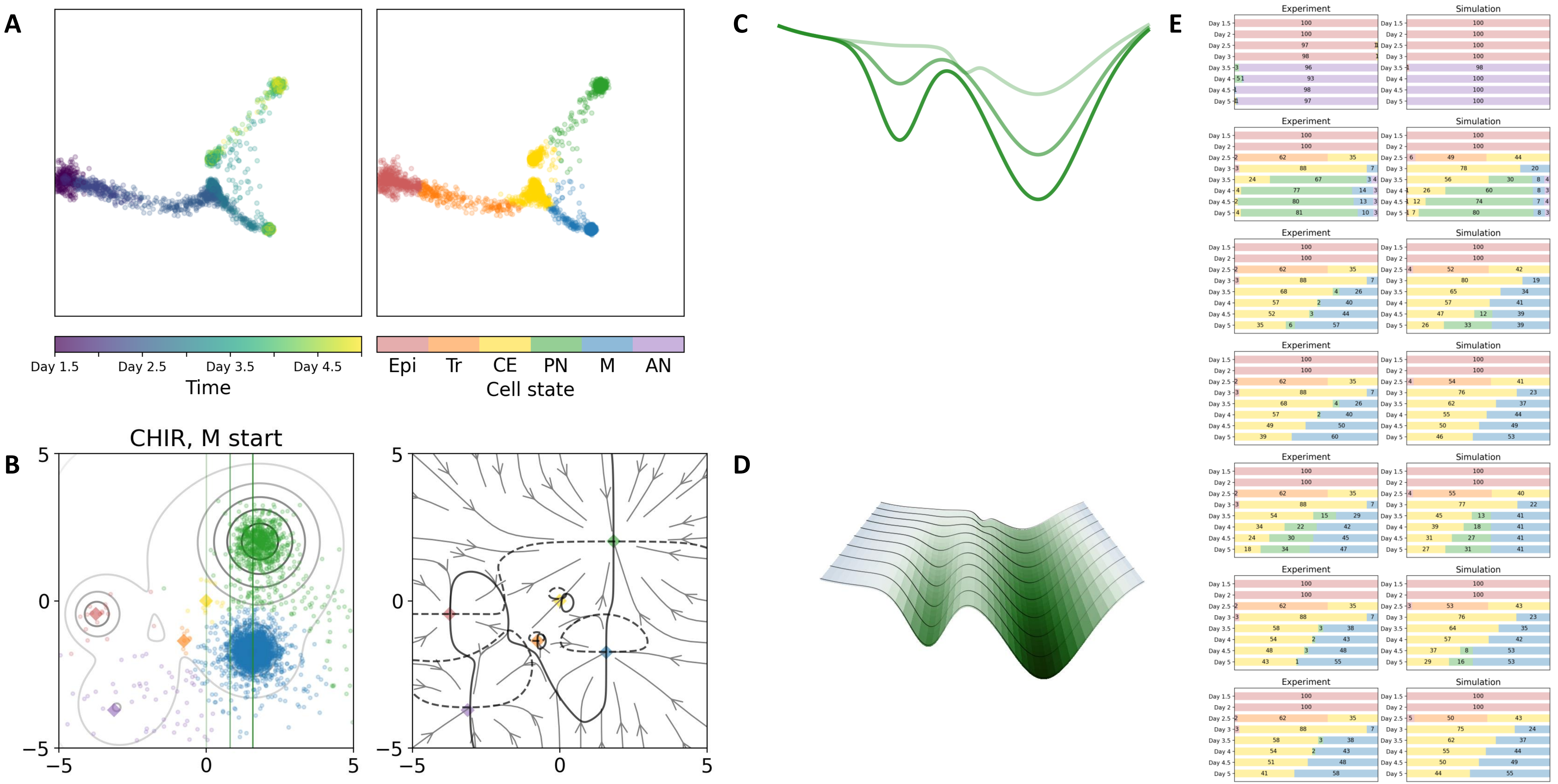

### 07 — Timecode: 20250531-011758

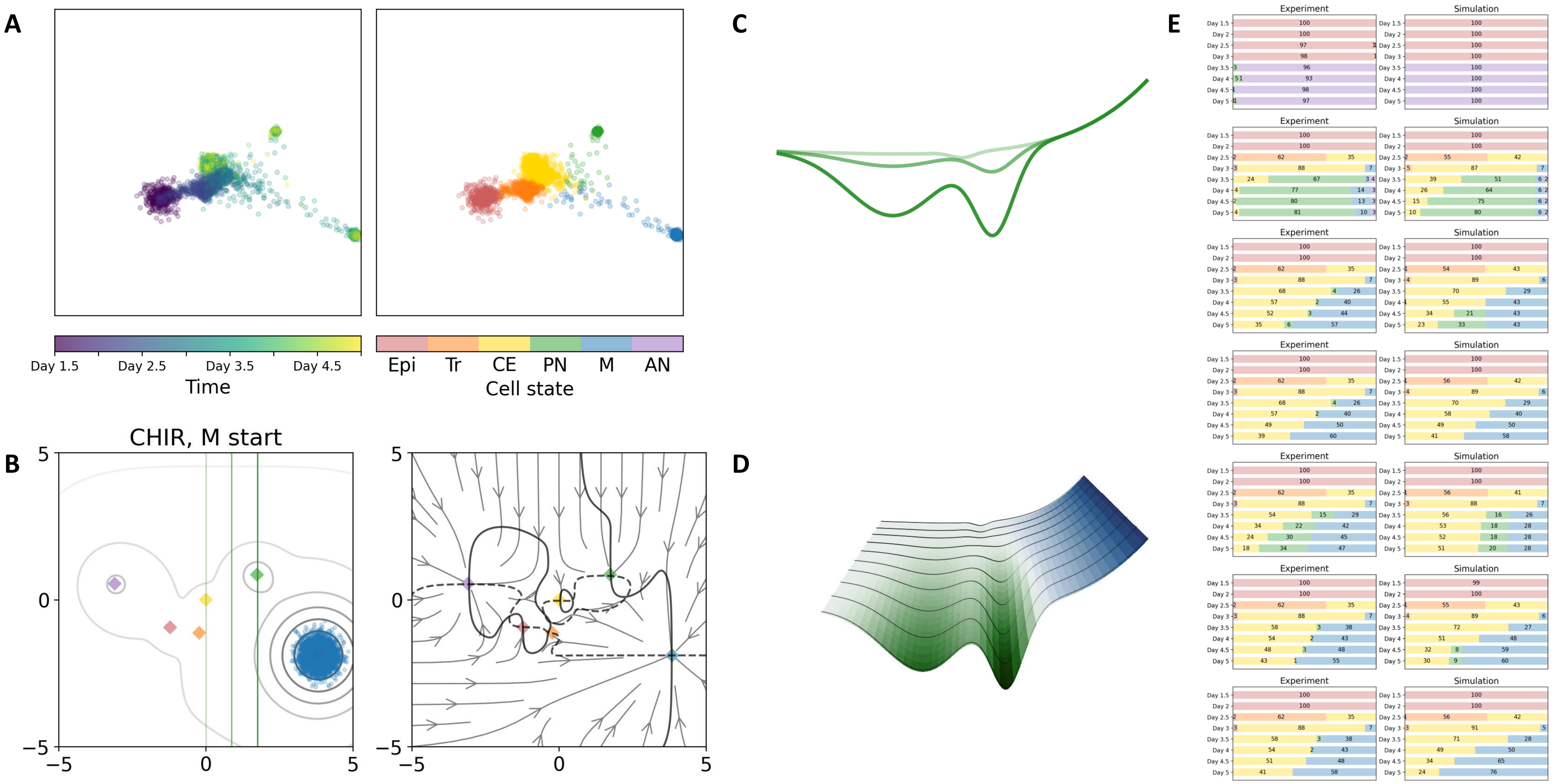

### 08 — Timecode: 20250531-031025

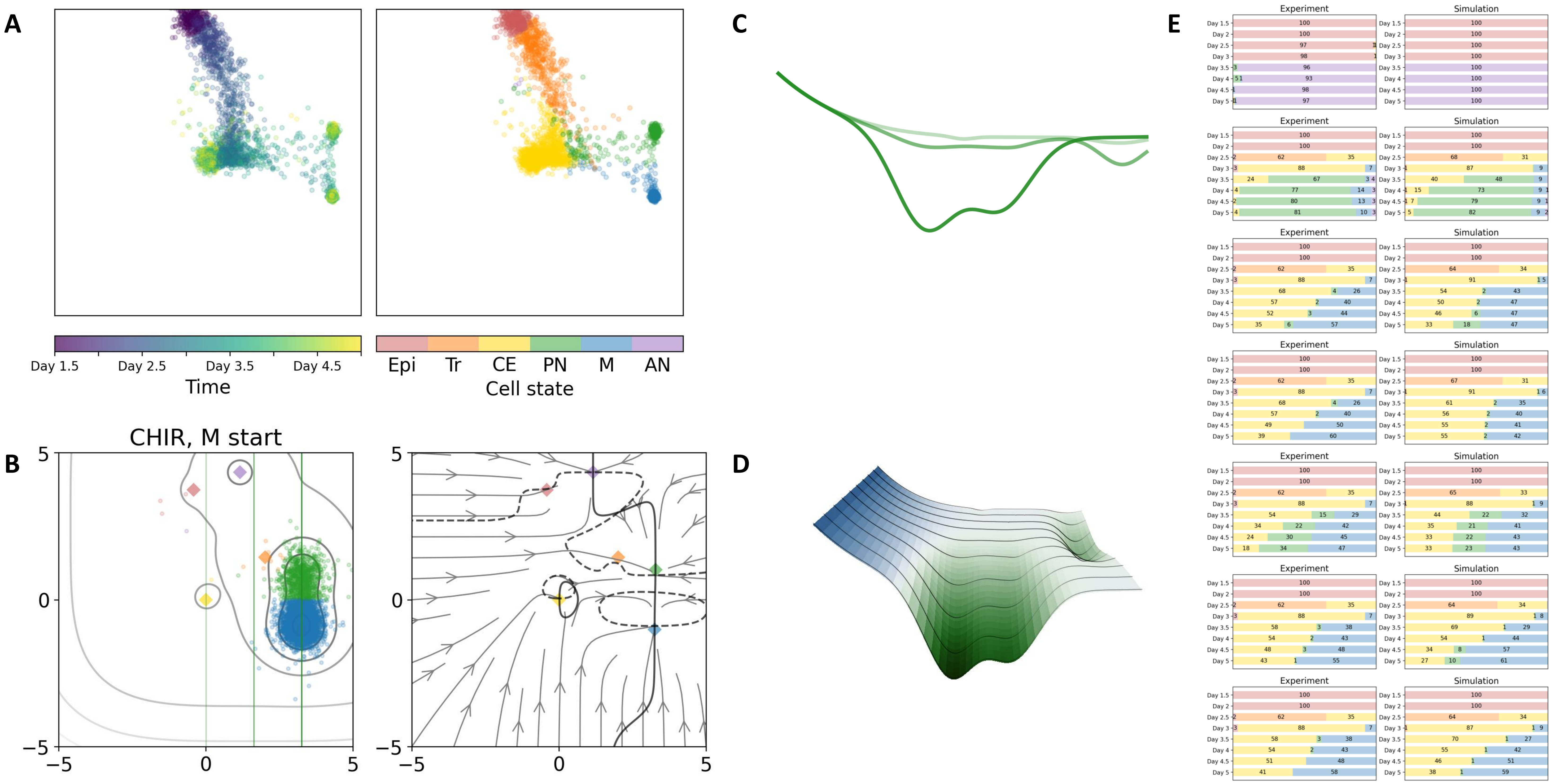

### 18 — Timecode: 20250531-121053

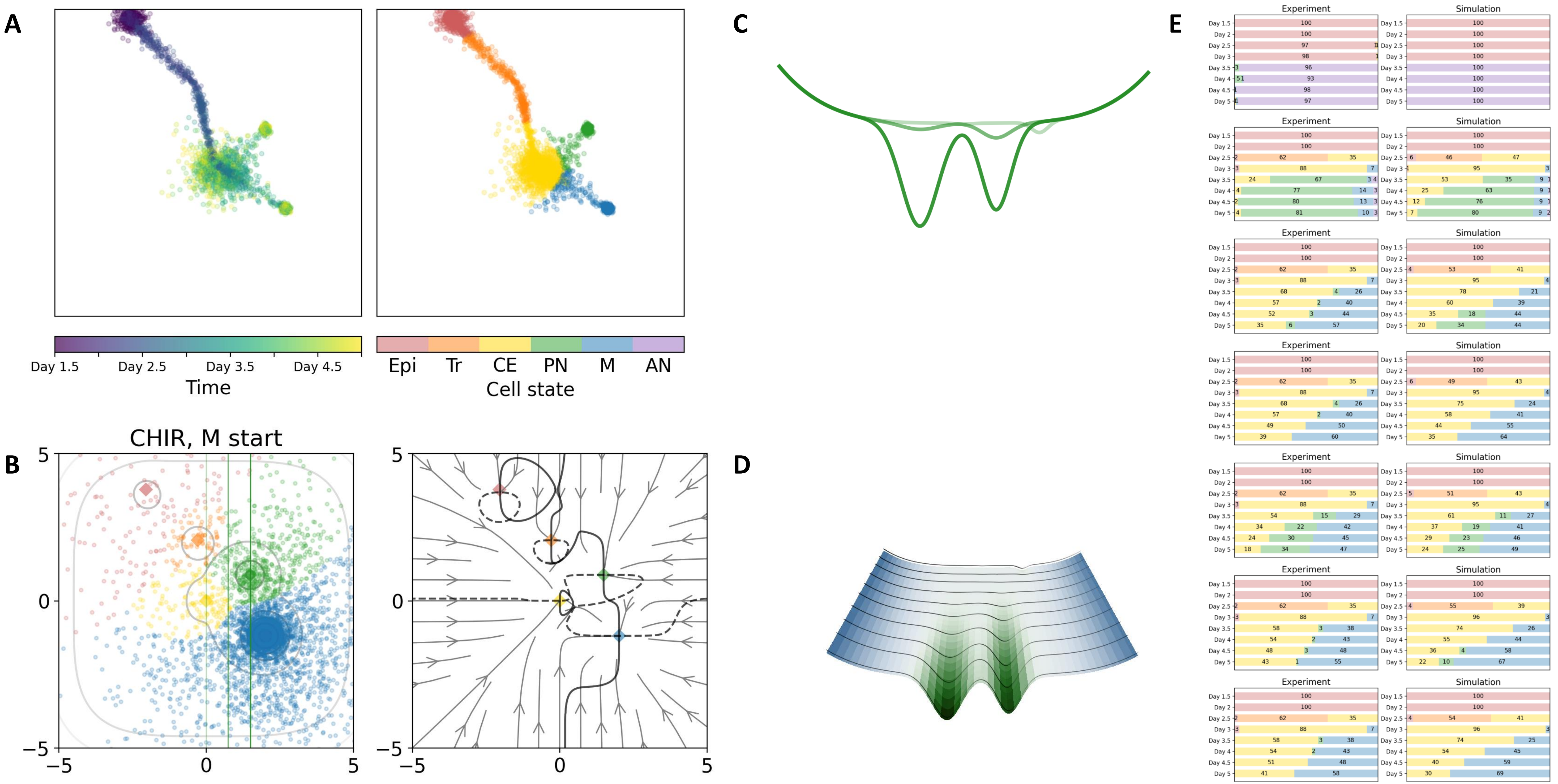

### 23 — Timecode: 20250601-000710

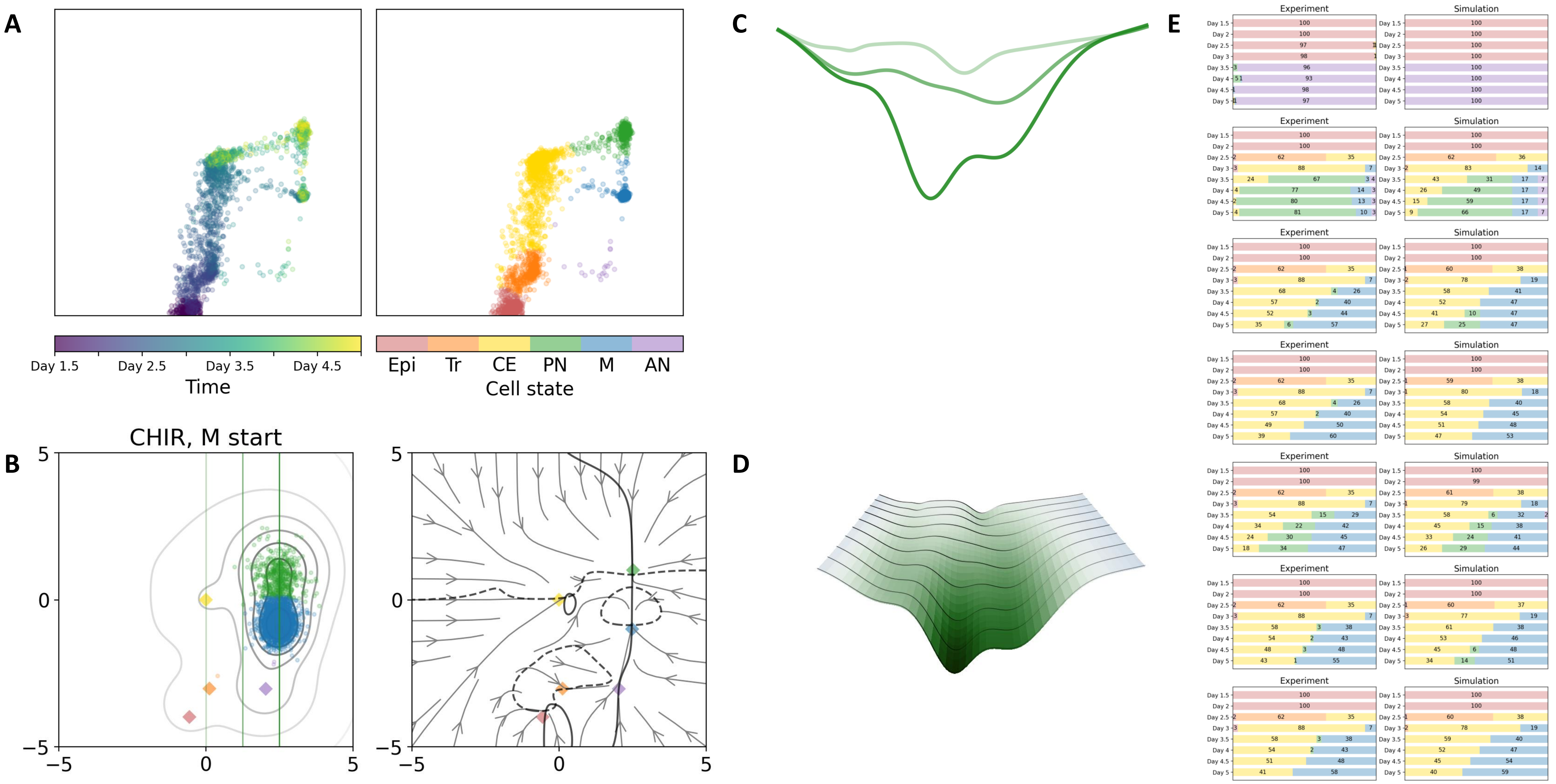

### 25 — Timecode: 20250601-010249

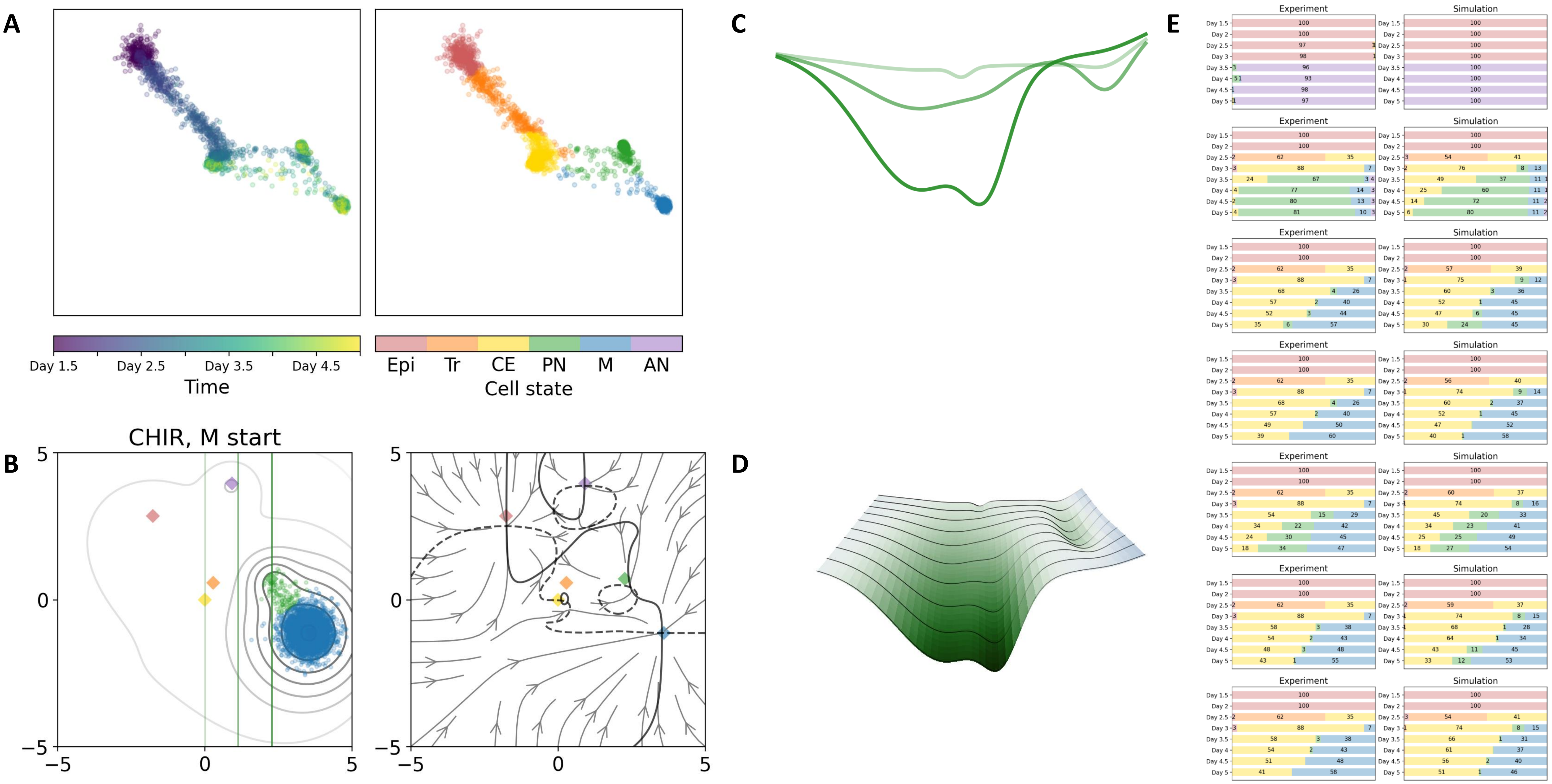

### 32 — Timecode: 20250601-073856

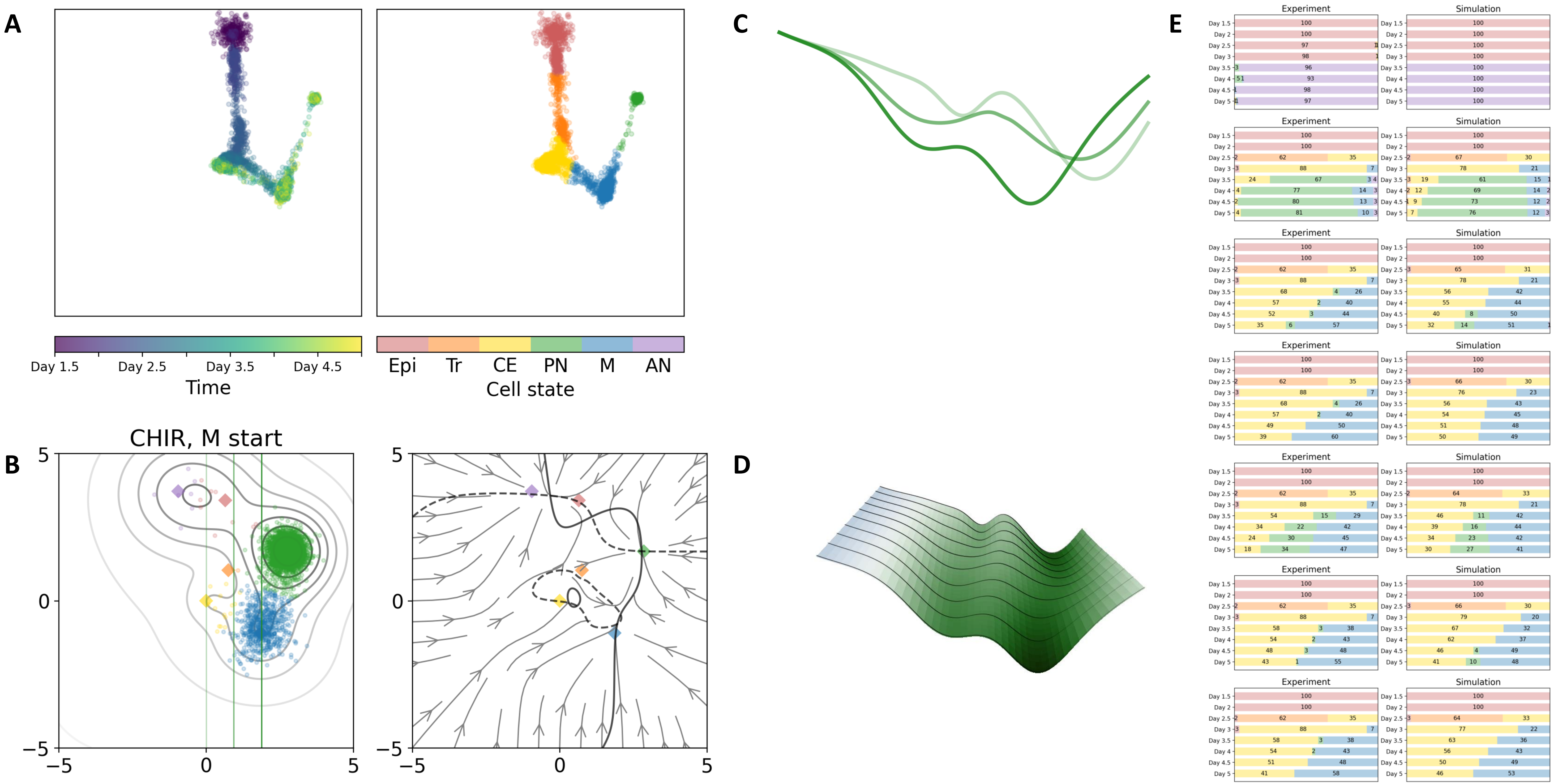

### 34 — Timecode: 20250601-093439

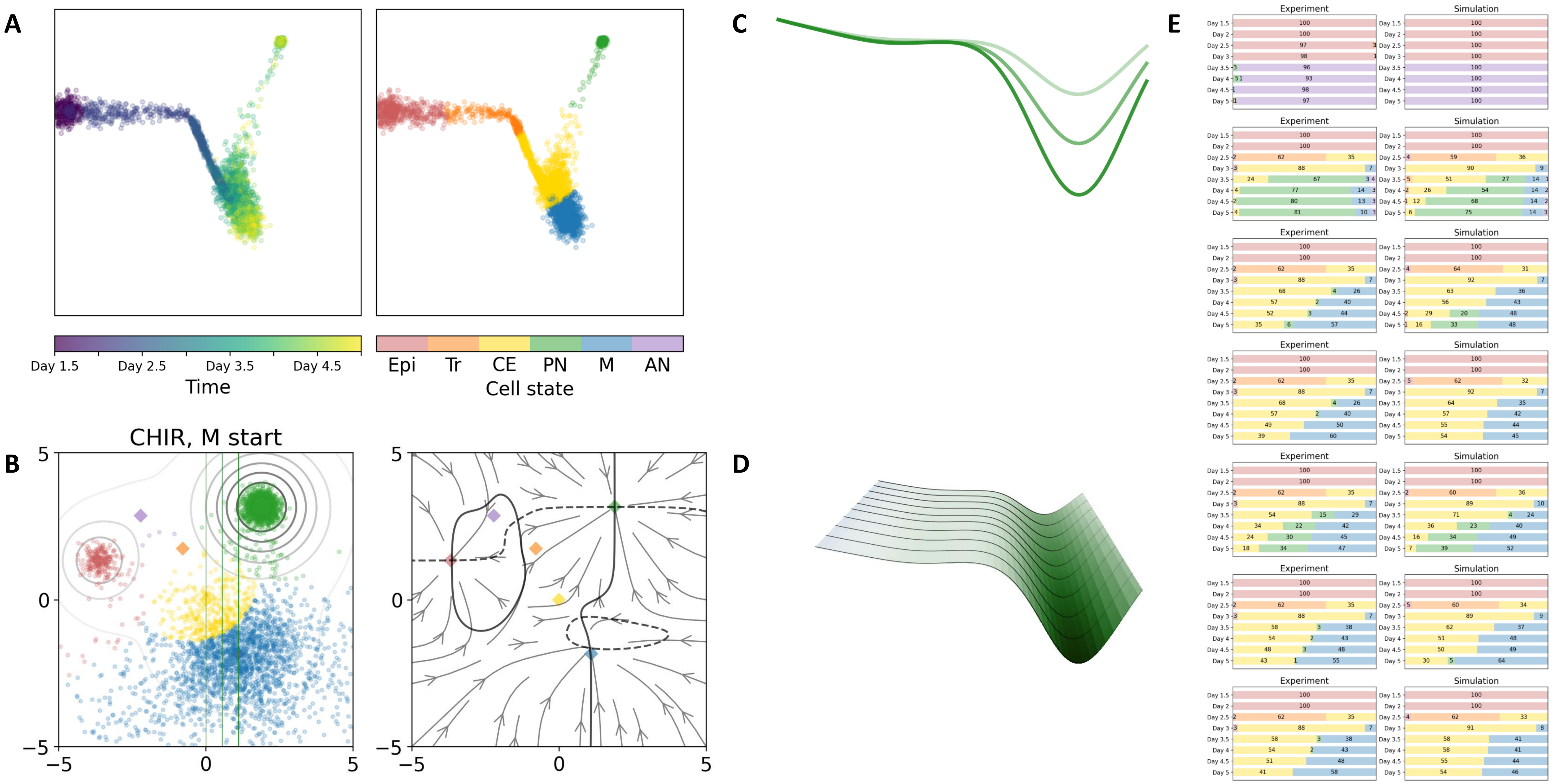

### 36 — Timecode: 20250601-134422

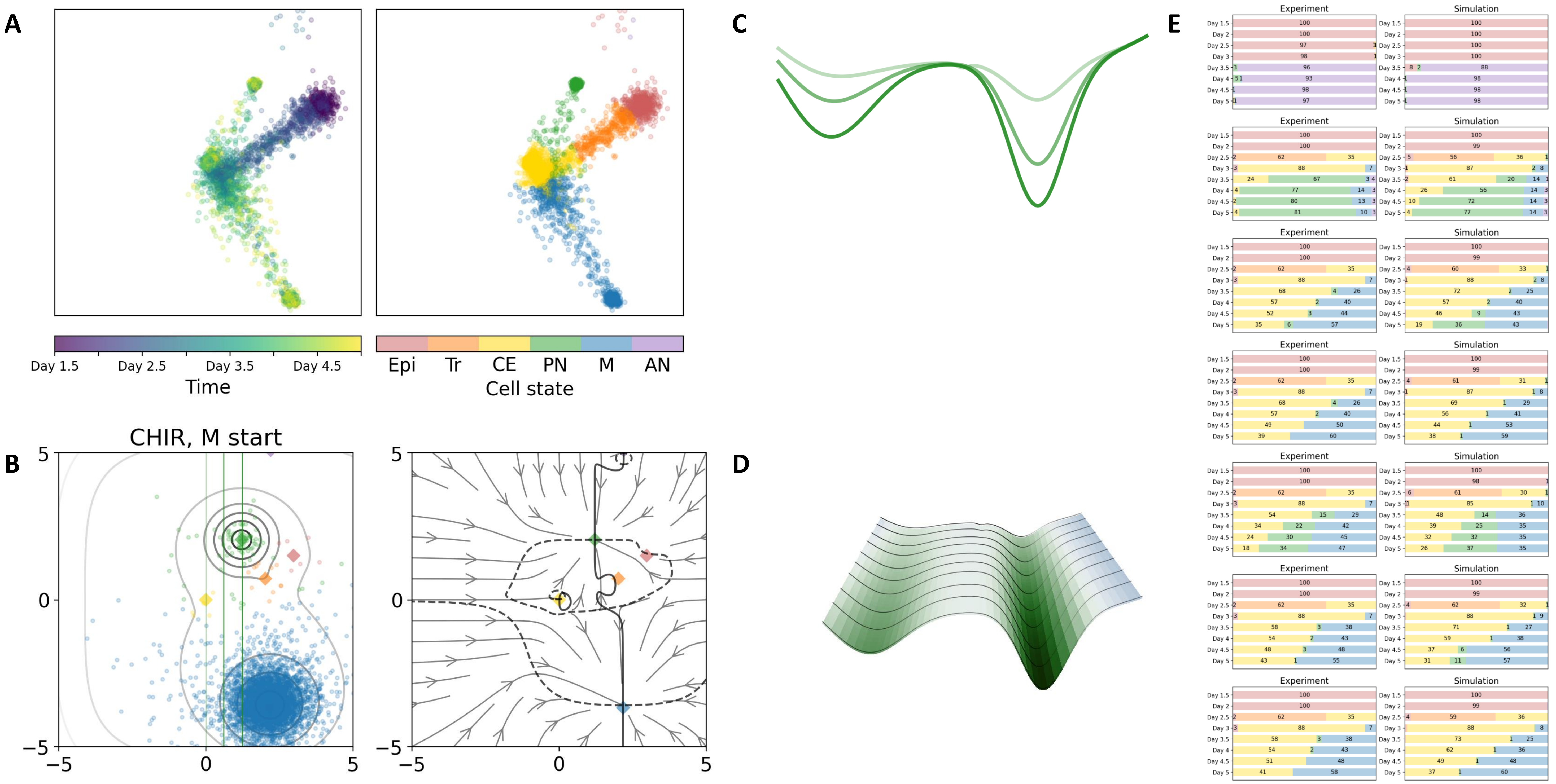

### 38 — Timecode: 20250601-151343

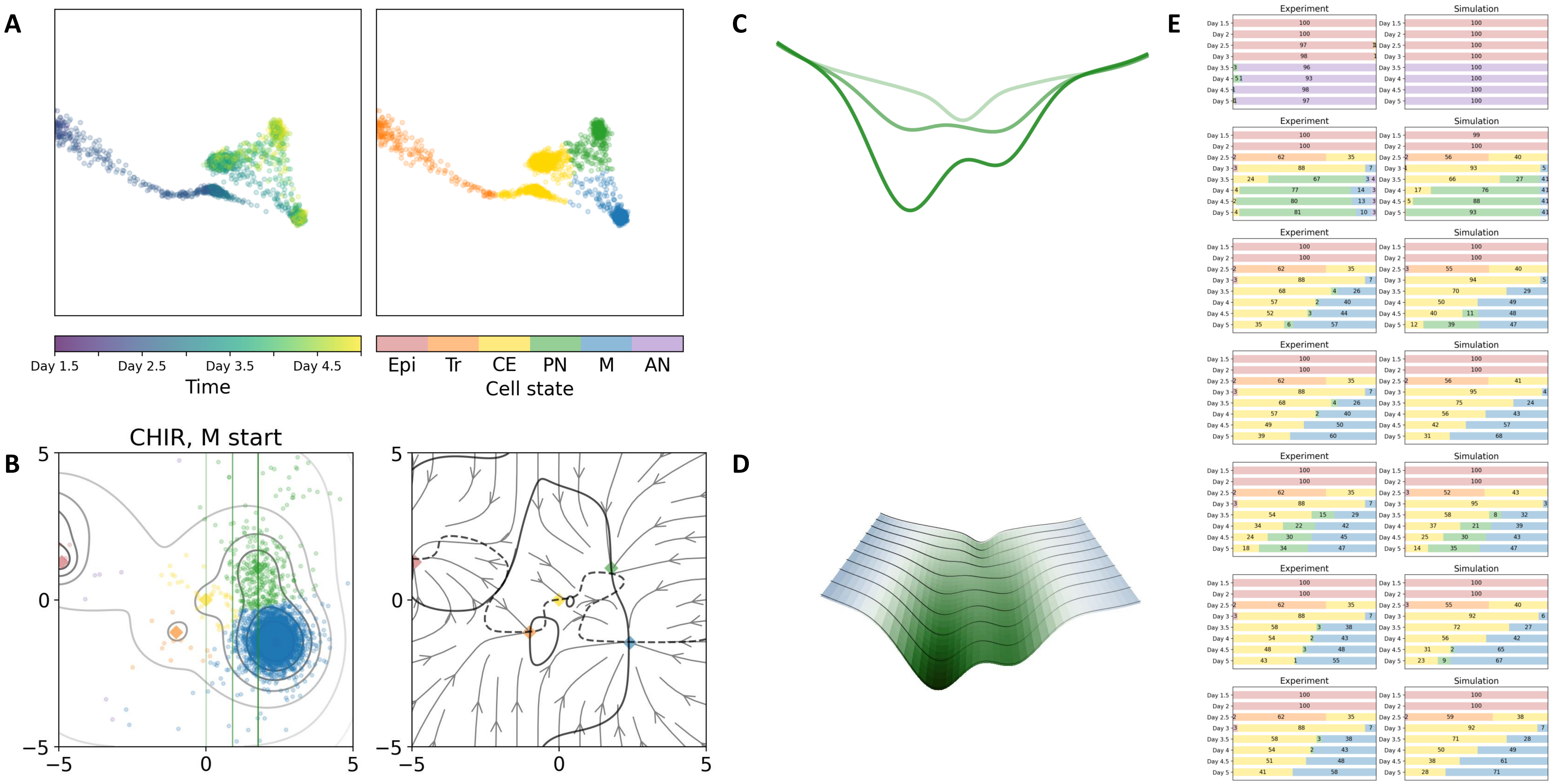

40 — Timecode: 20250601-160738

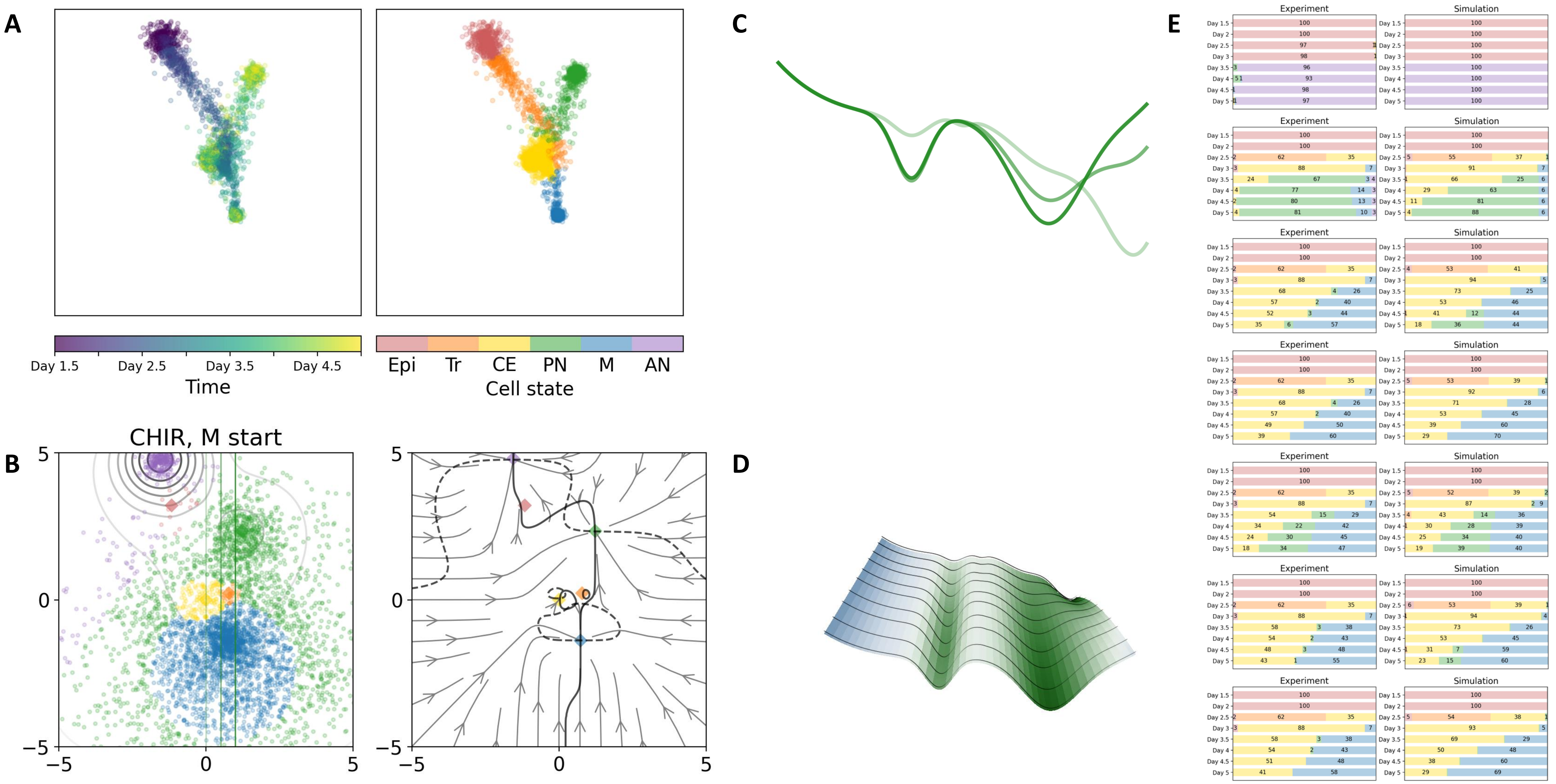

### 41 — Timecode: 20250601-162538

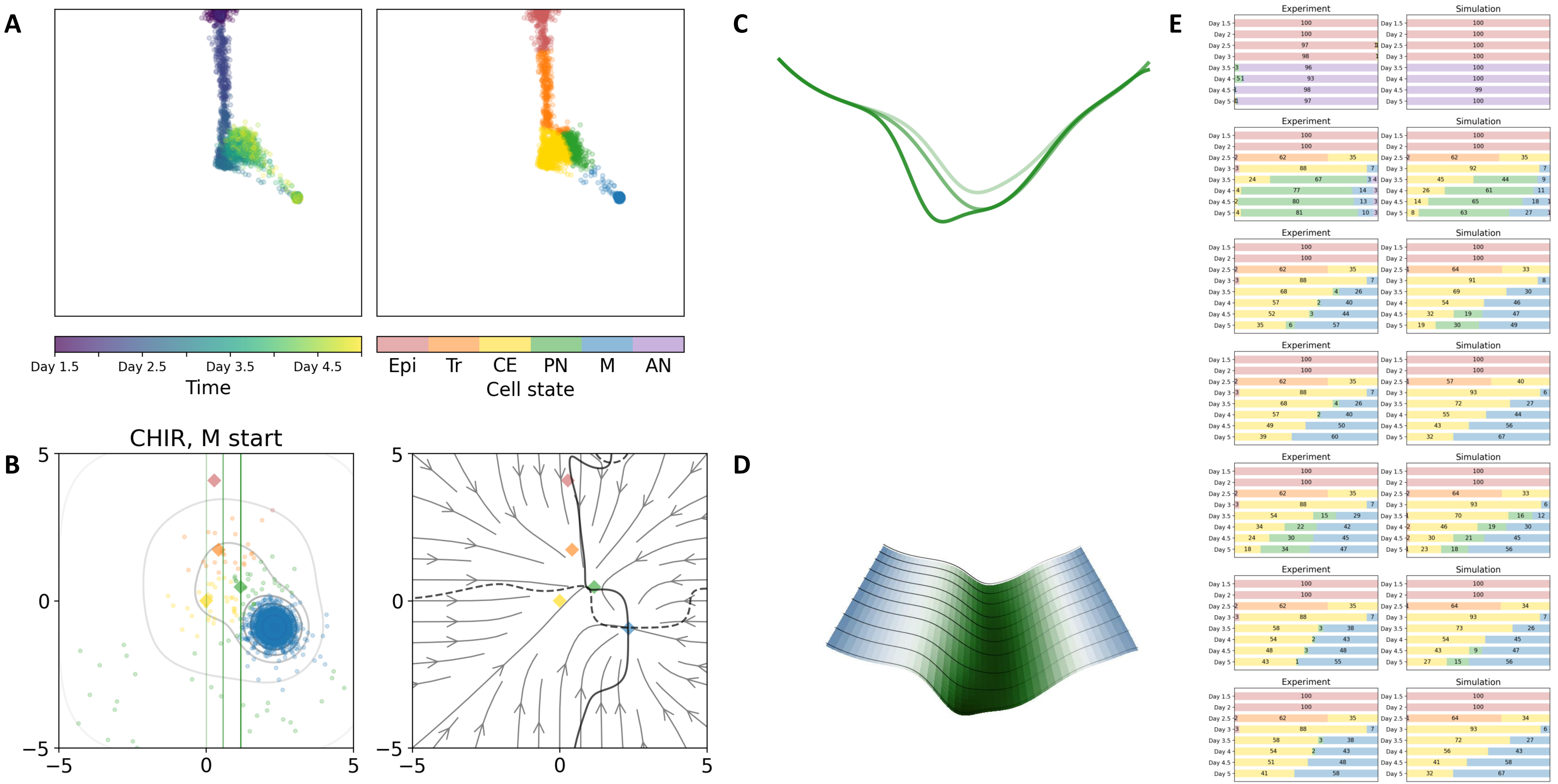

### 42 — Timecode: 20250601-204405

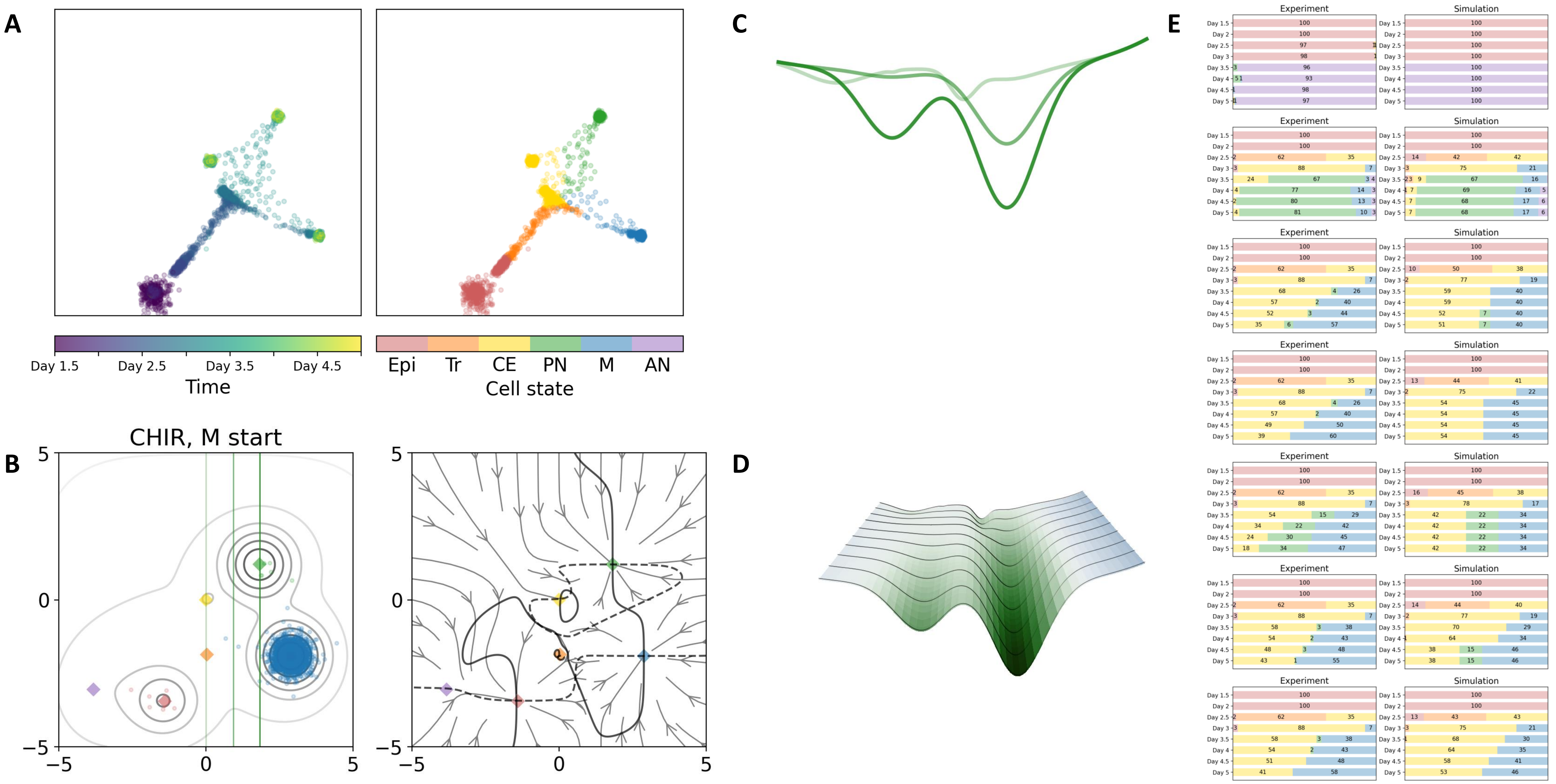

### 45 — Timecode: 20250602-031622

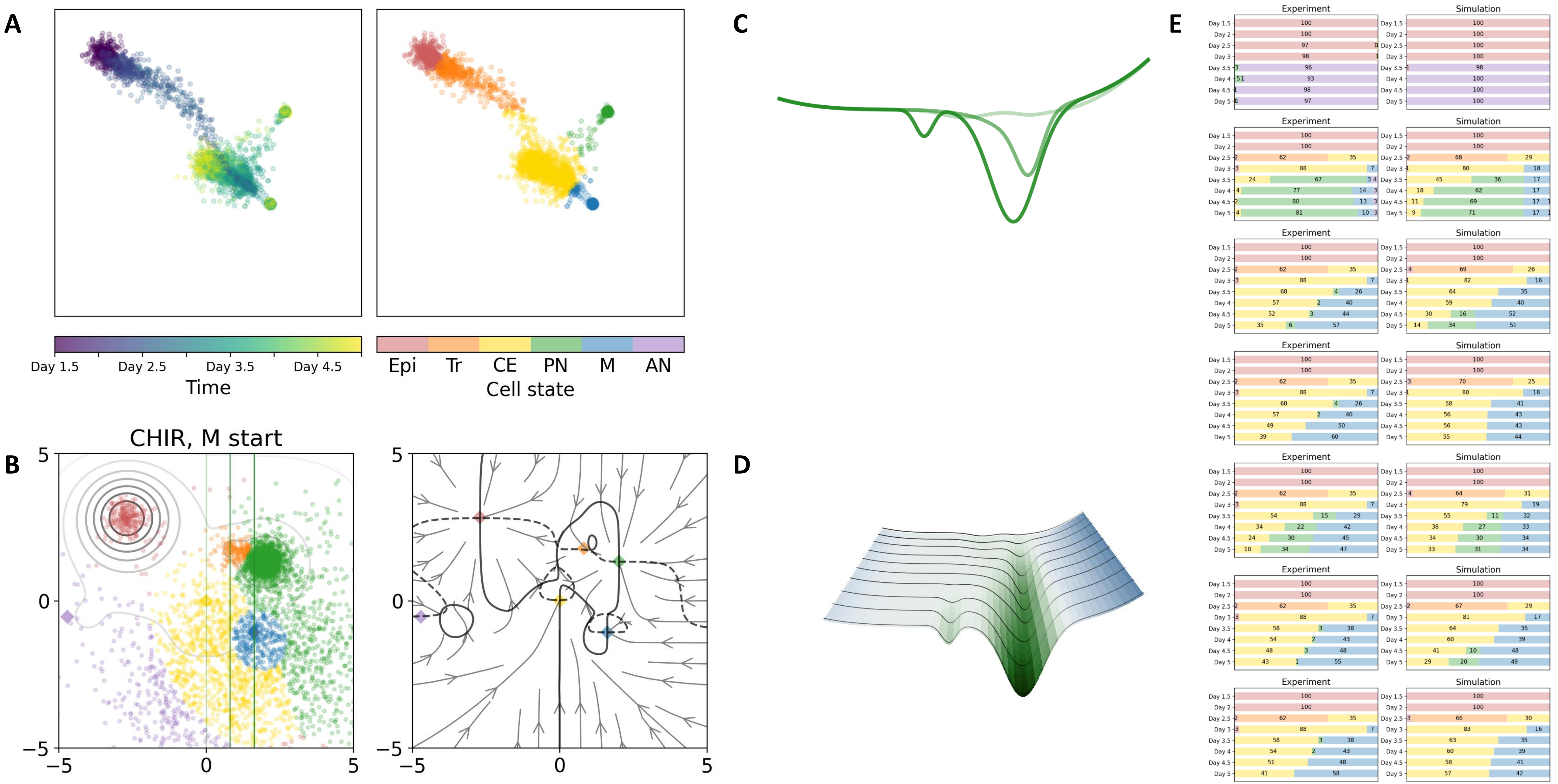

### 48 — Timecode: 20250602-062023

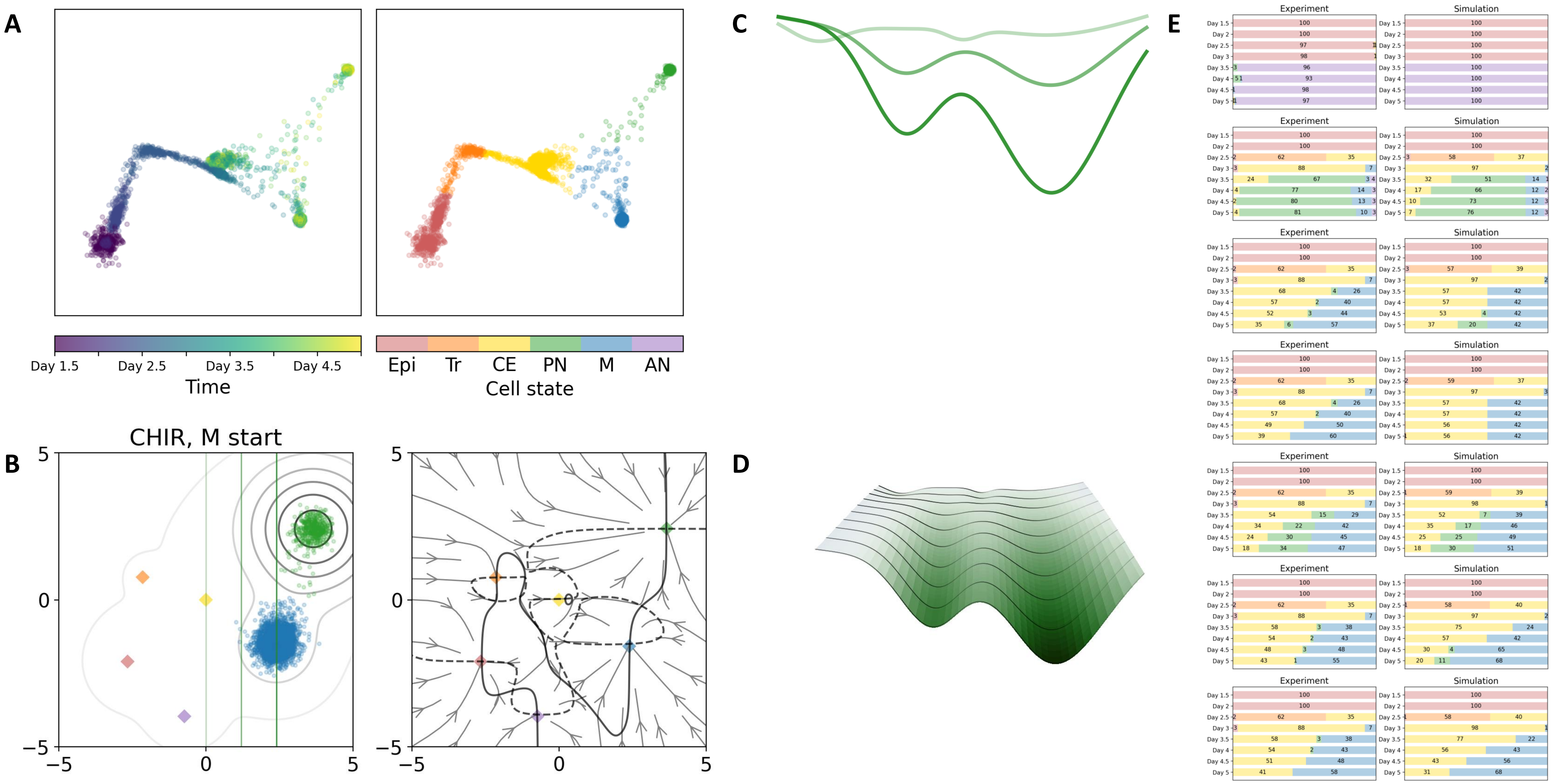

51 — Timecode: 20250602-073629

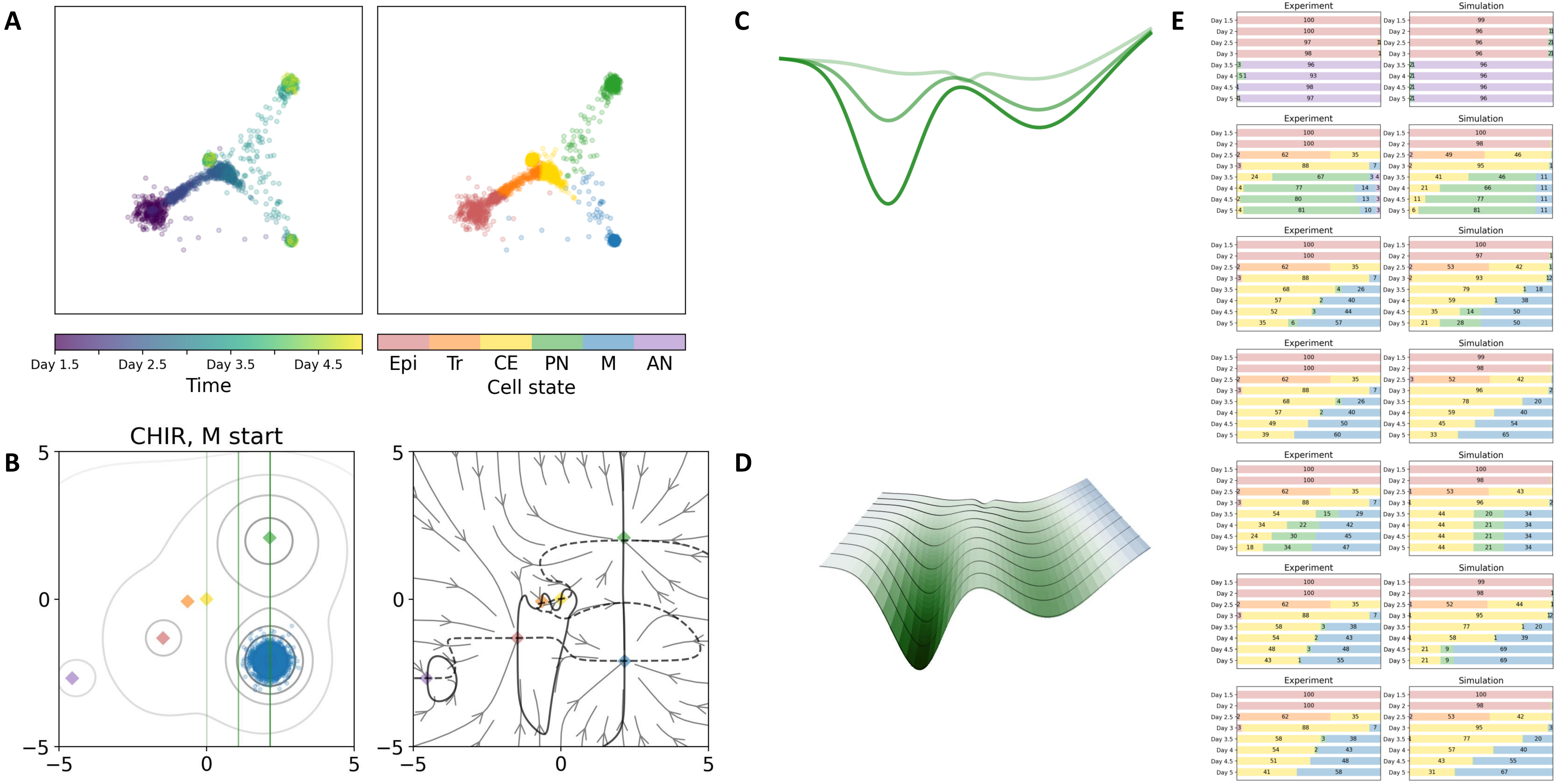

### 56 — Timecode: 20250602-133728

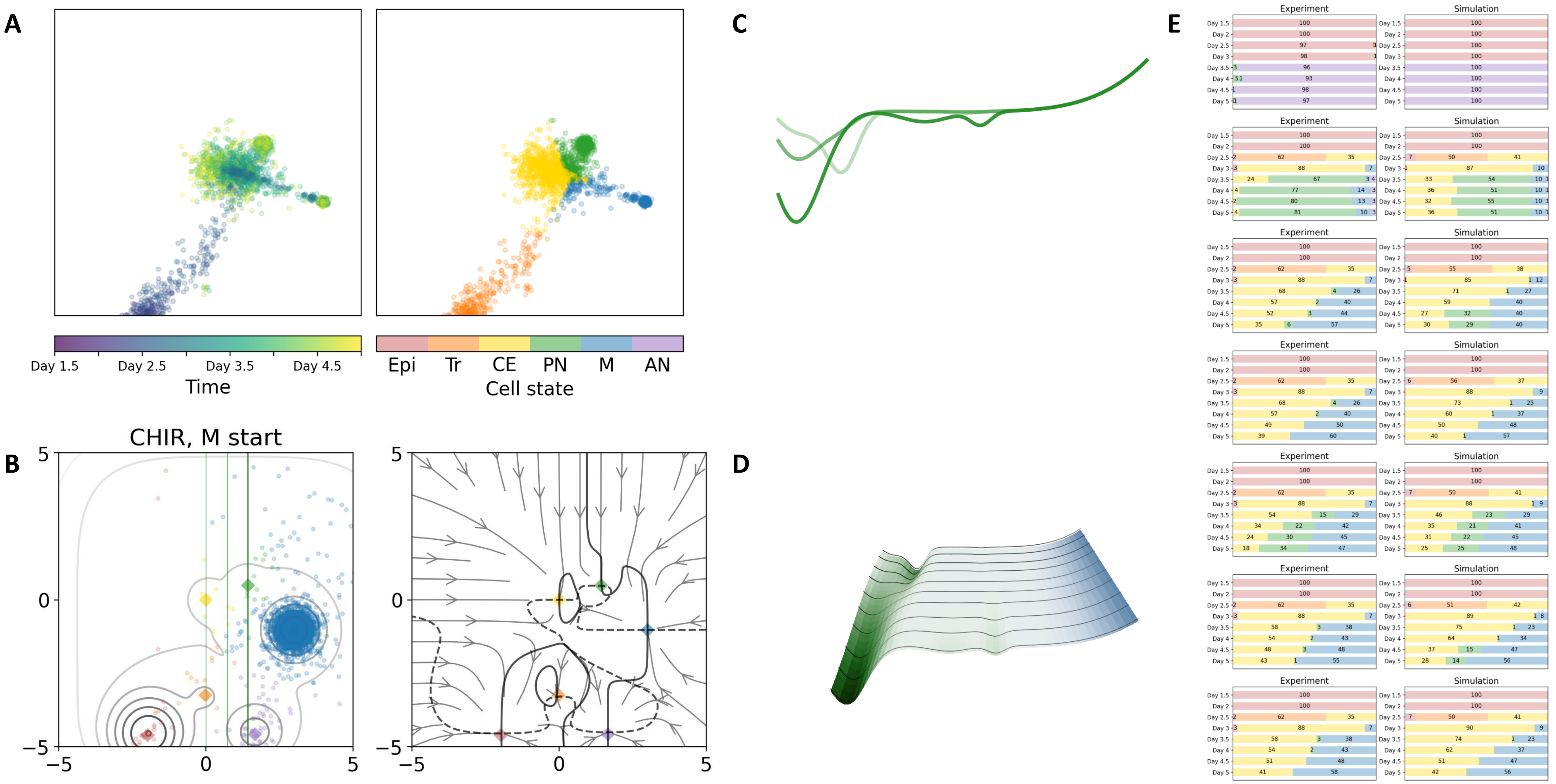

59 — Timecode: 20250602-172416

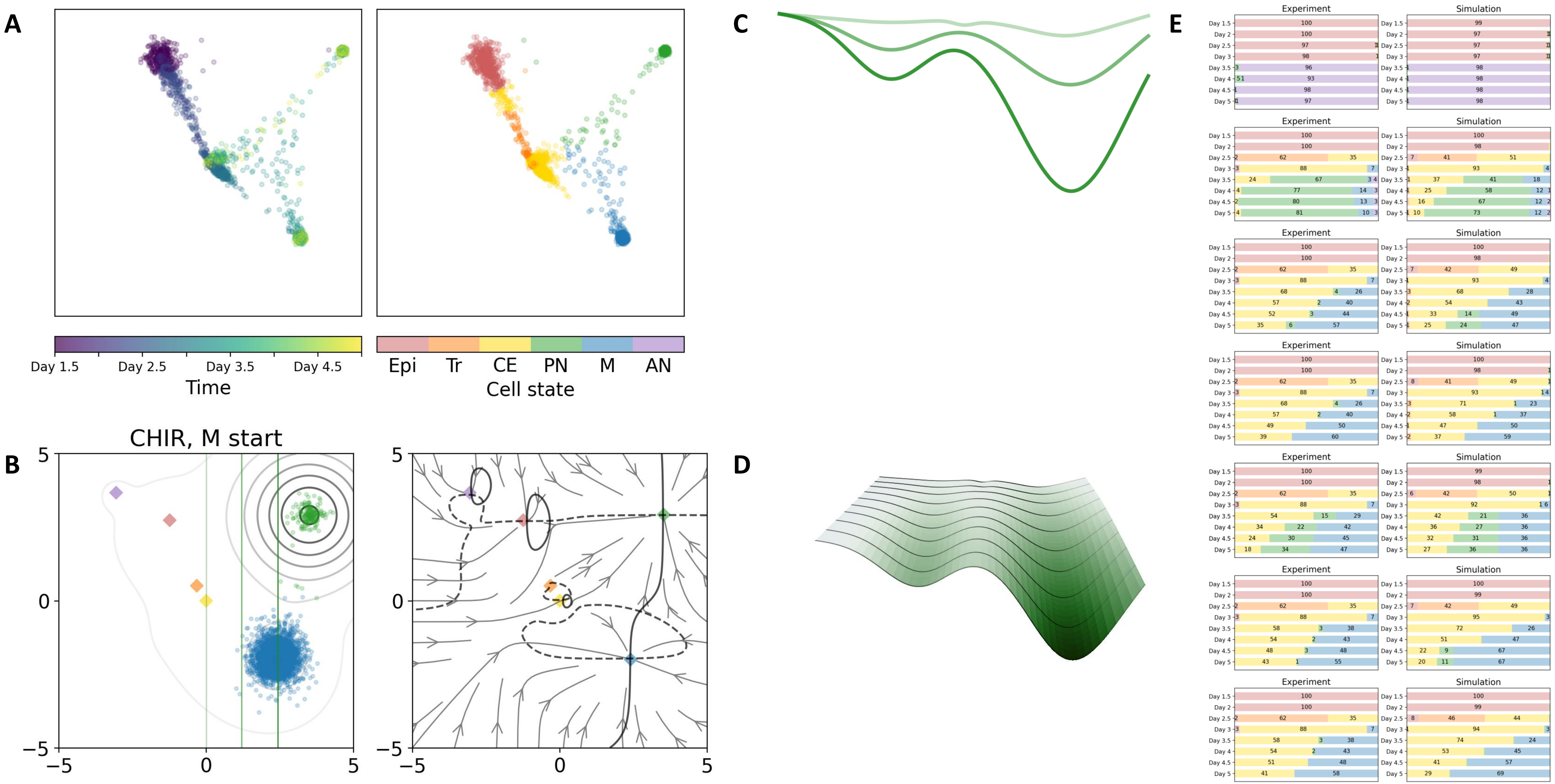

### Heteroclinic flip with overlap

### 11 — Timecode: 20250531-050416

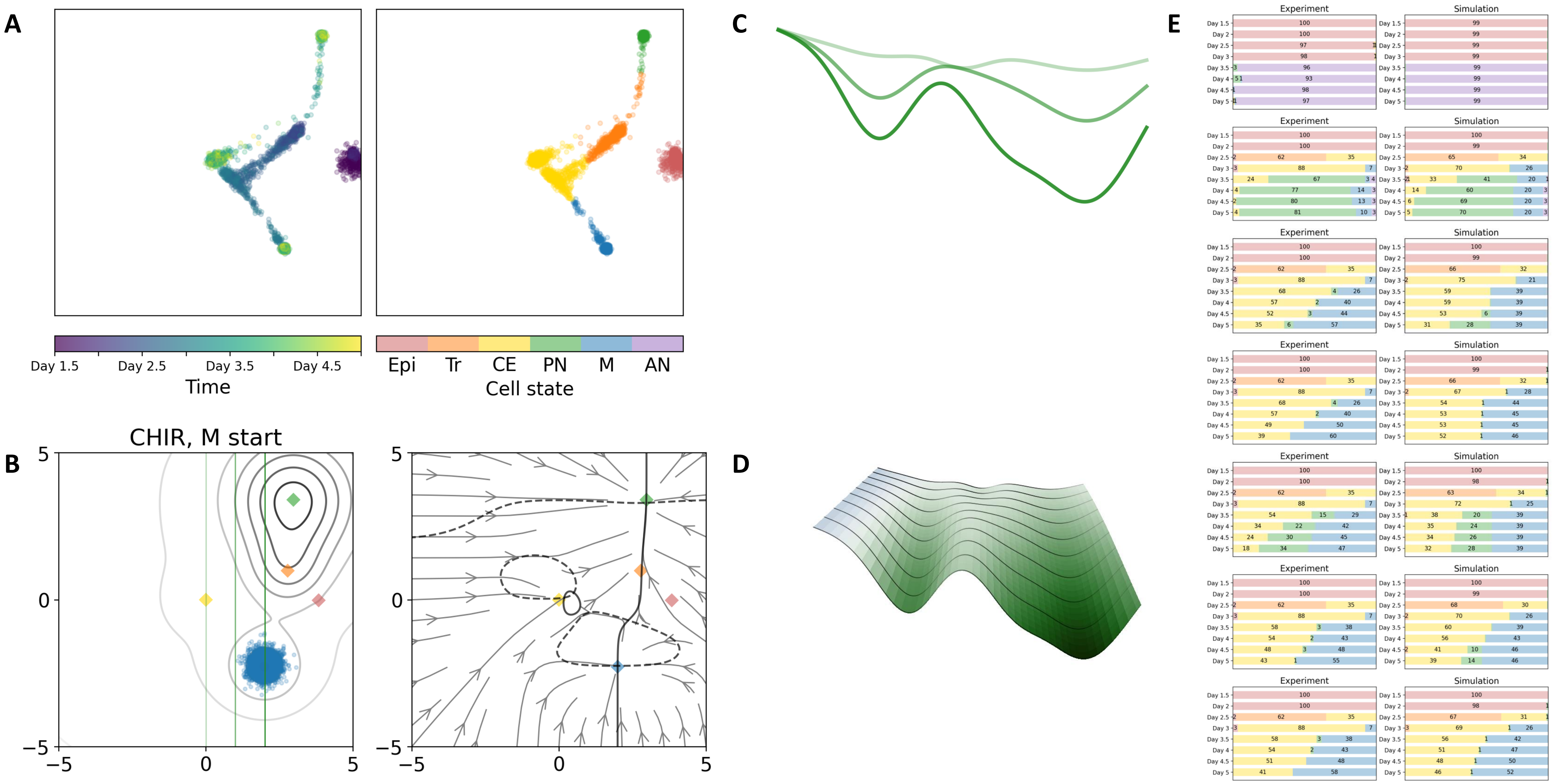

### 09 — Timecode: 20250531-040730

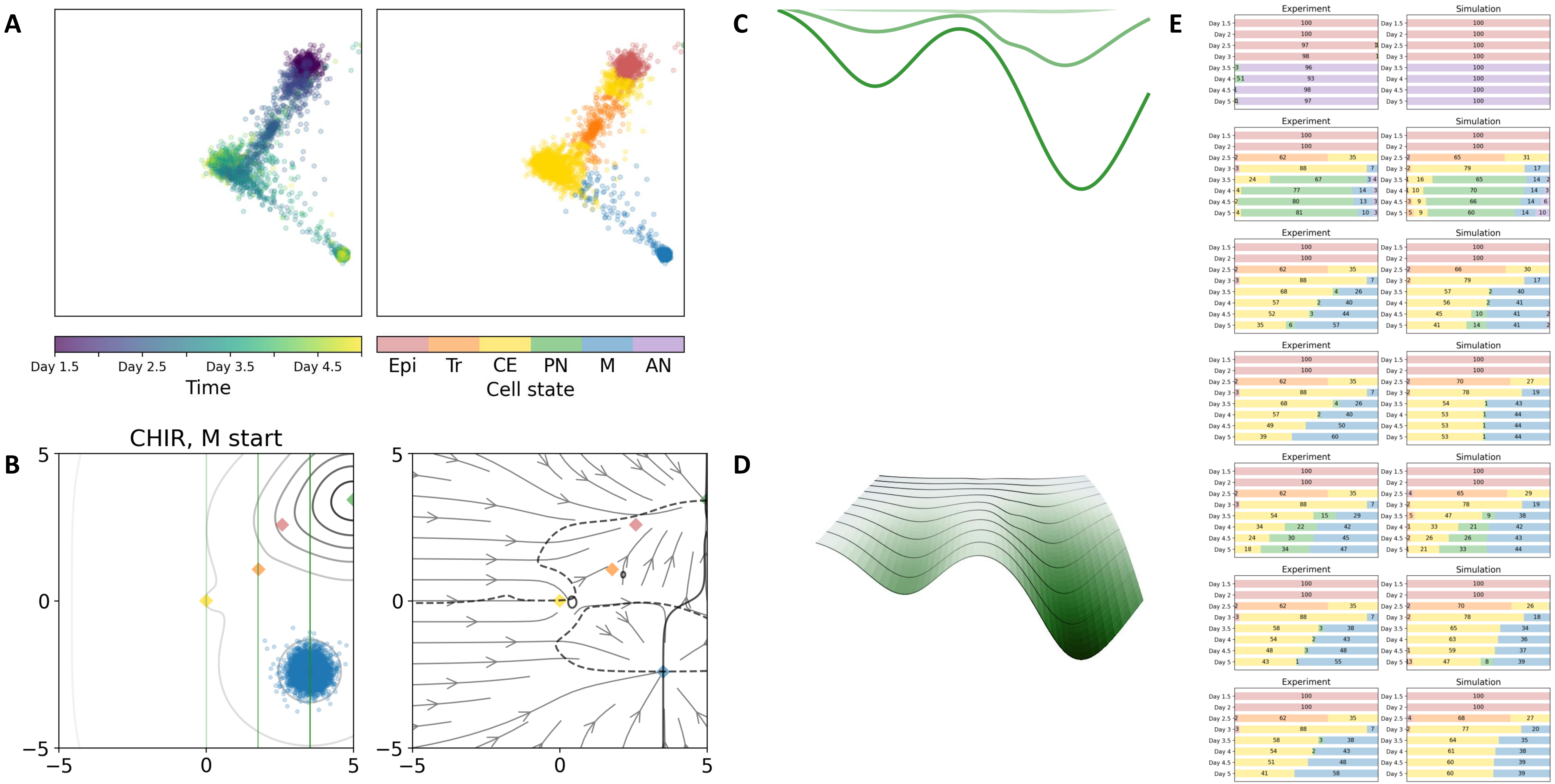

### 14 — Timecode: 20250531-073758

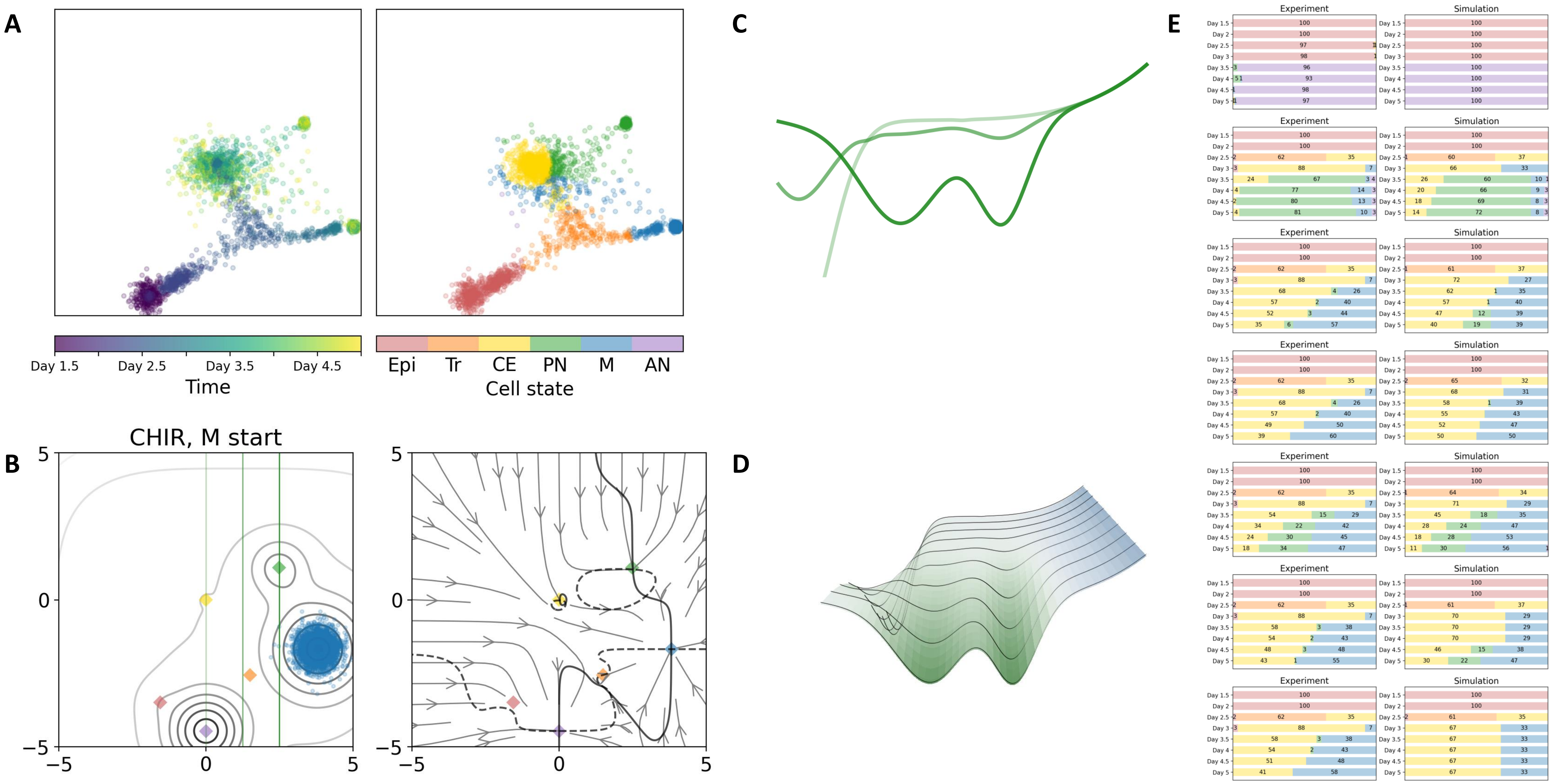

### 19 — Timecode: 20250531-122855

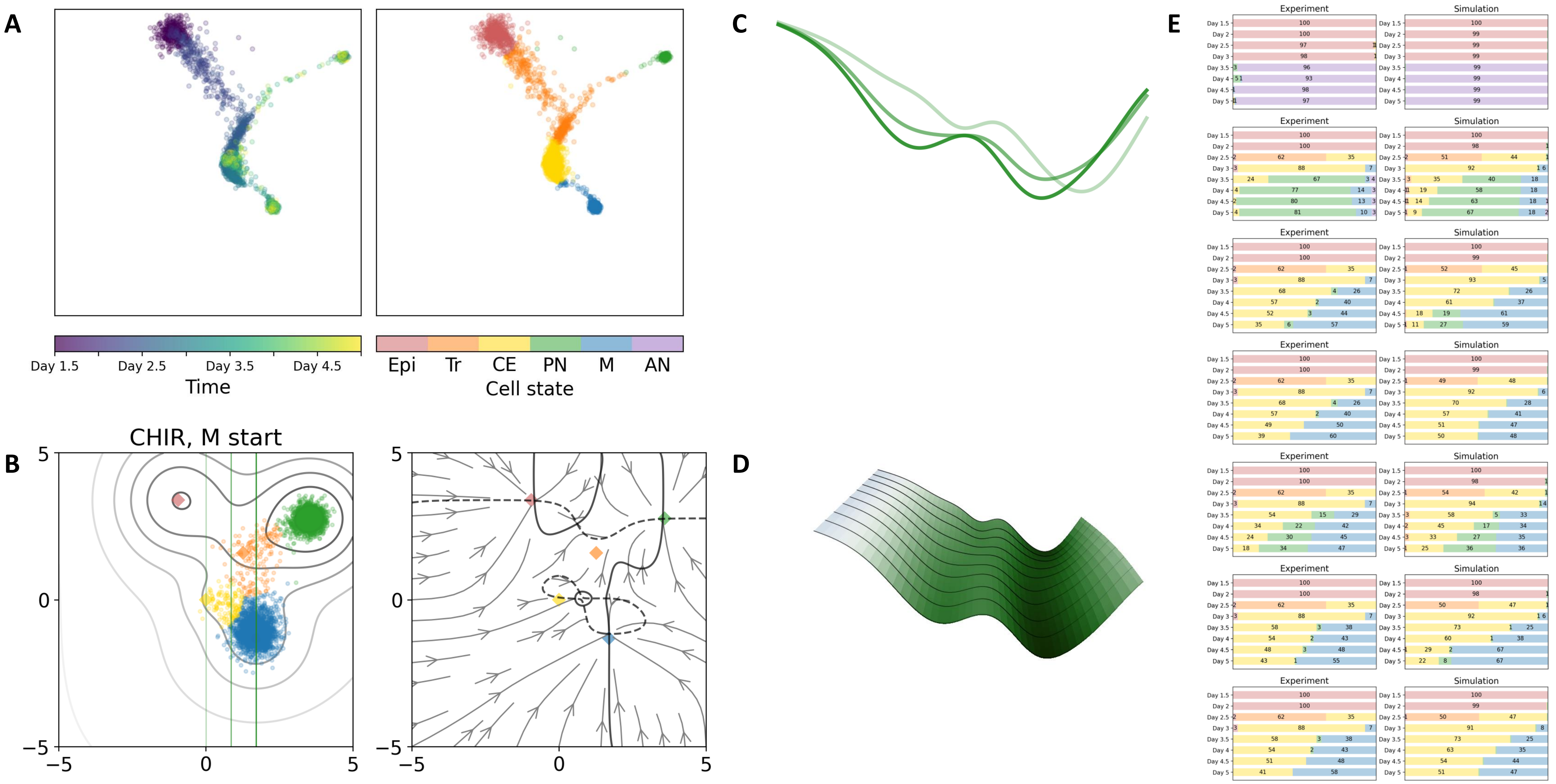

### 26 — Timecode: 20250601-015532

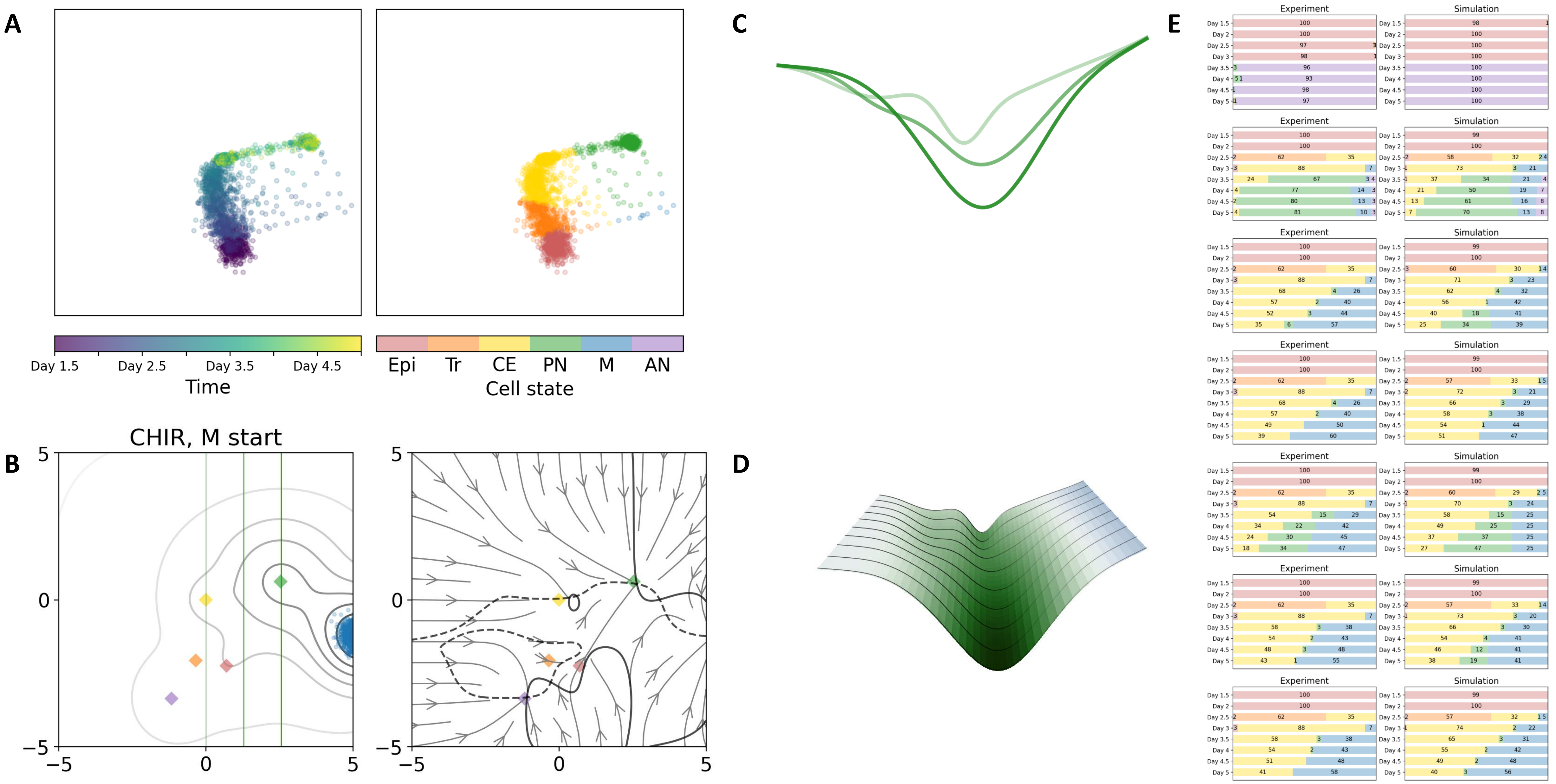

### 33 — Timecode: 20250601-091540

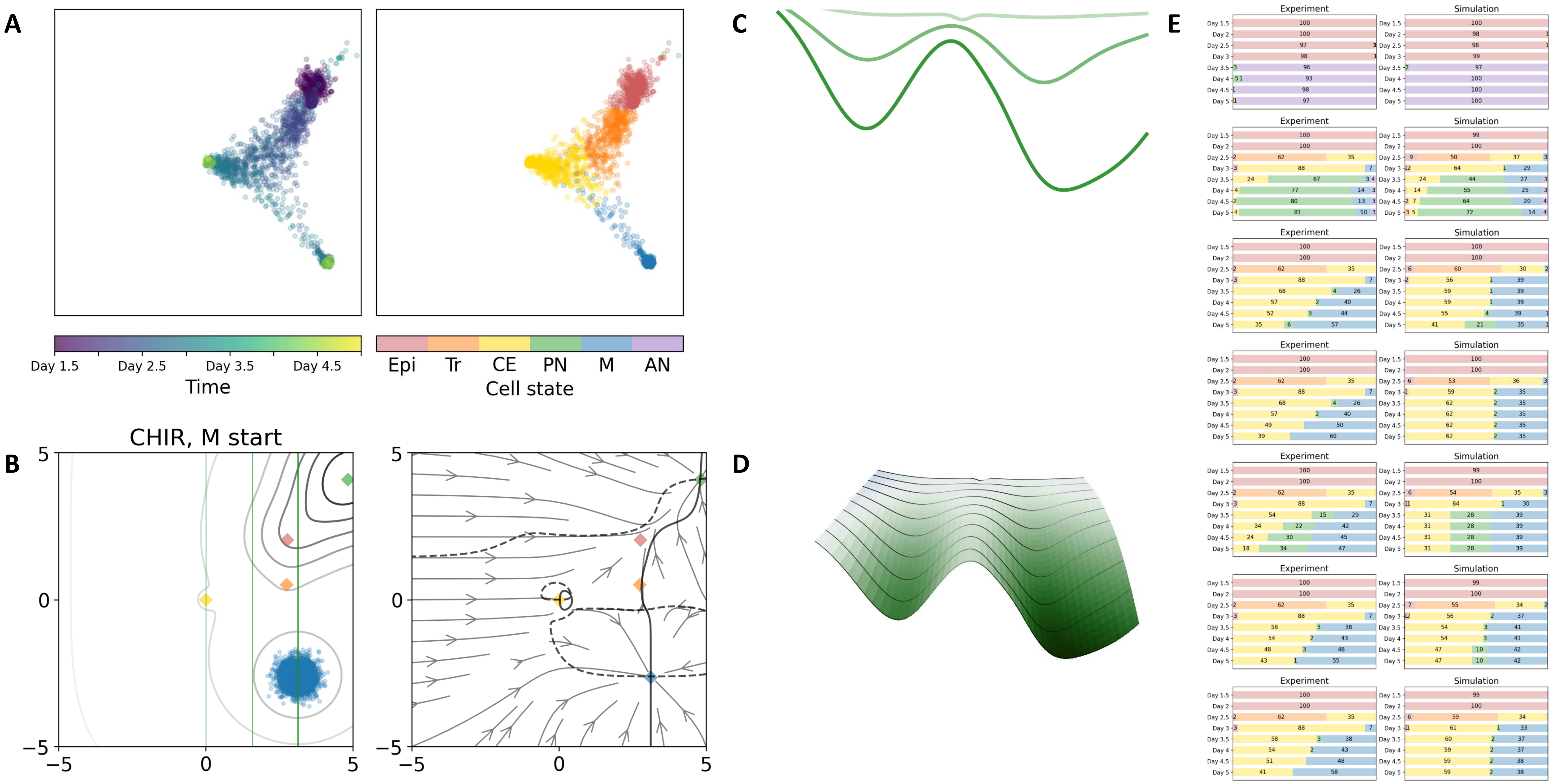

### 52 — Timecode: 20250602-075552

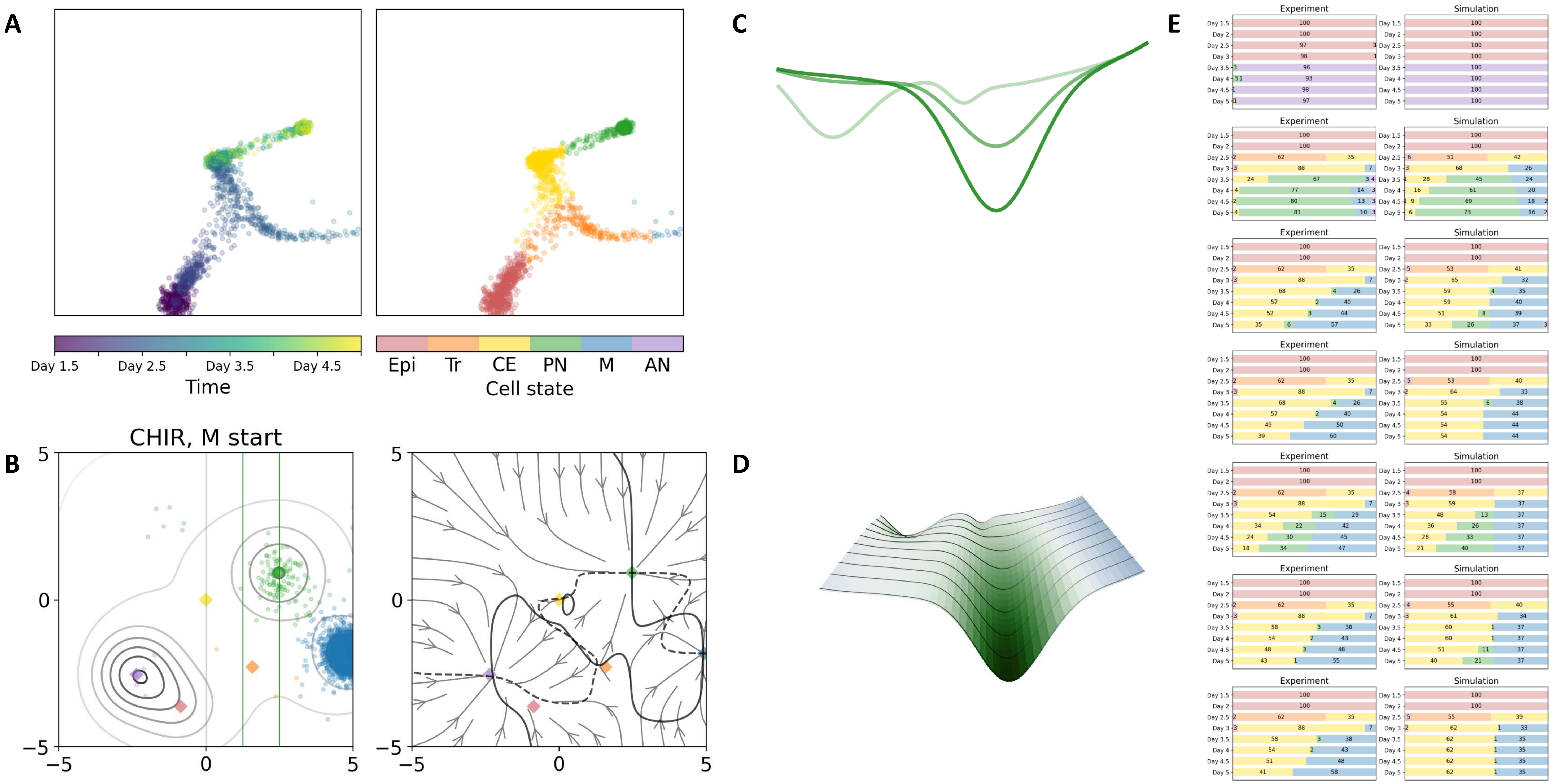

### 53 — Timecode: 20250602-091439

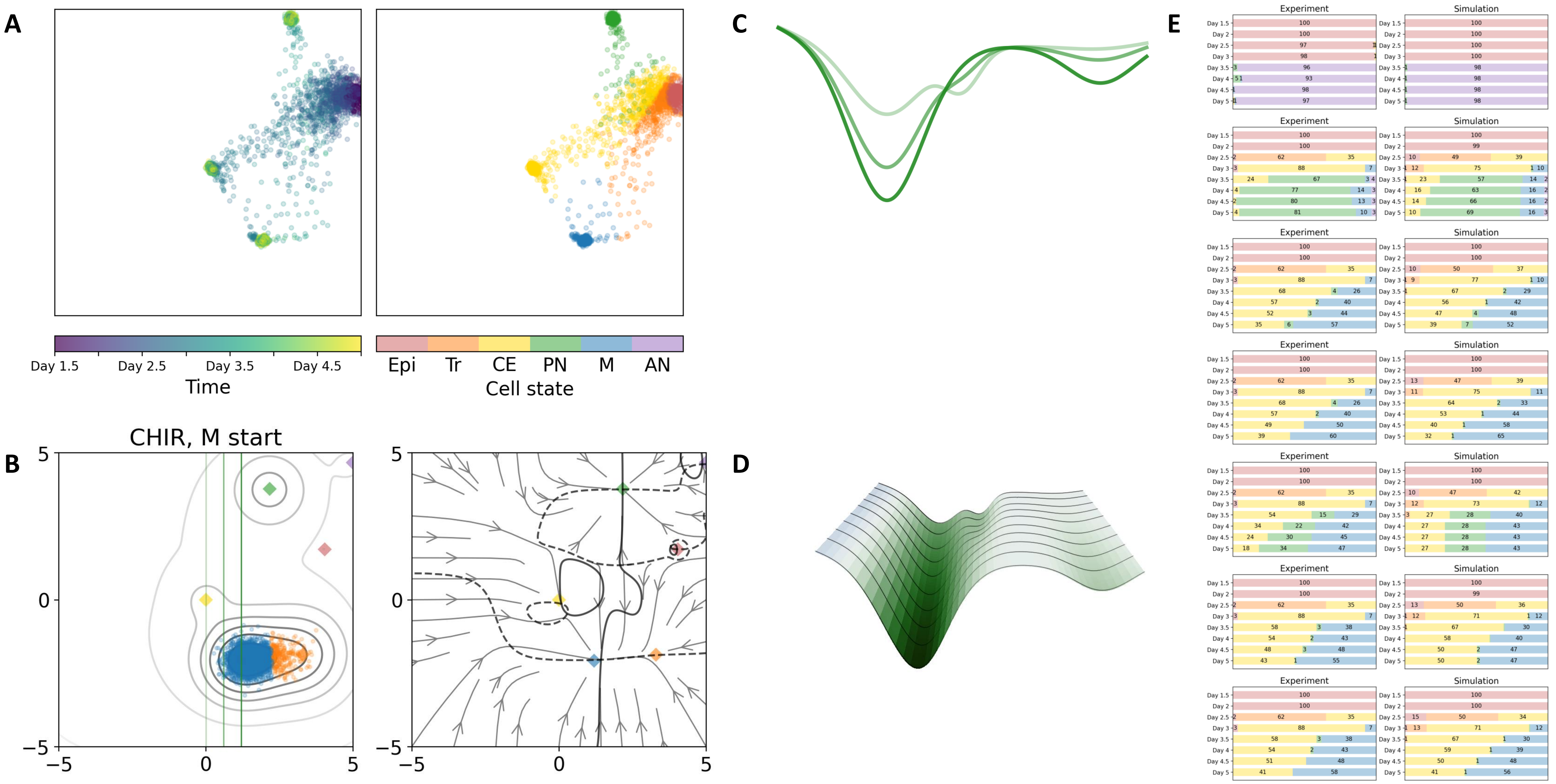

### Double cusp landscapes

### 10 — Timecode: 20250531-044417

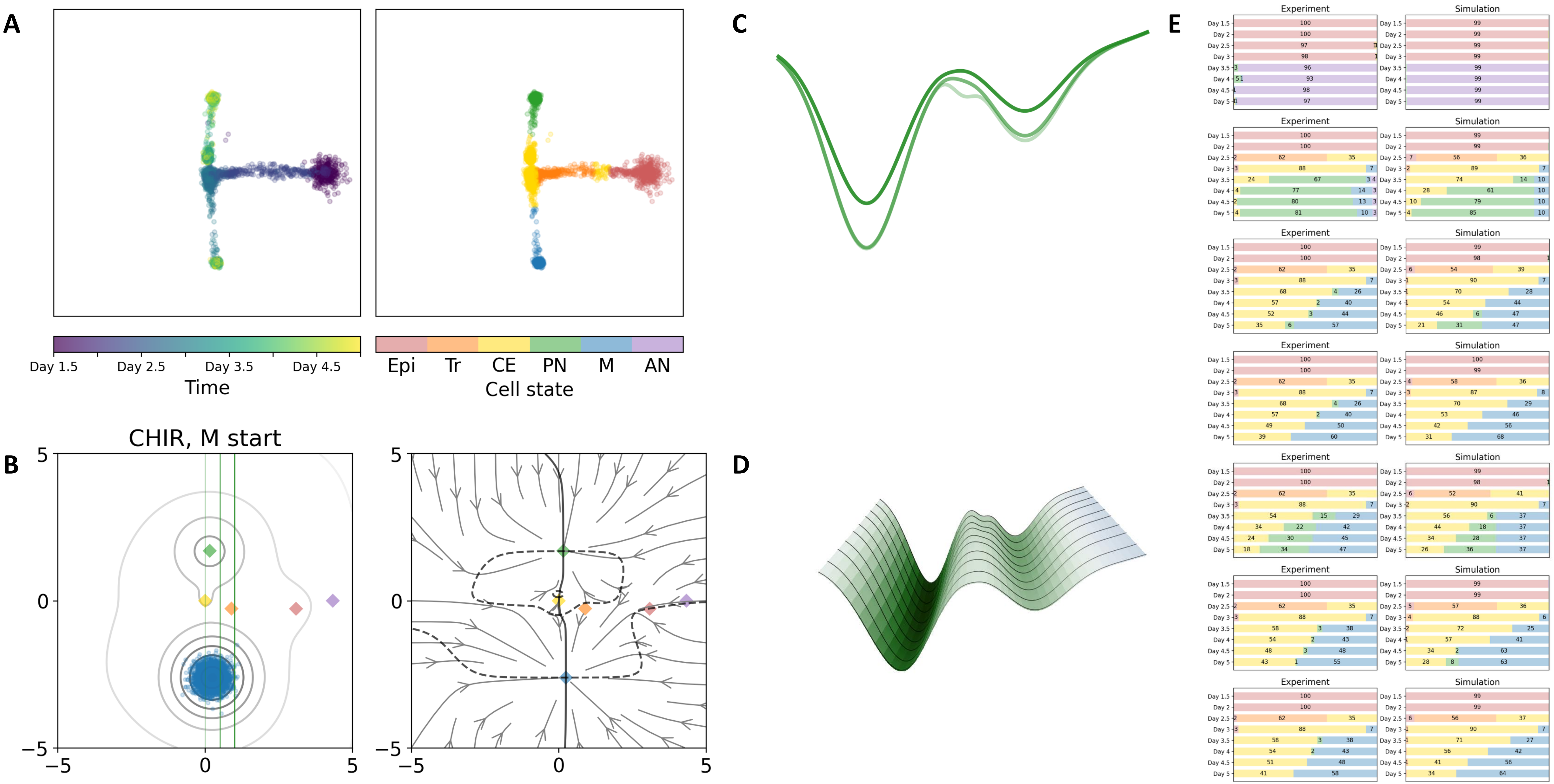

### 12 — Timecode: 20250531-060537

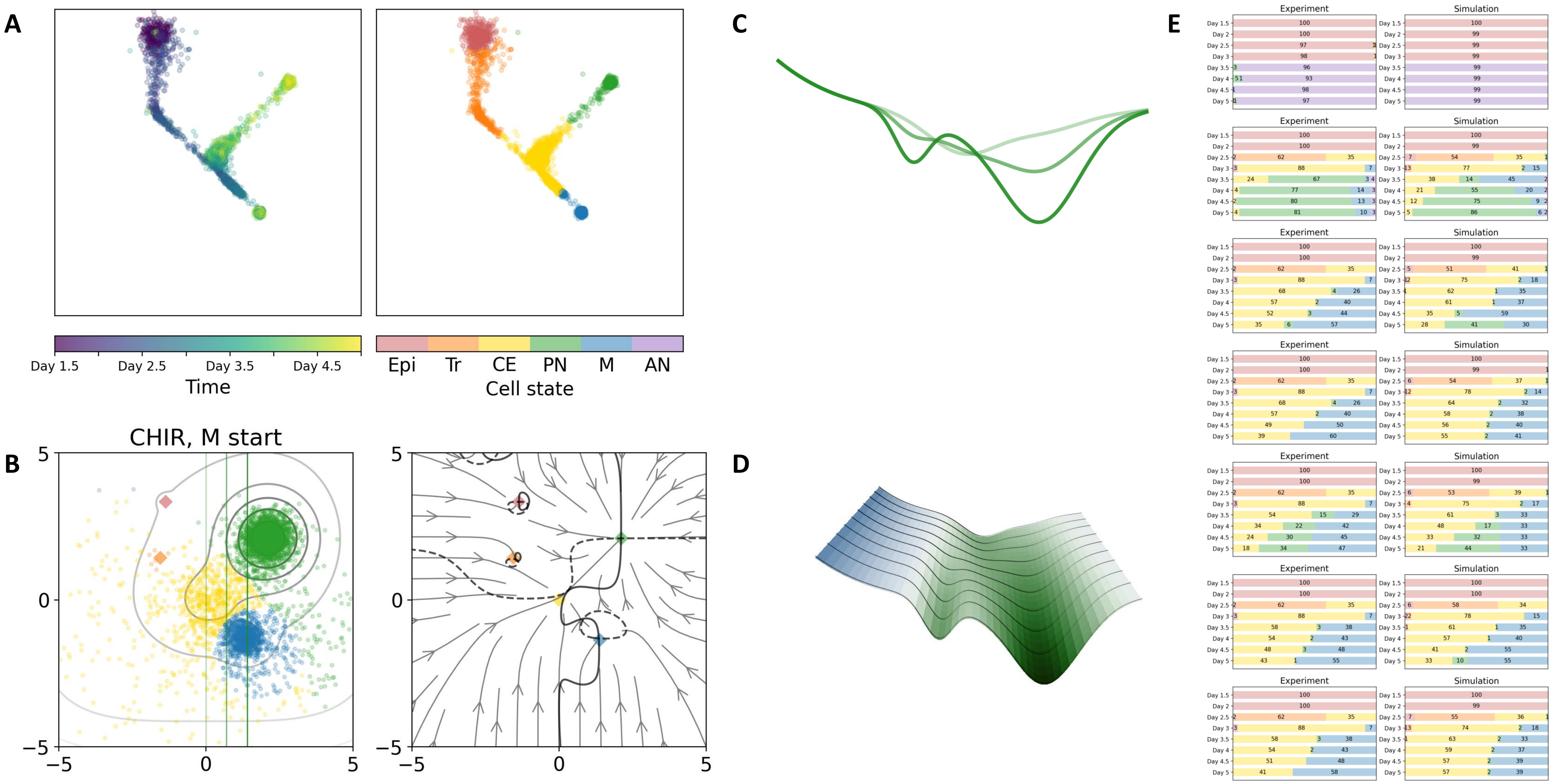

17 — Timecode: 20250531-115250

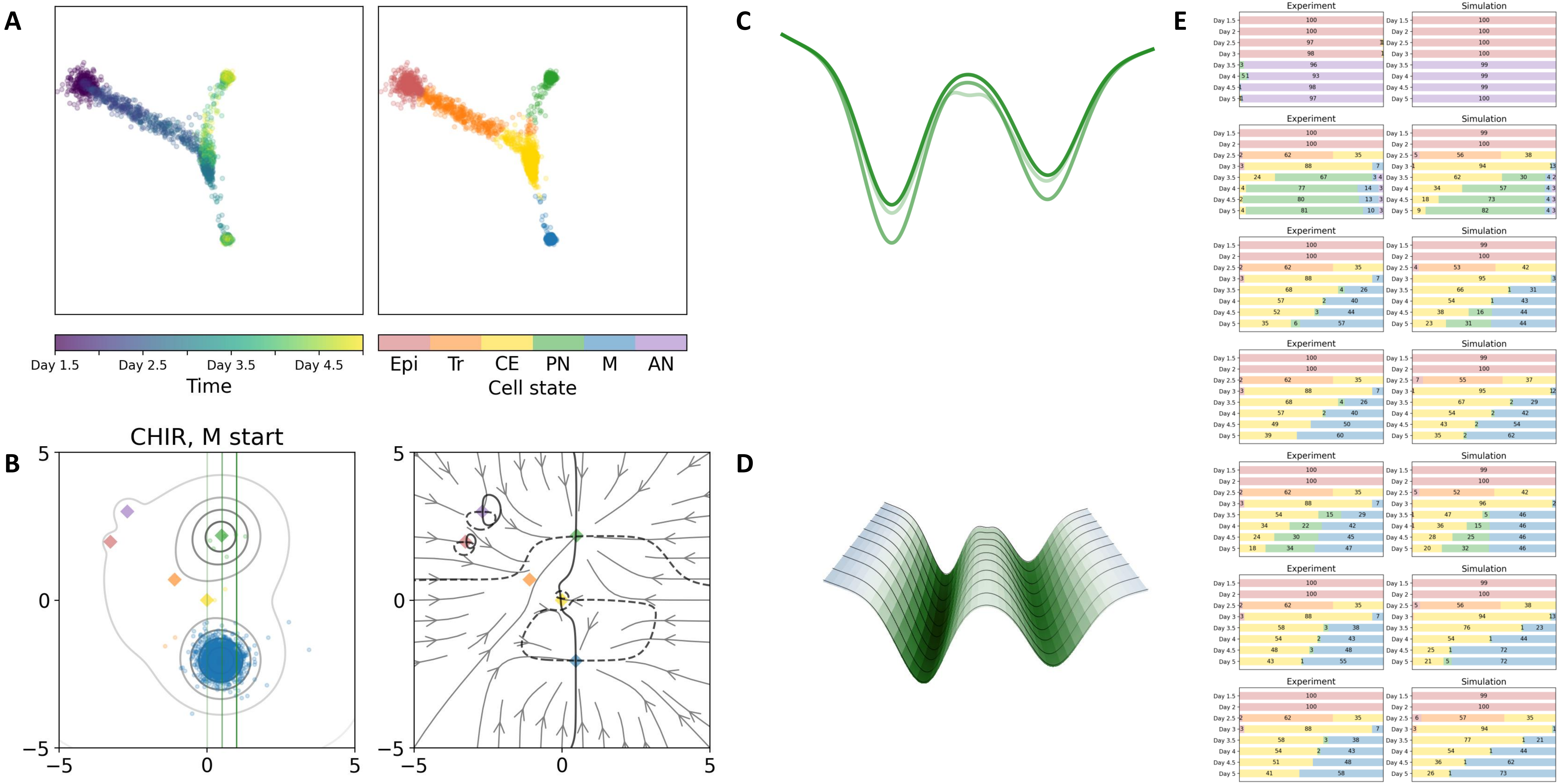

### 27 — Timecode: 20250601-041838

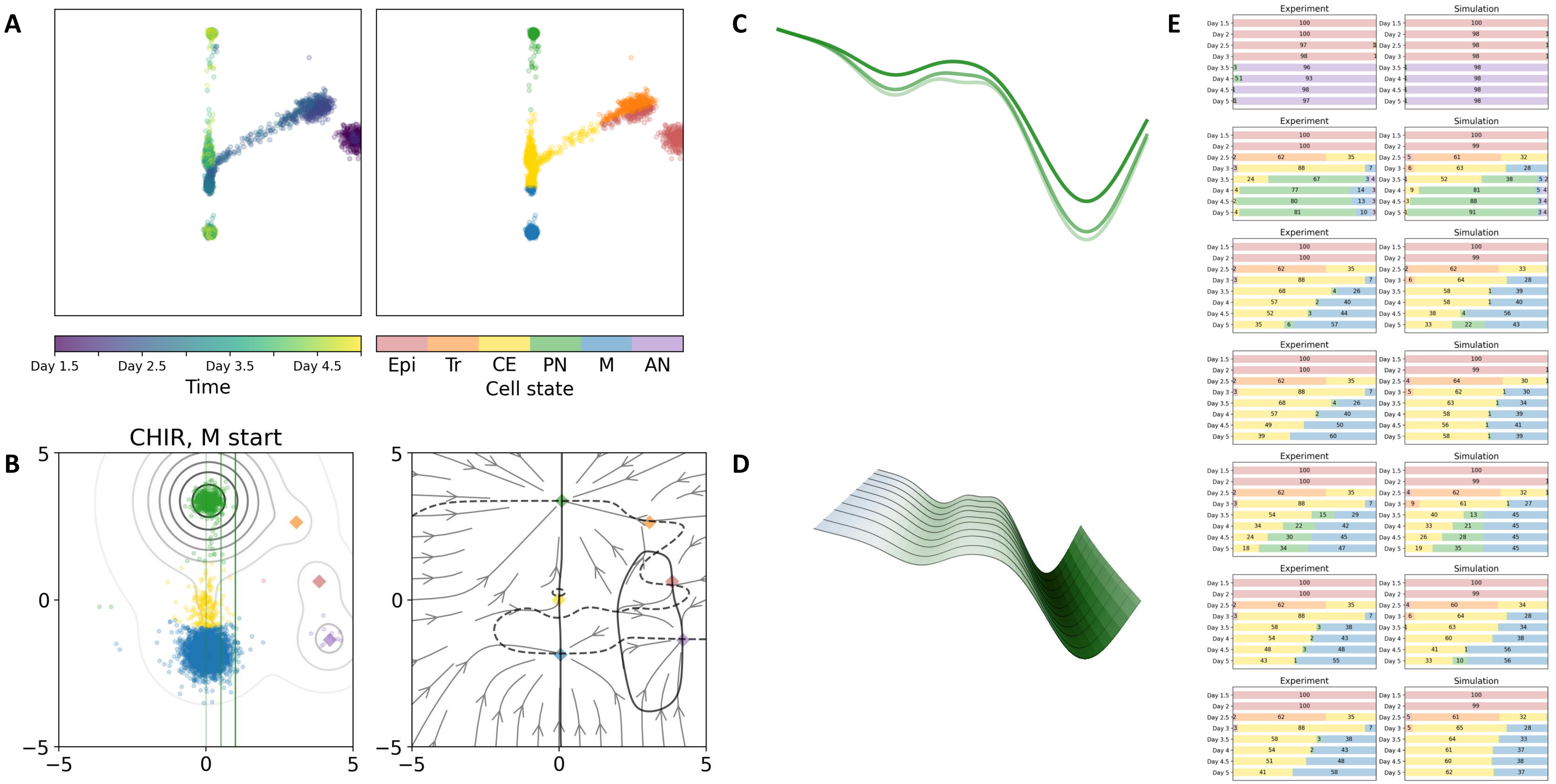

### 28 — Timecode: 20250601-043603

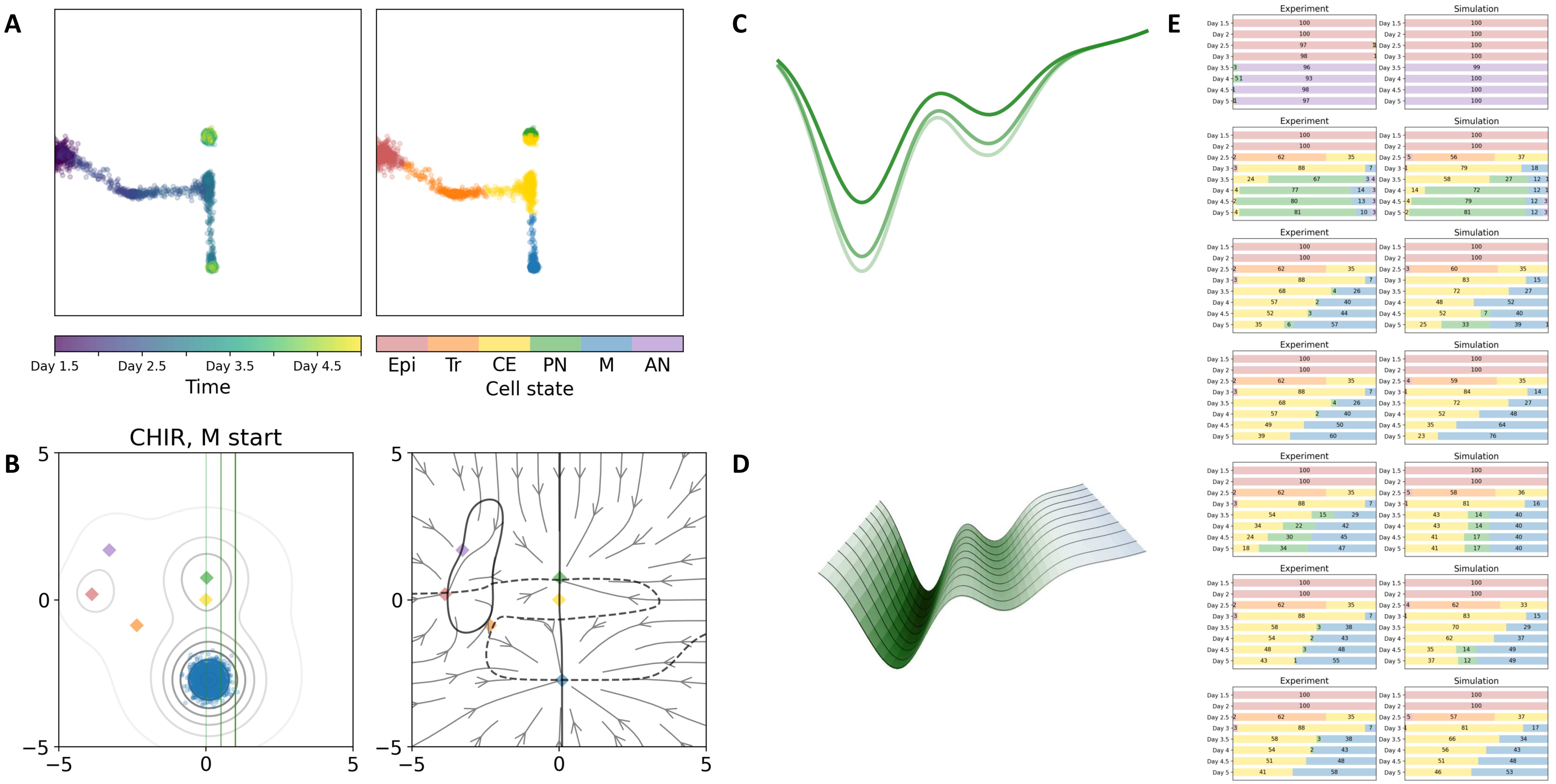

### 29 — Timecode: 20250601-051221

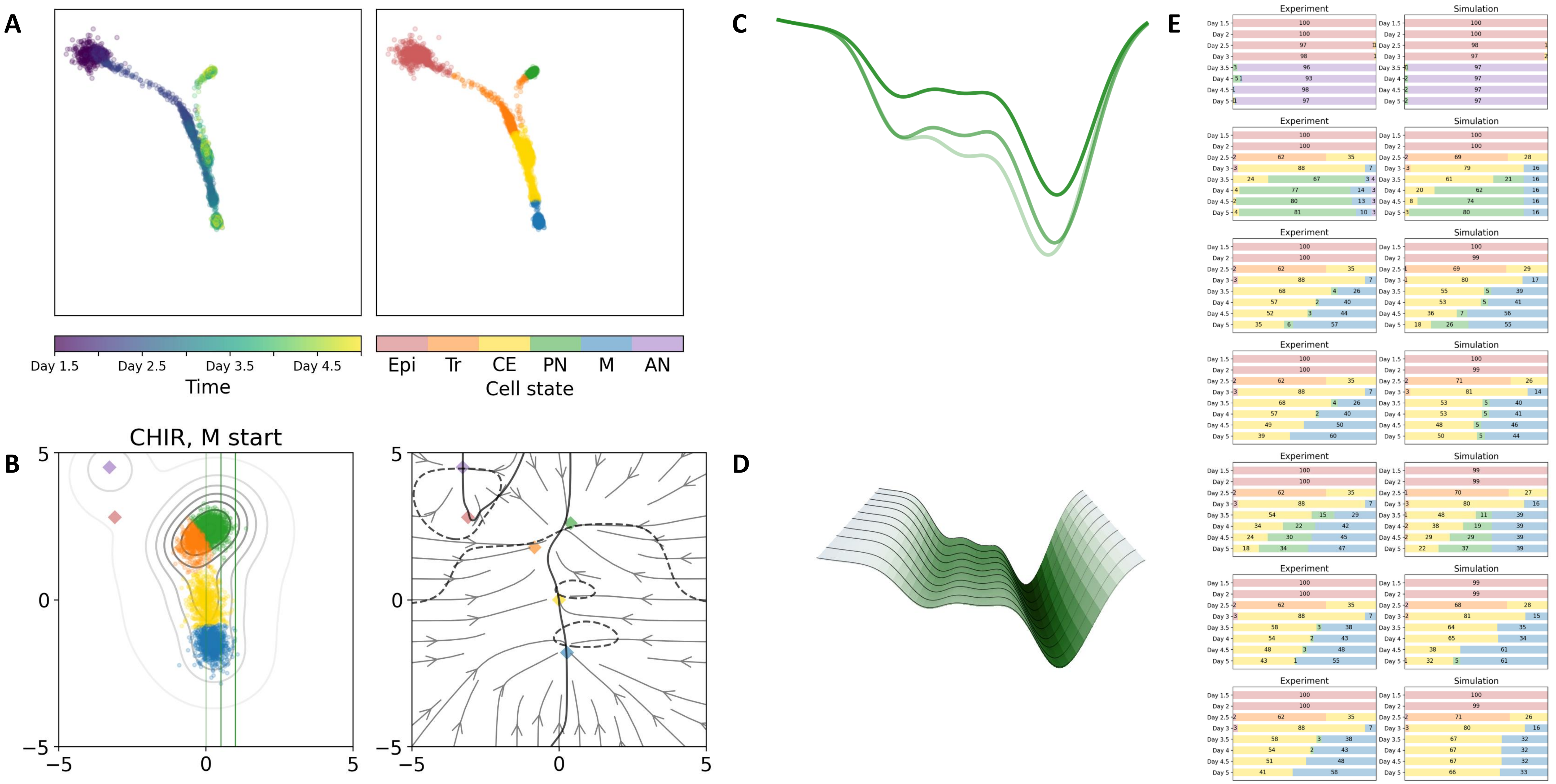

### 30 — Timecode: 20250601-053035

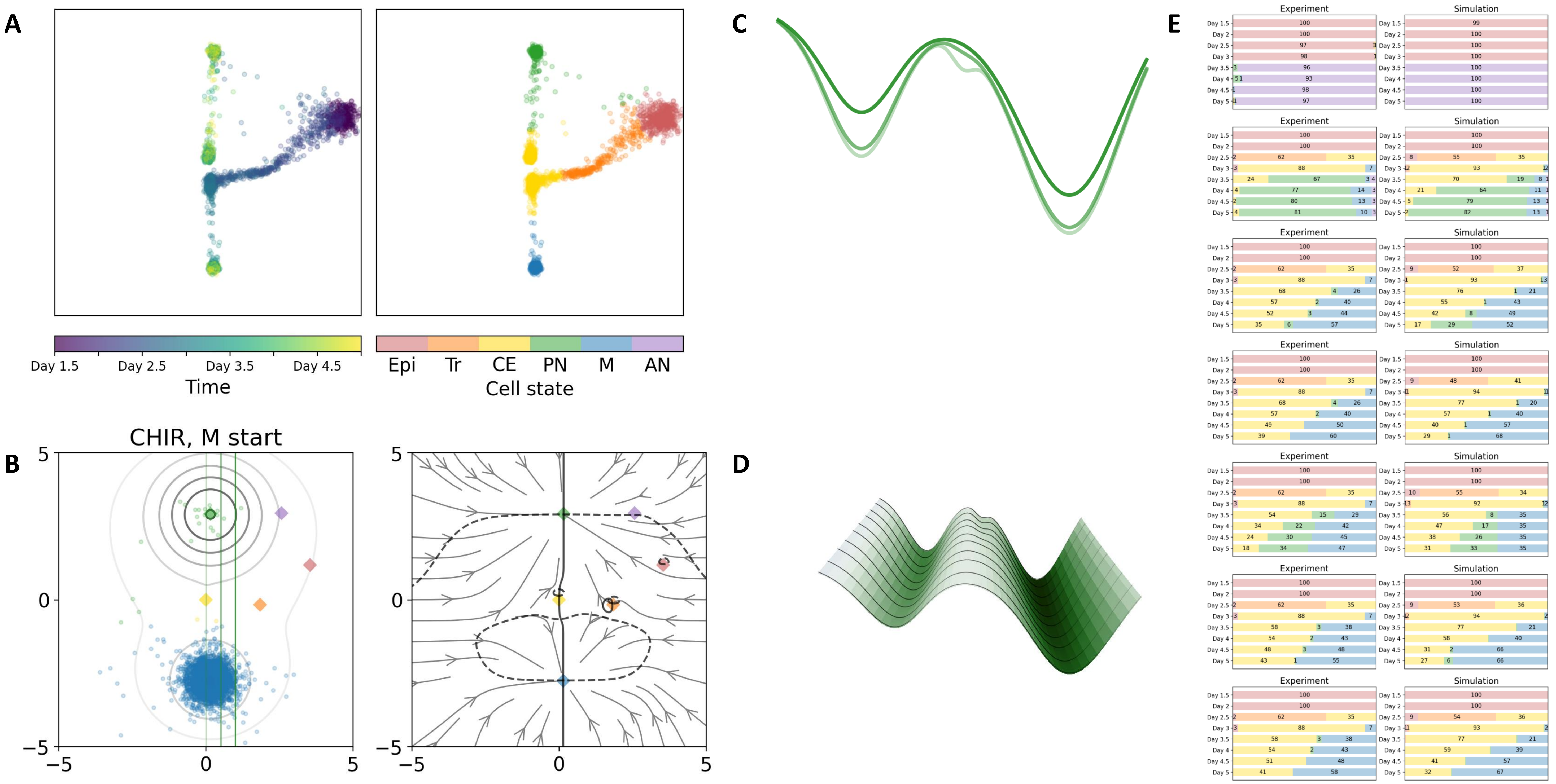

### 35 — Timecode: 20250601-114857

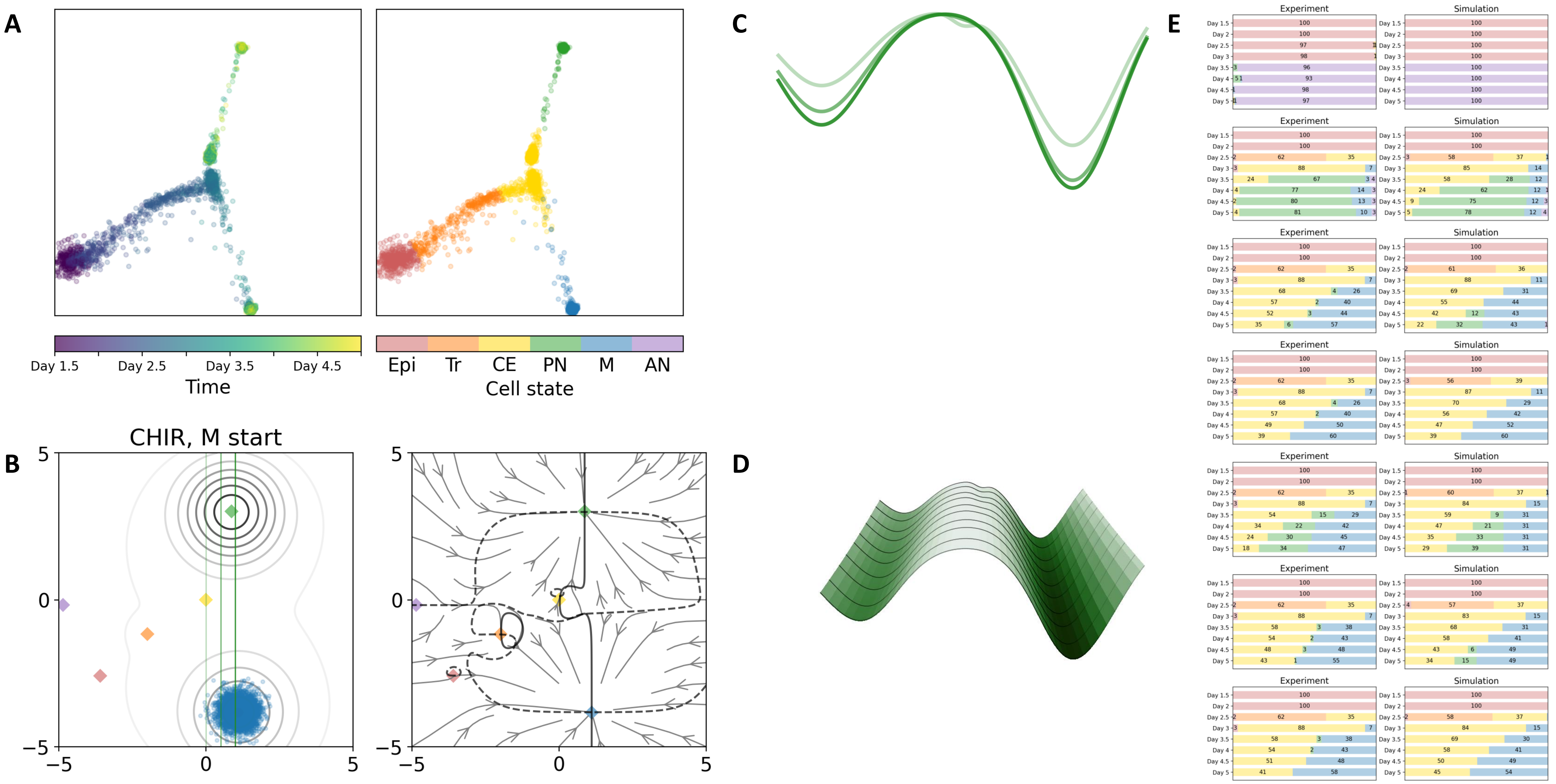

### 37 — Timecode: 20250601-145502

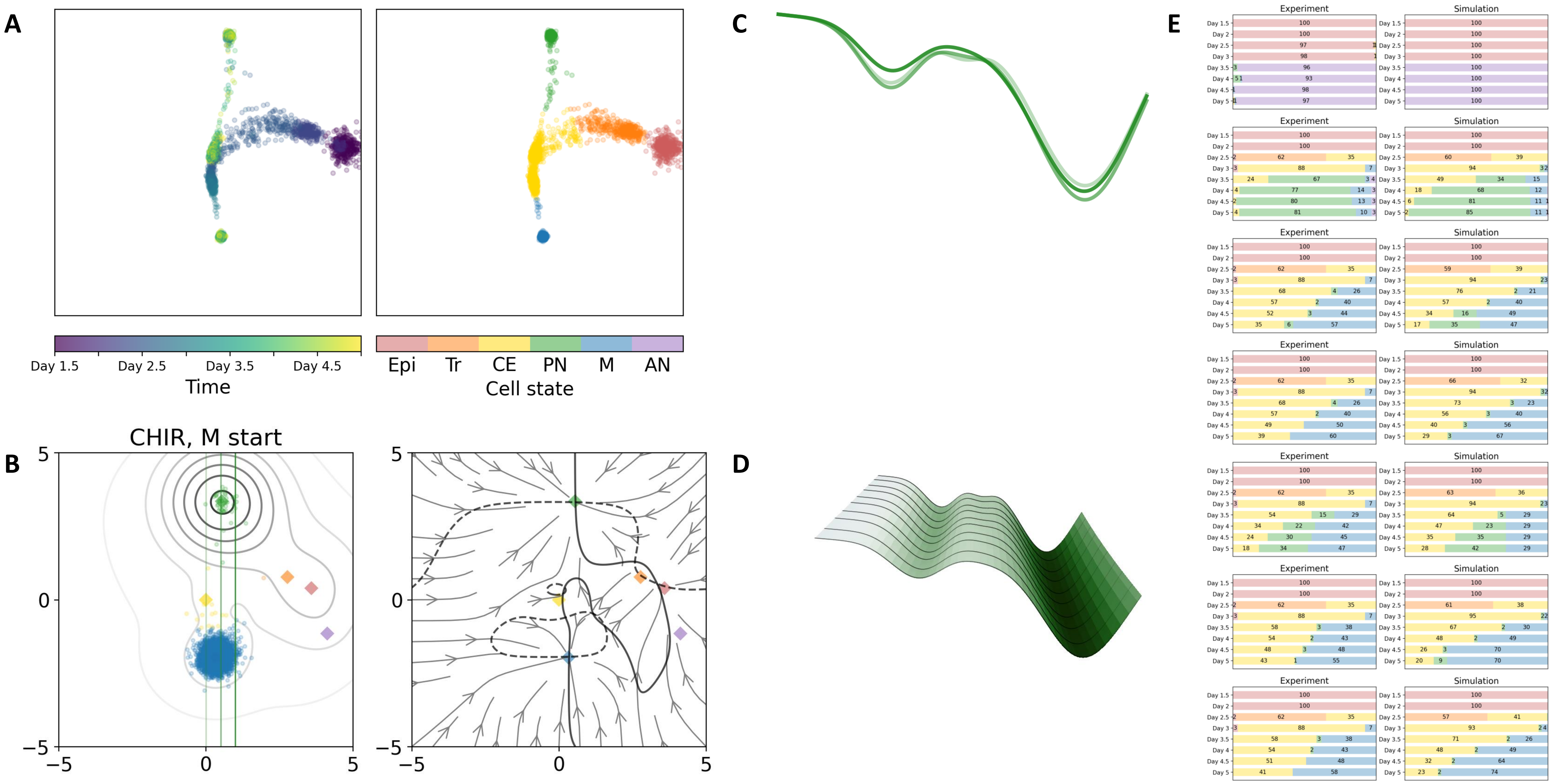

49 — Timecode: 20250602-063942

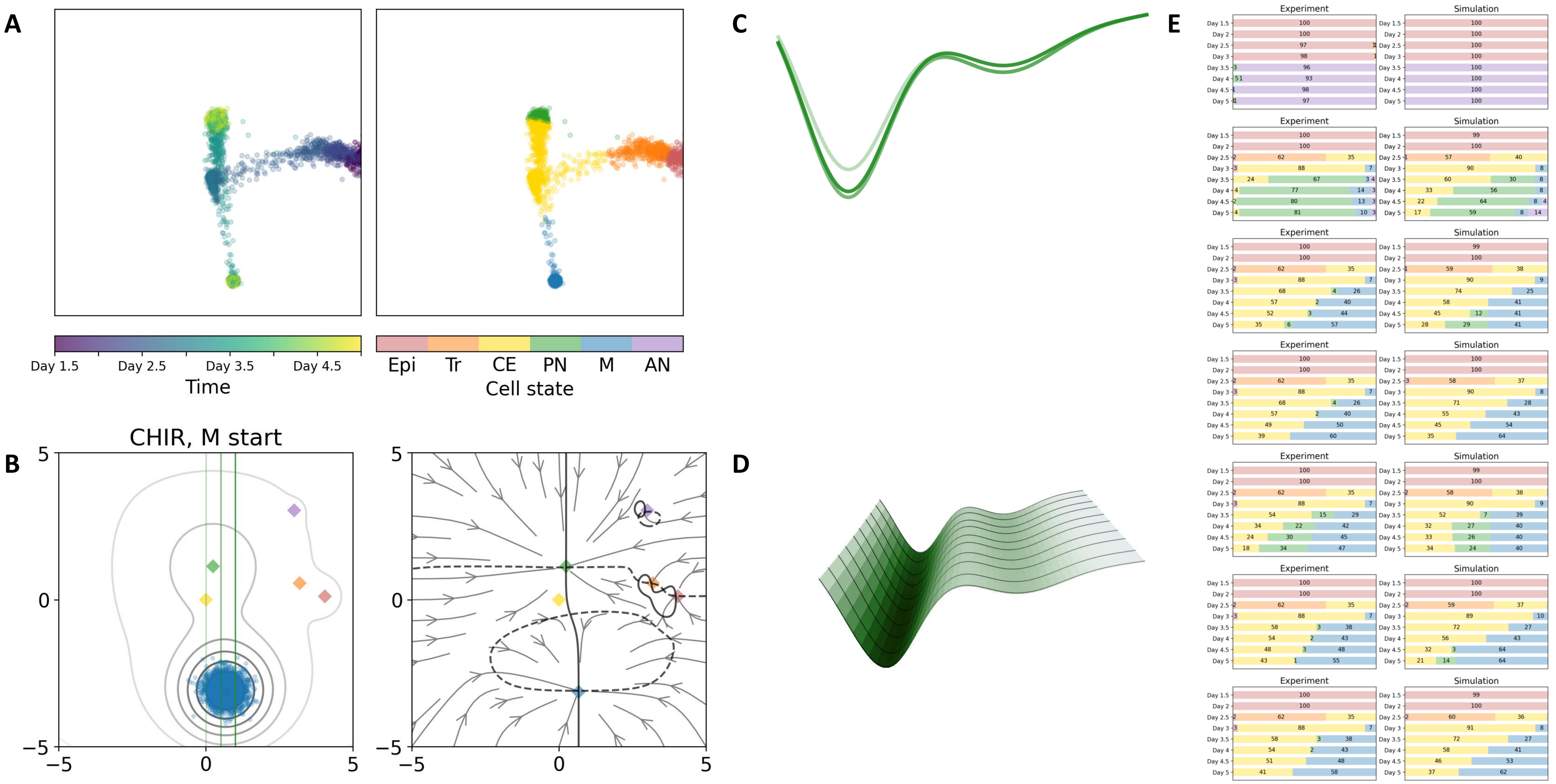

### 55 — Timecode: 20250602-125948

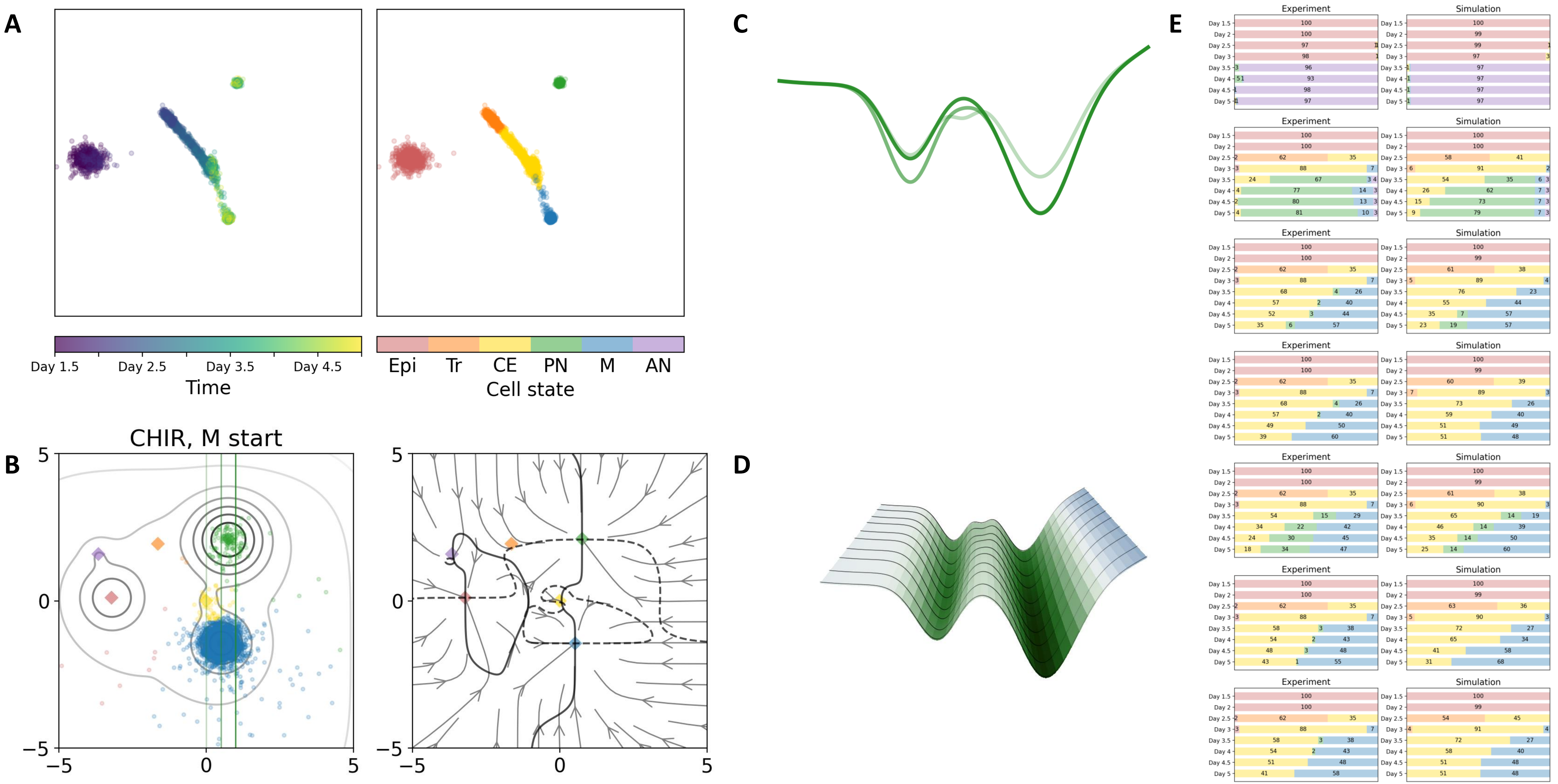

57 — Timecode: 20250602-152625

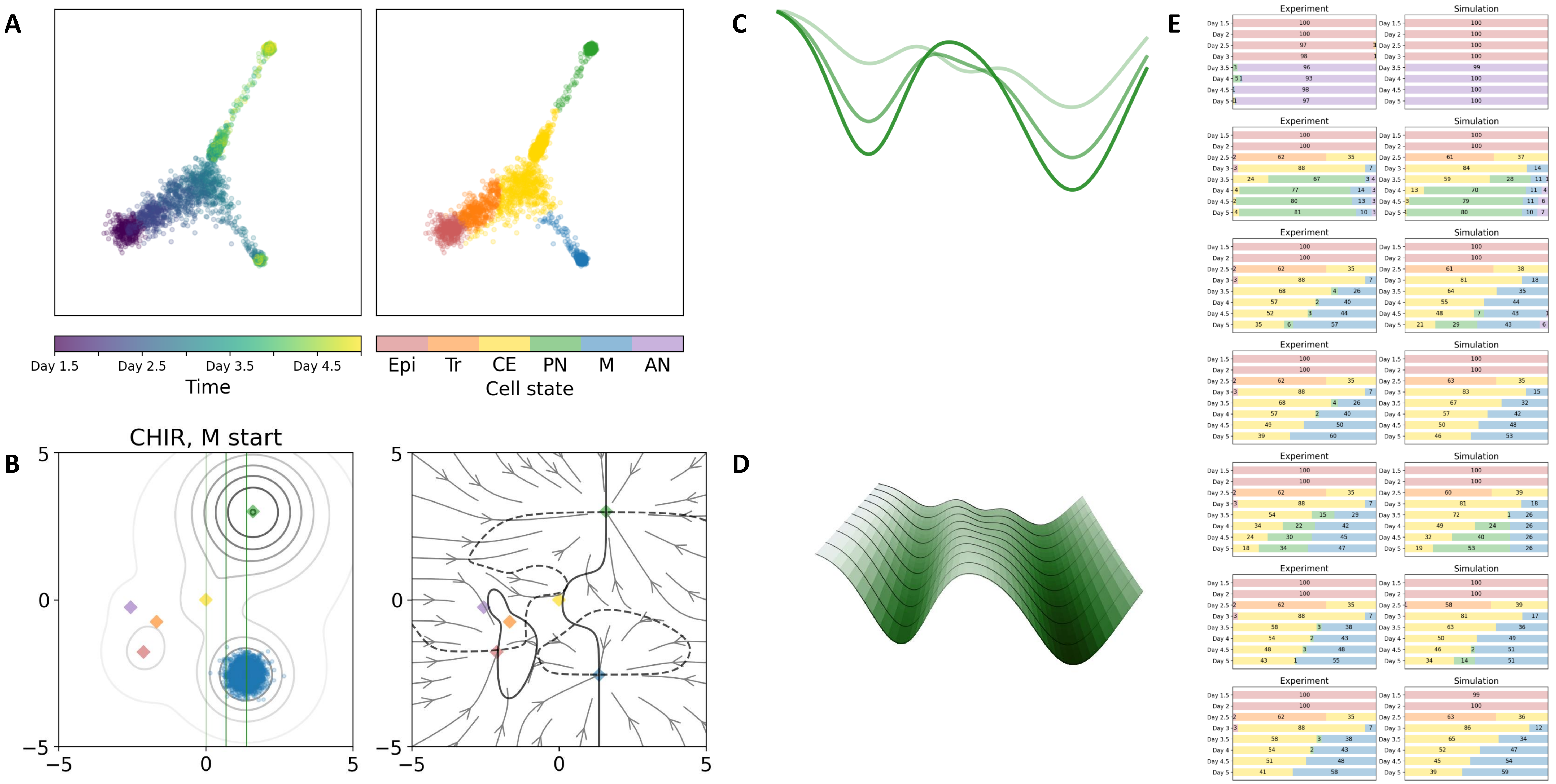

### Flat double cusp

### 03 — Timecode: 20250530-210216

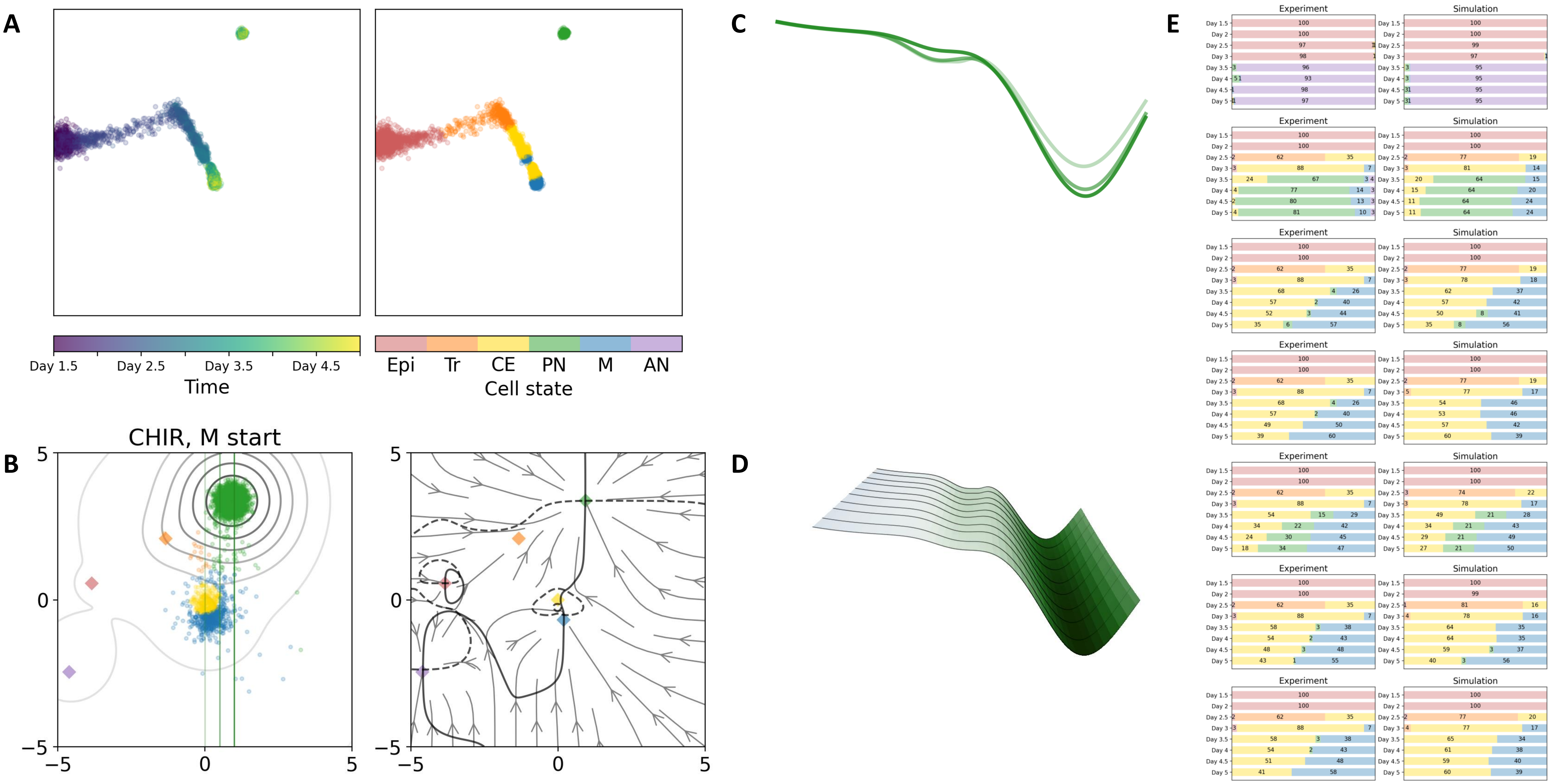

### 04 — Timecode: 20250530-234414

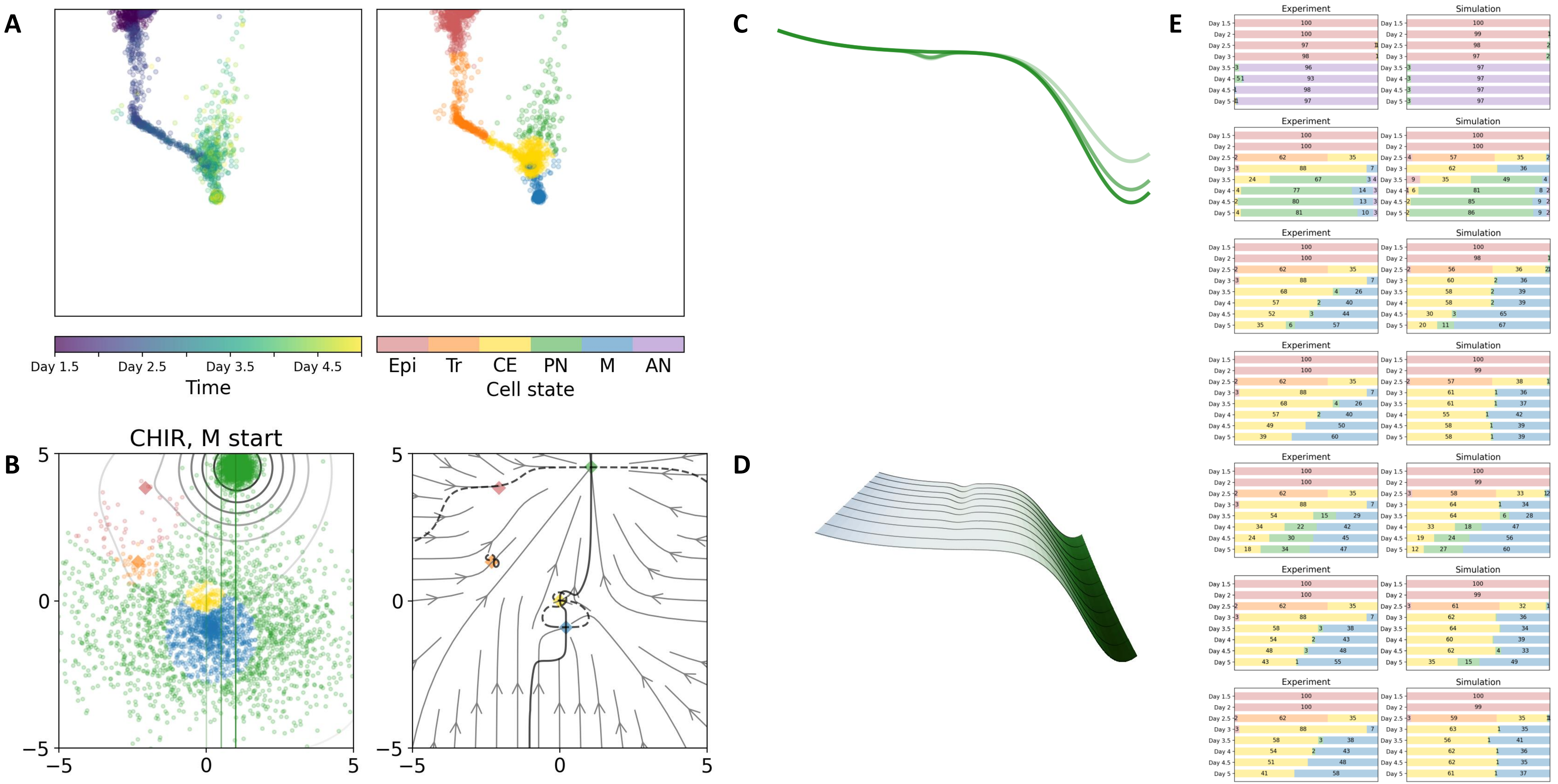

### 05 — Timecode: 20250531-002107

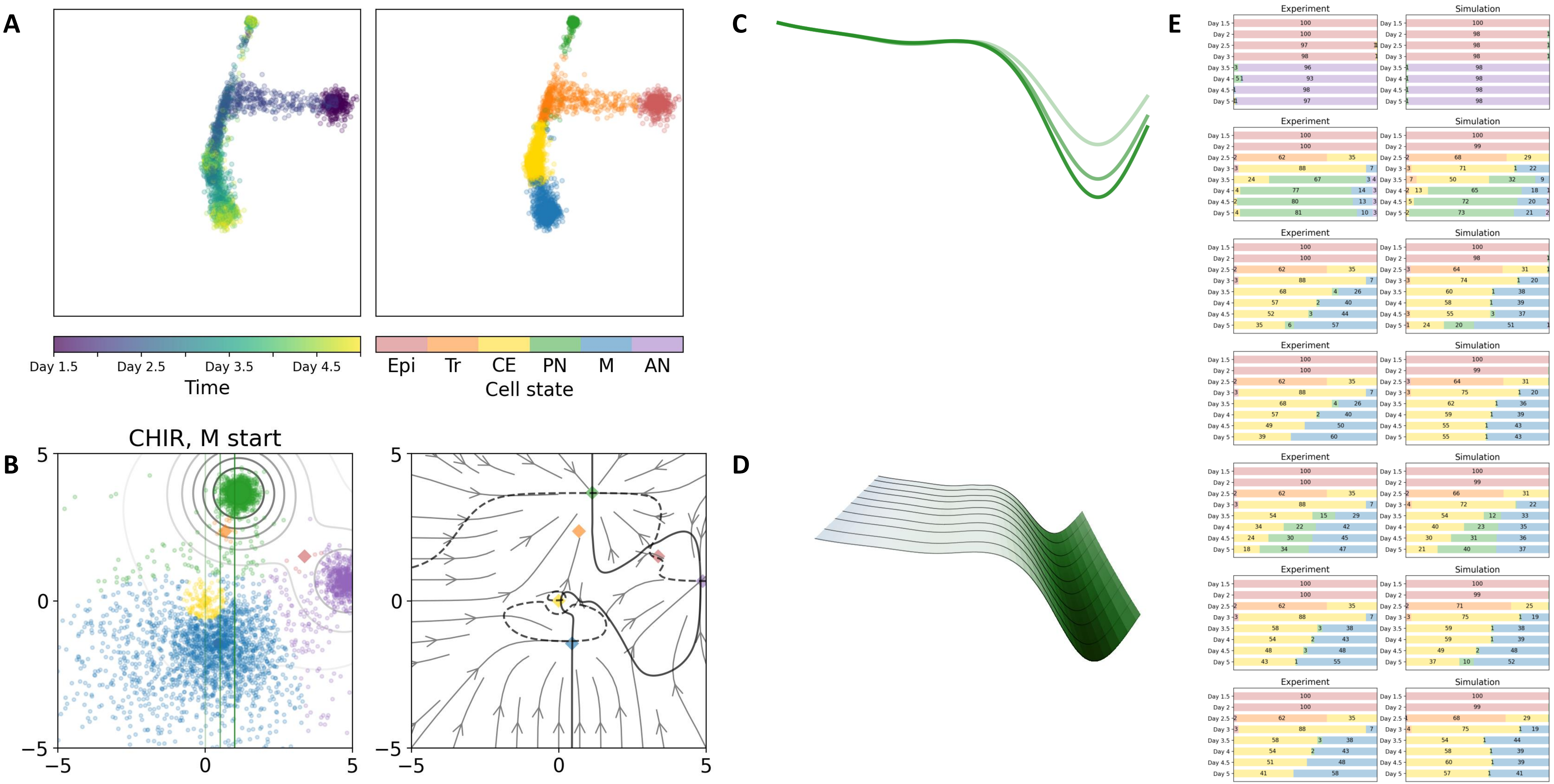

### 13 — Timecode: 20250531-064222

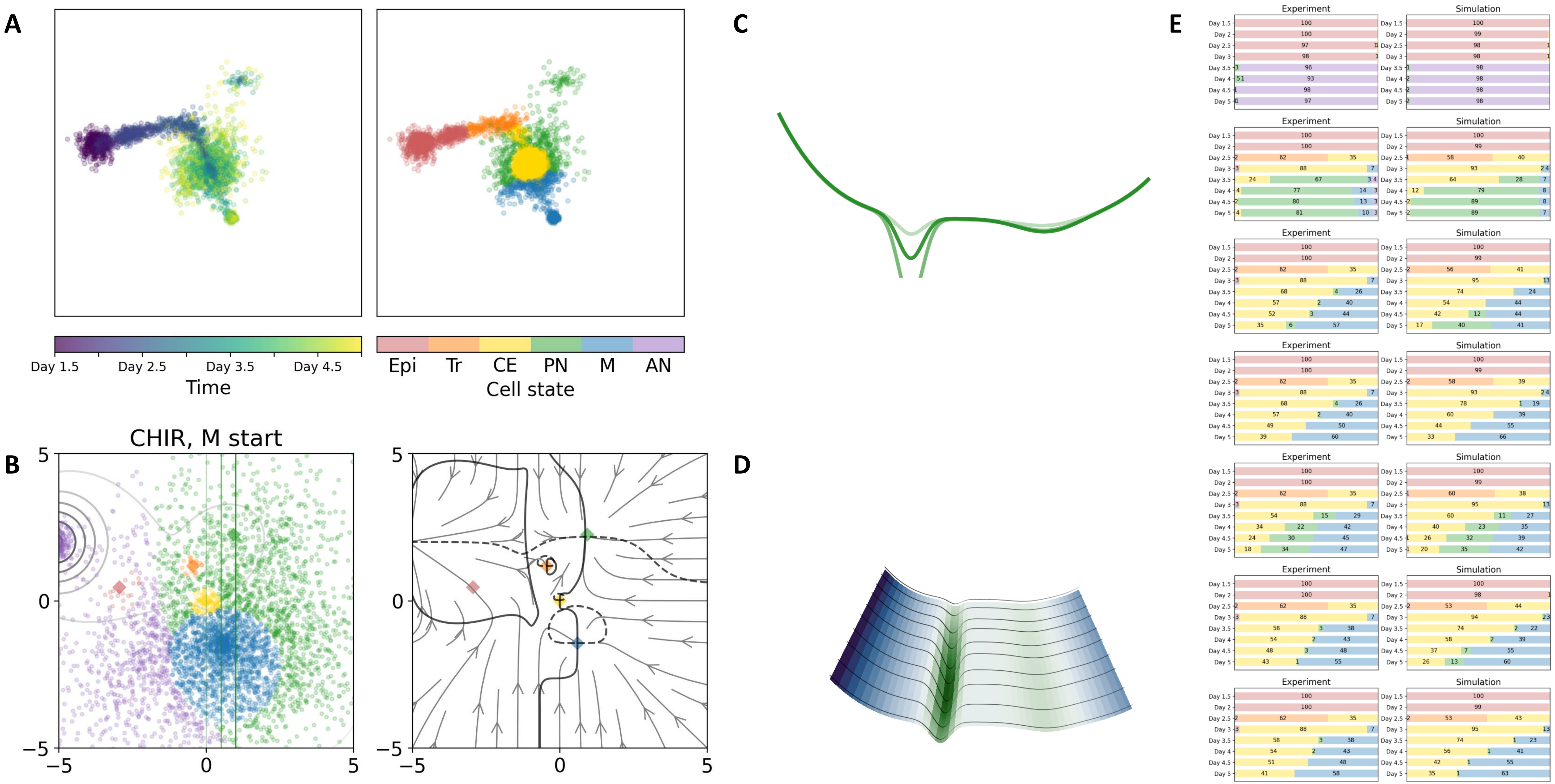

### 16 — Timecode: 20250531-105654

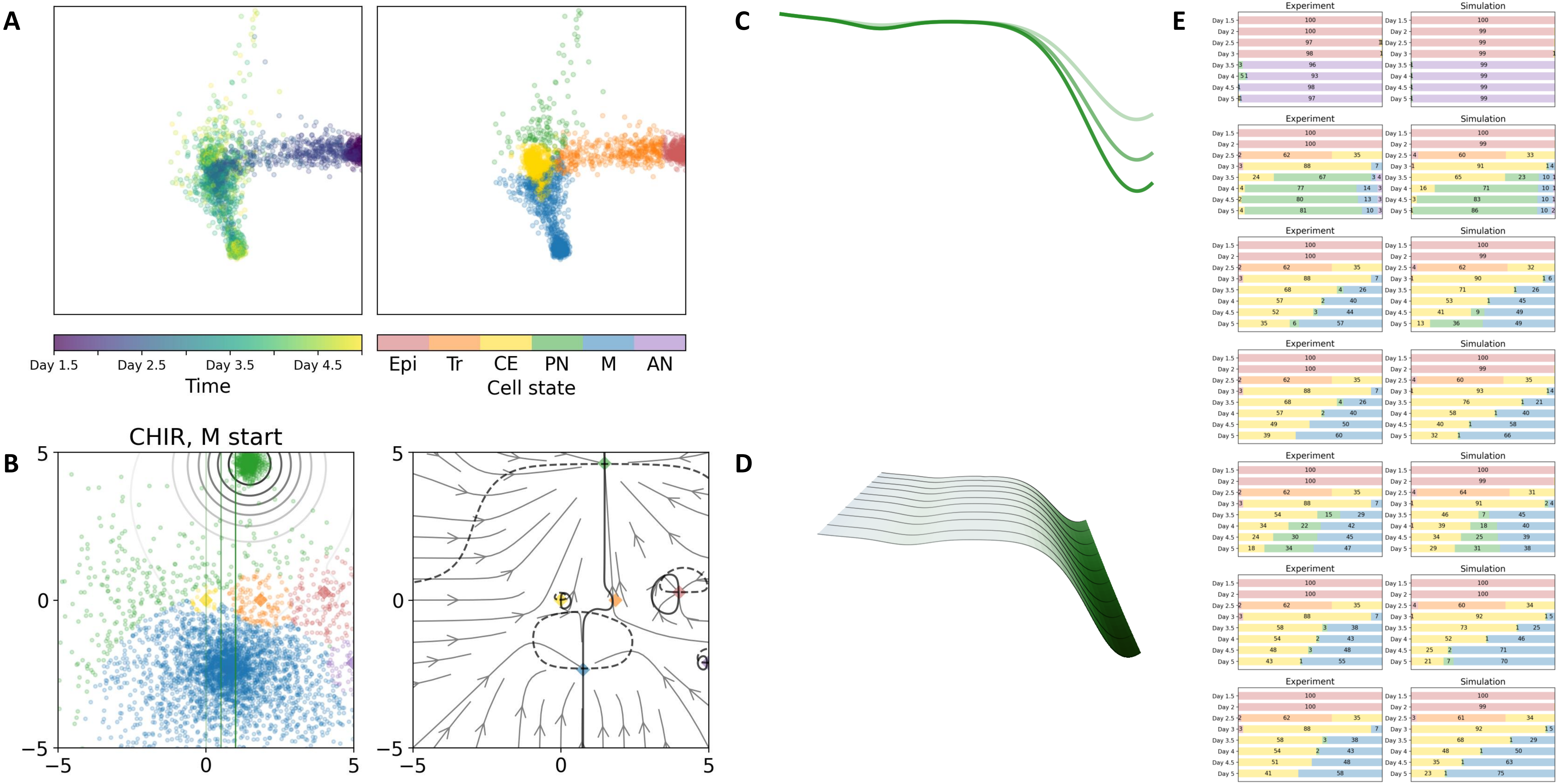

### 21 — Timecode: 20250531-174321

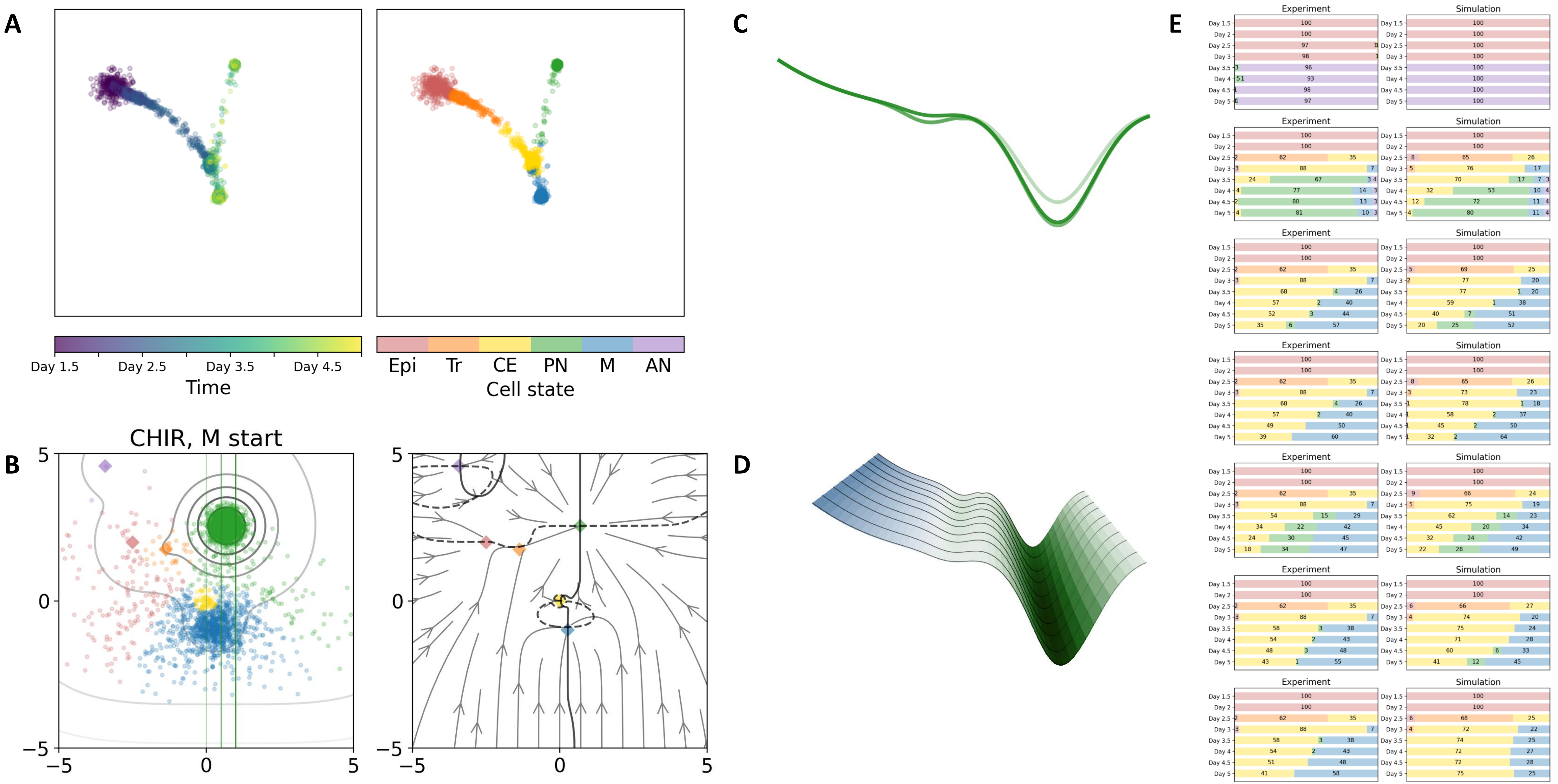

### 31 — Timecode: 20250601-070205

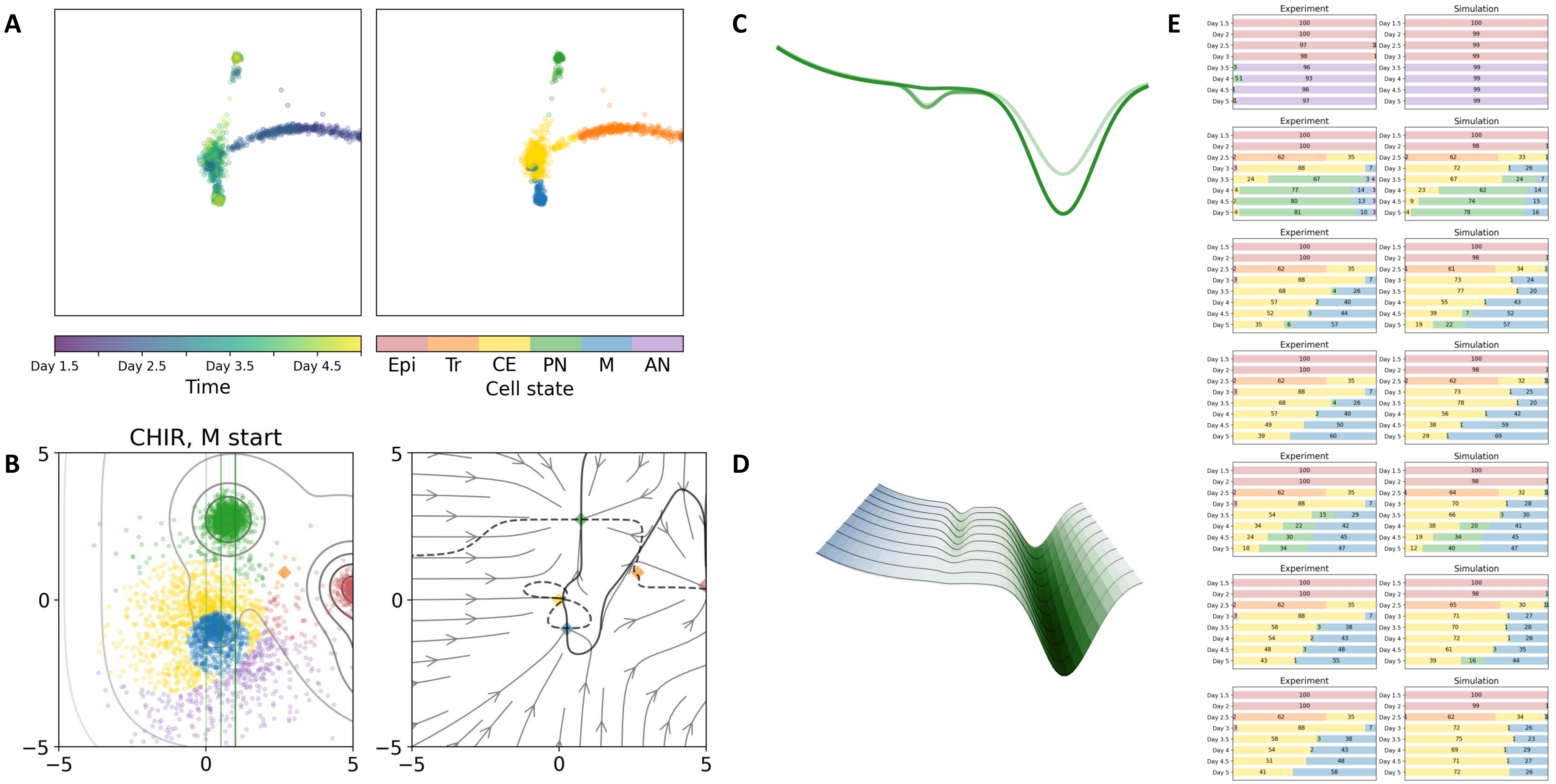

### 46 — Timecode: 20250602-033533

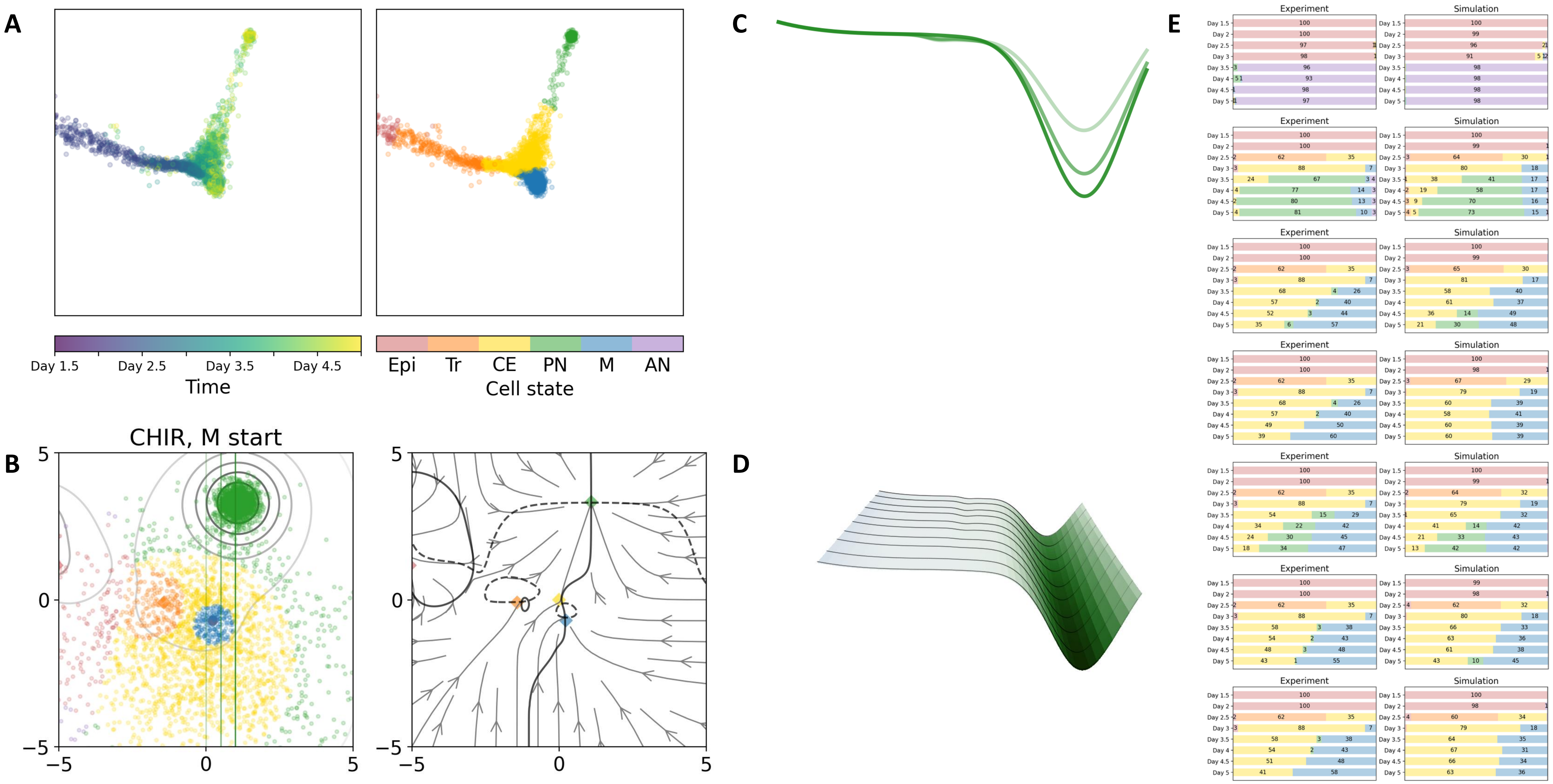

### 47 — Timecode: 20250602-042754

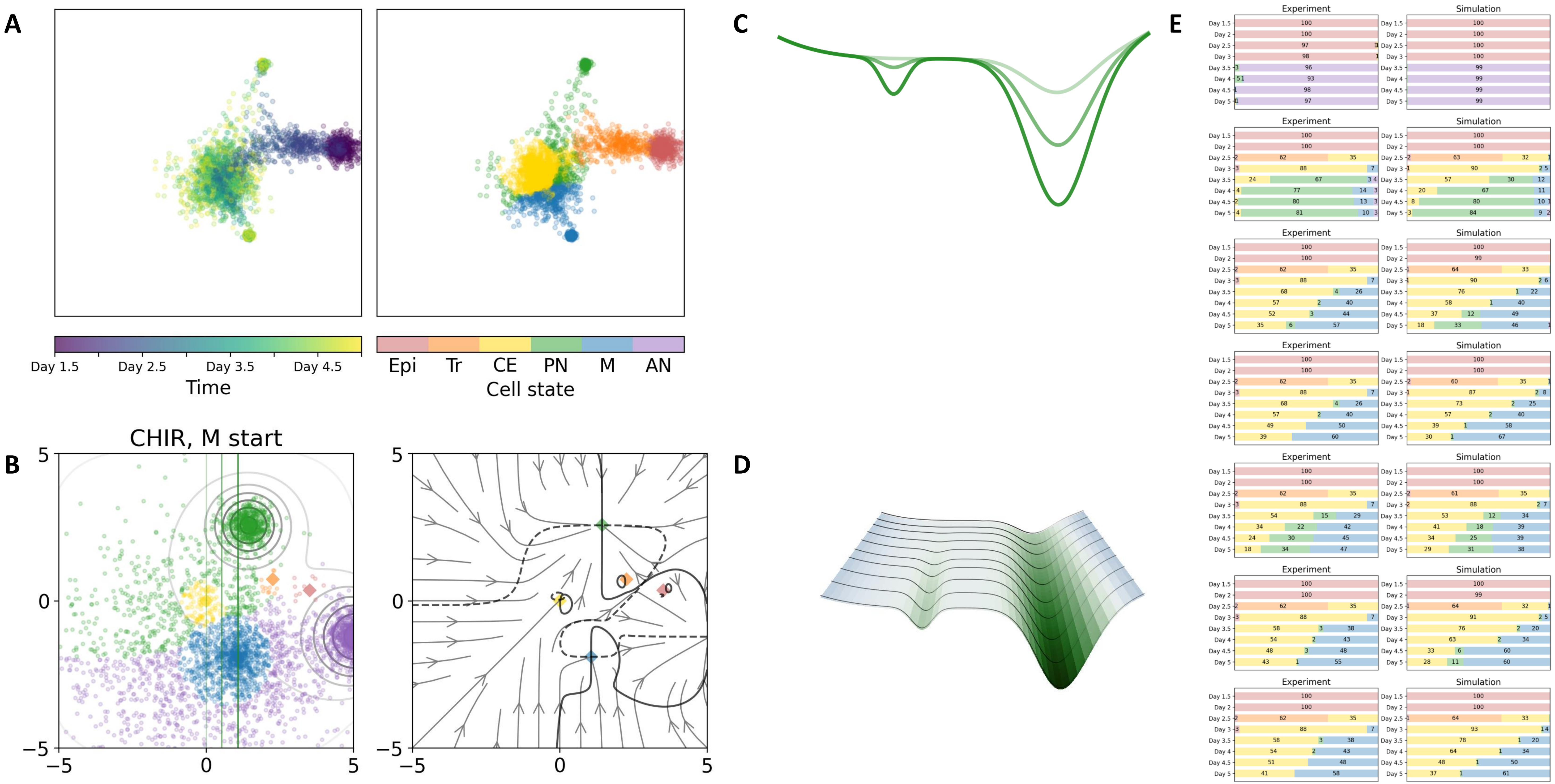

### 54 — Timecode: 20250602-100921

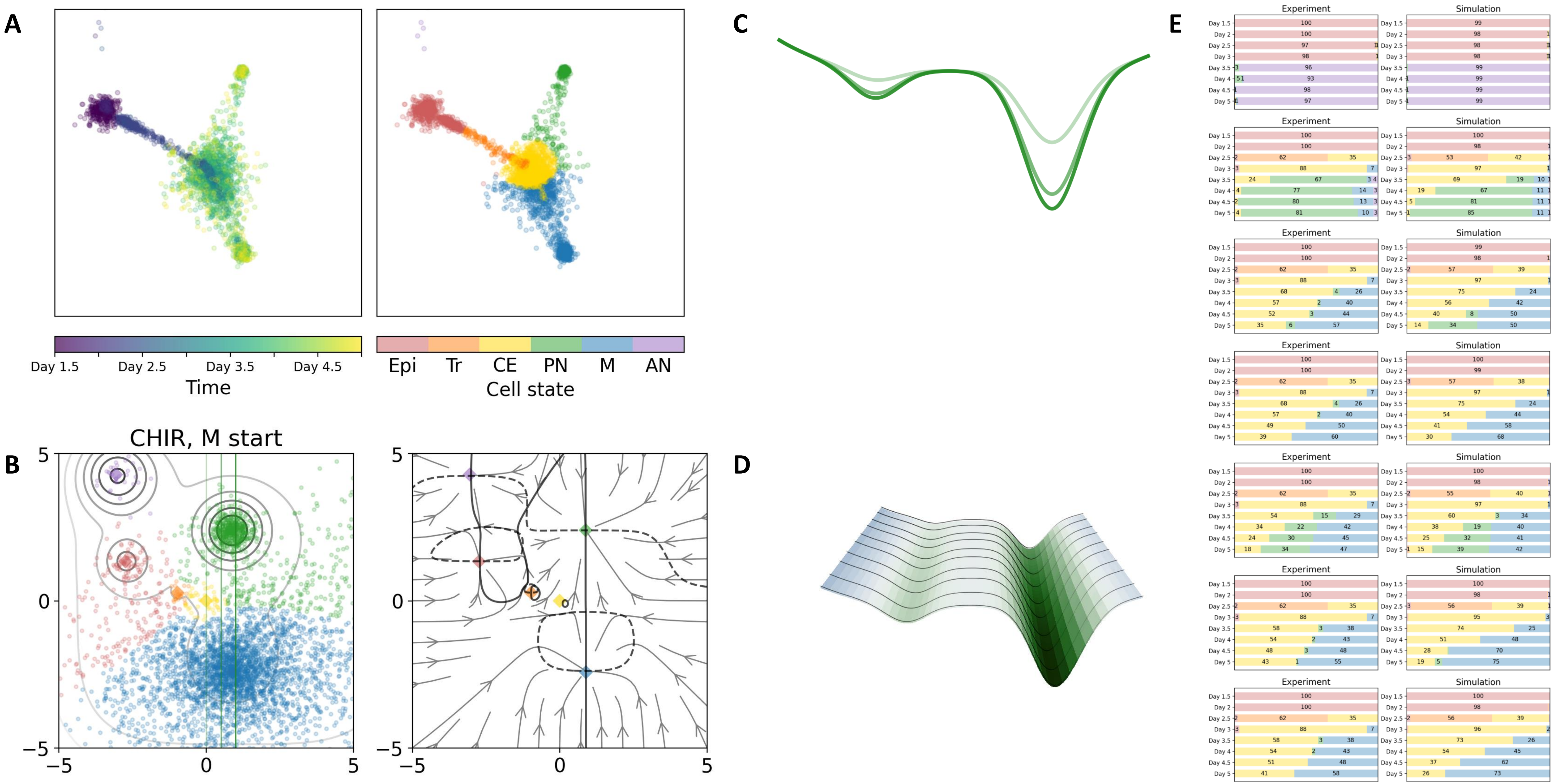

### 58 — Timecode: 20250602-160410

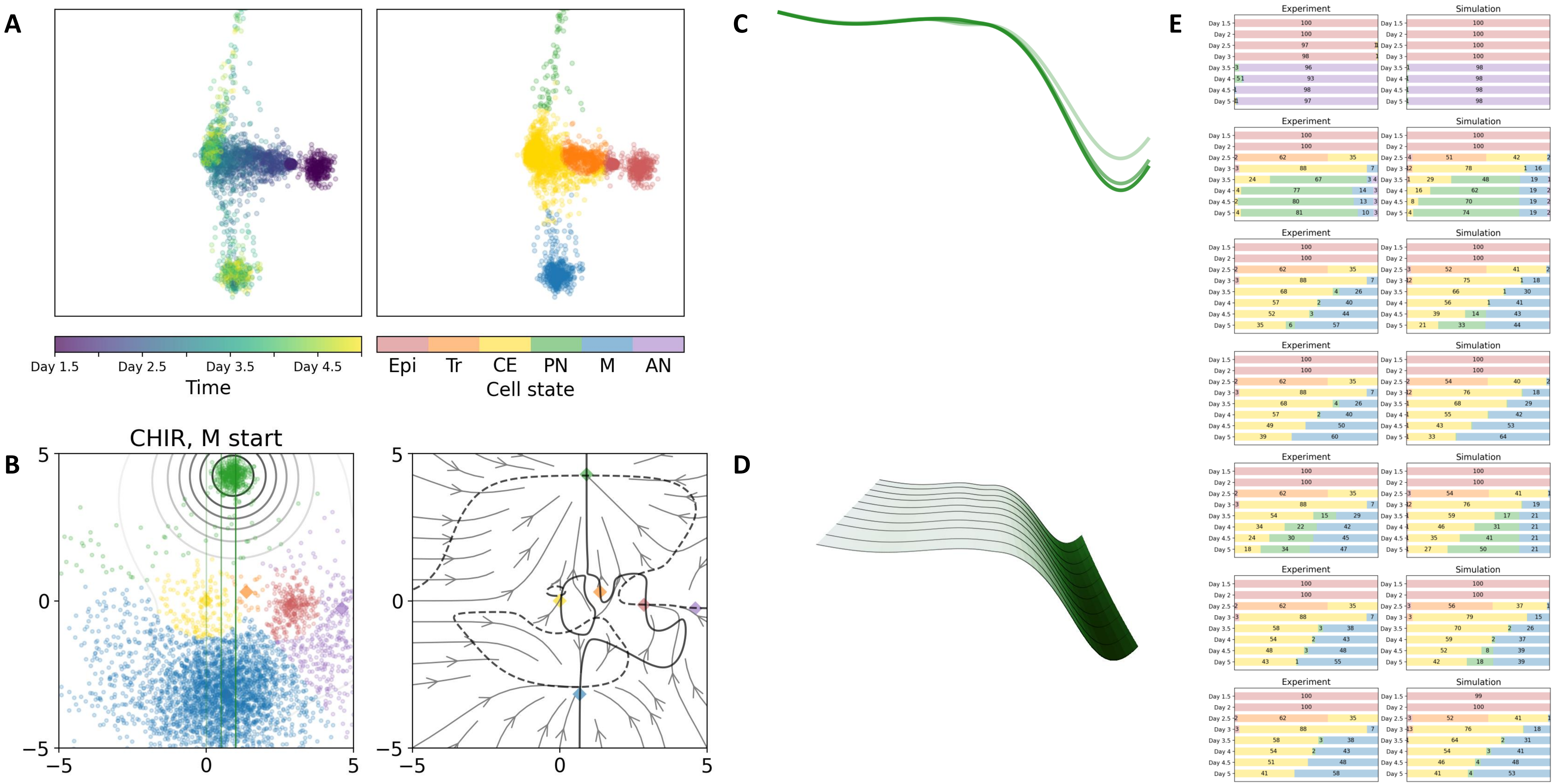

### Transient/mixed states

### 01 — Timecode: 20250530-172138

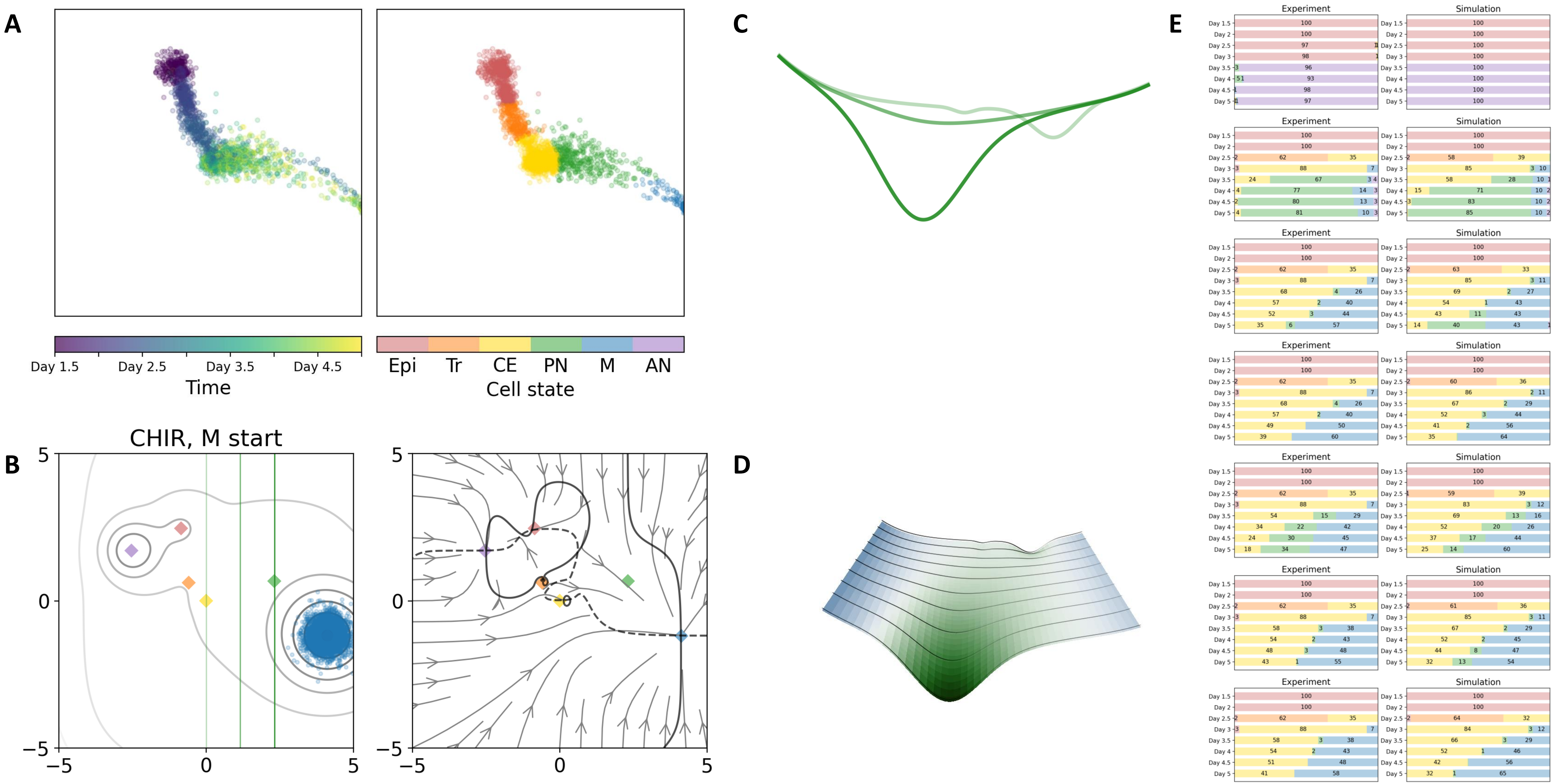

### 06 — Timecode: 20250531-004109

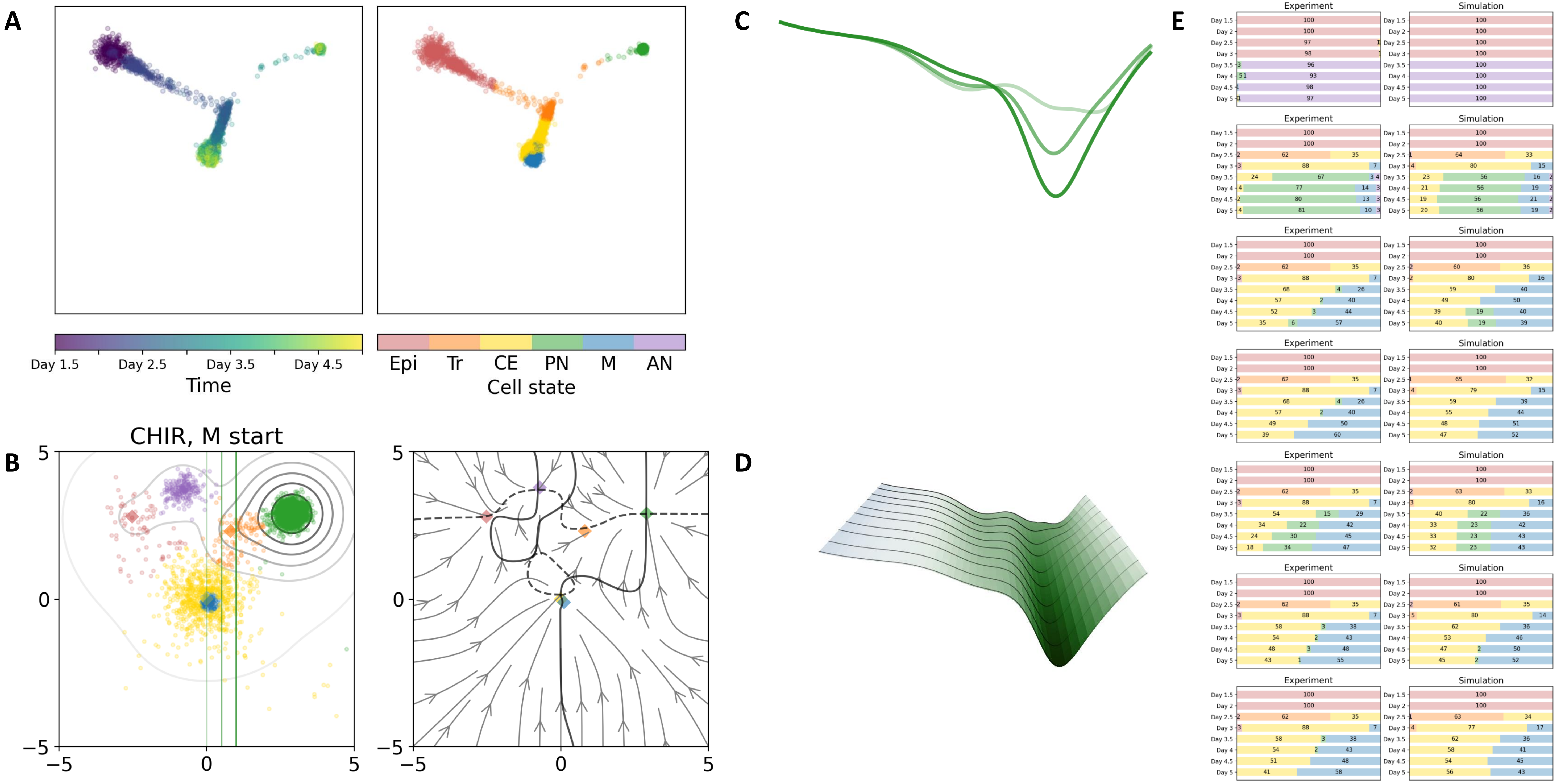

15 — Timecode: 20250531-081158

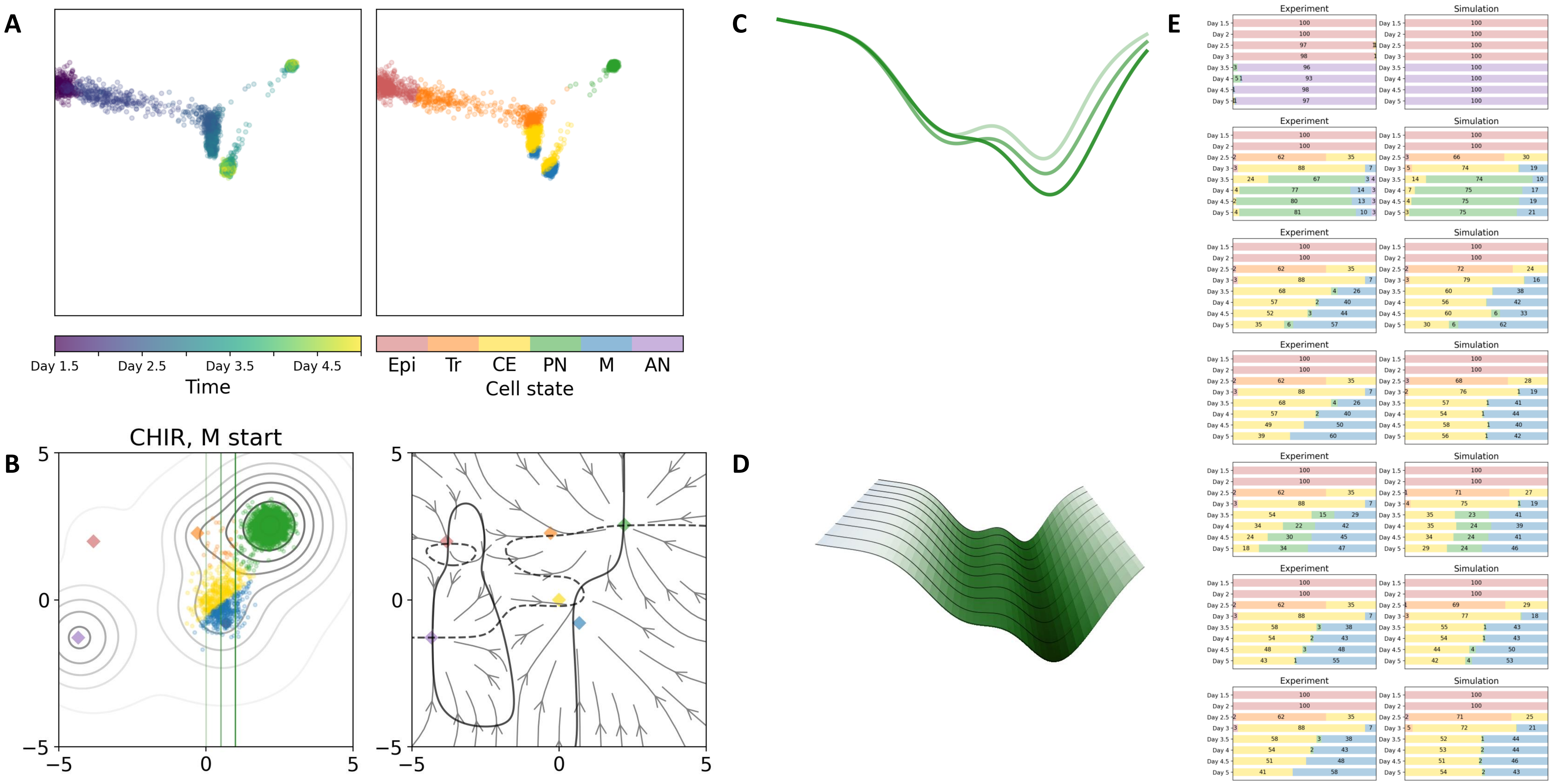

### 20 — Timecode: 20250531-145452

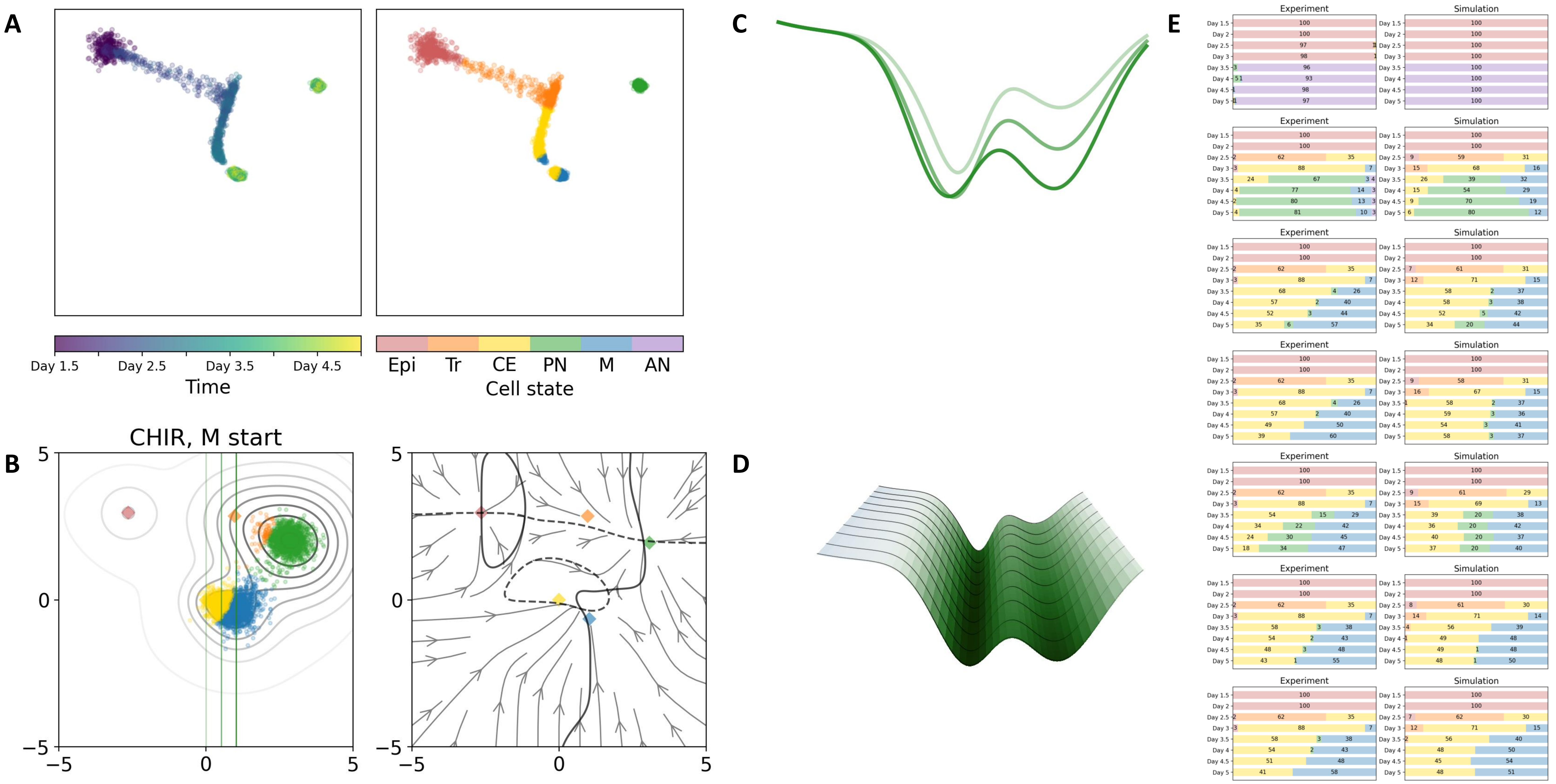

### 22 — Timecode: 20250531-222103

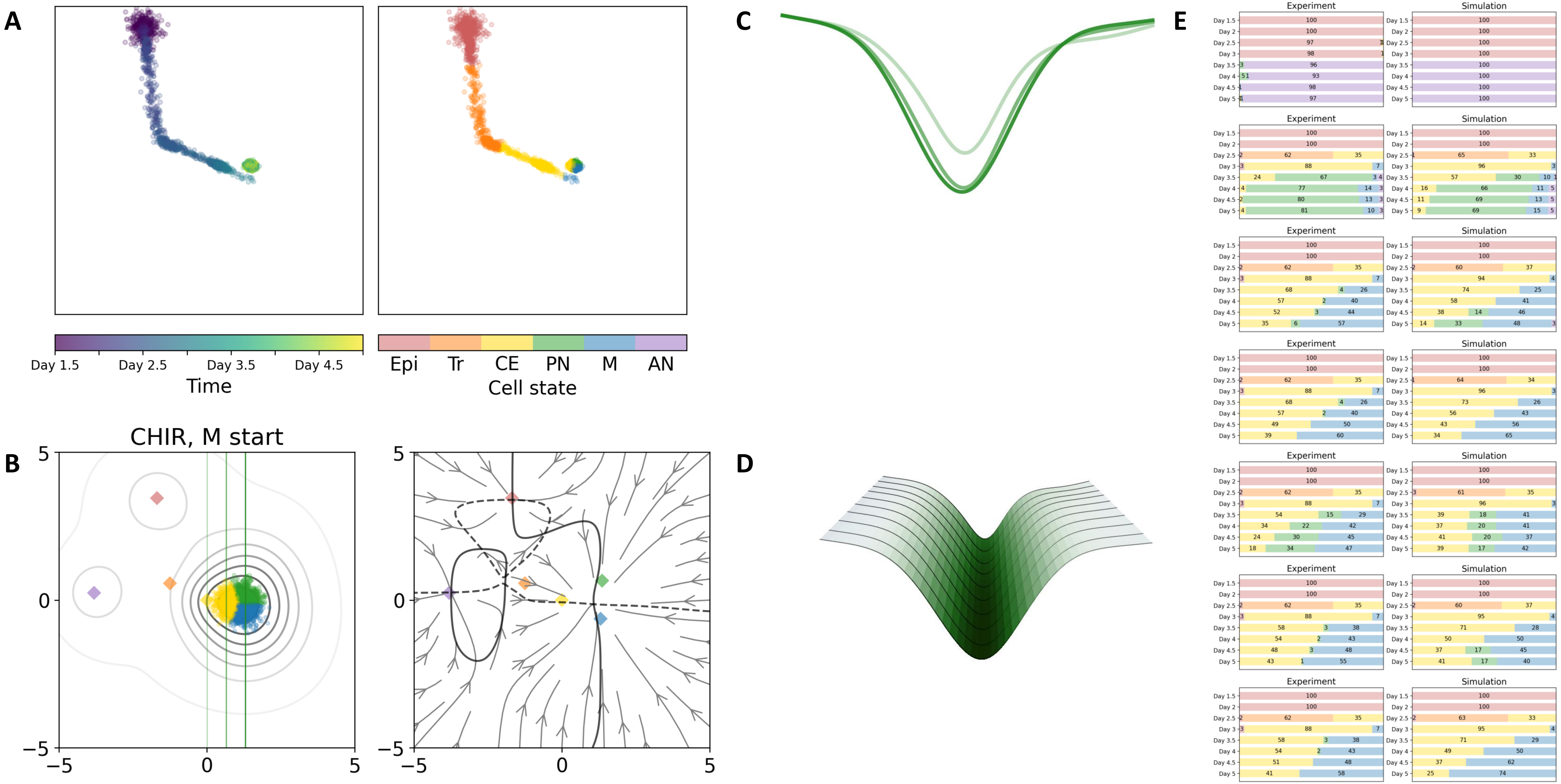

### 24 — Timecode: 20250601-002613

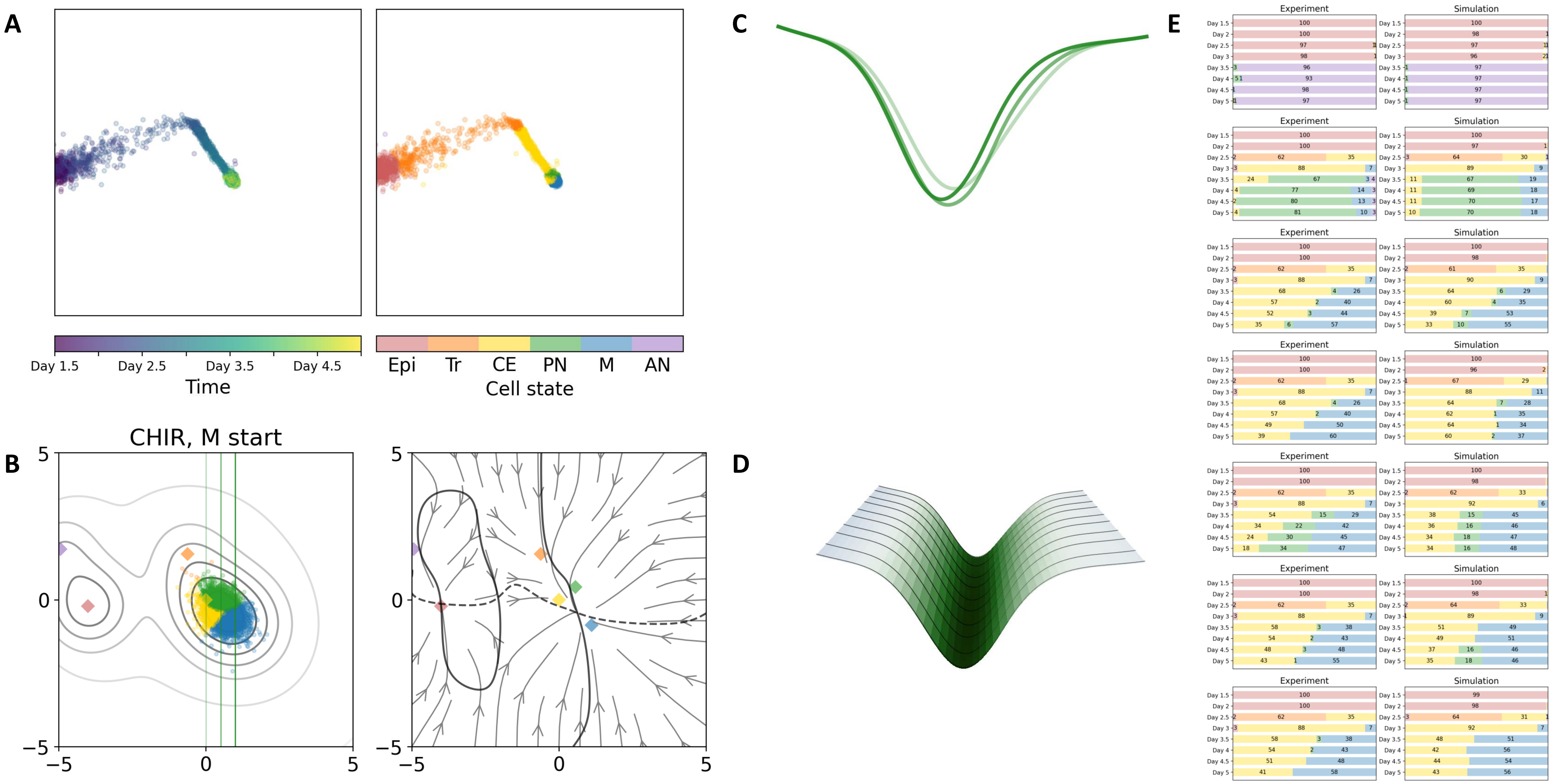

### 39 — Timecode: 20250601-155031

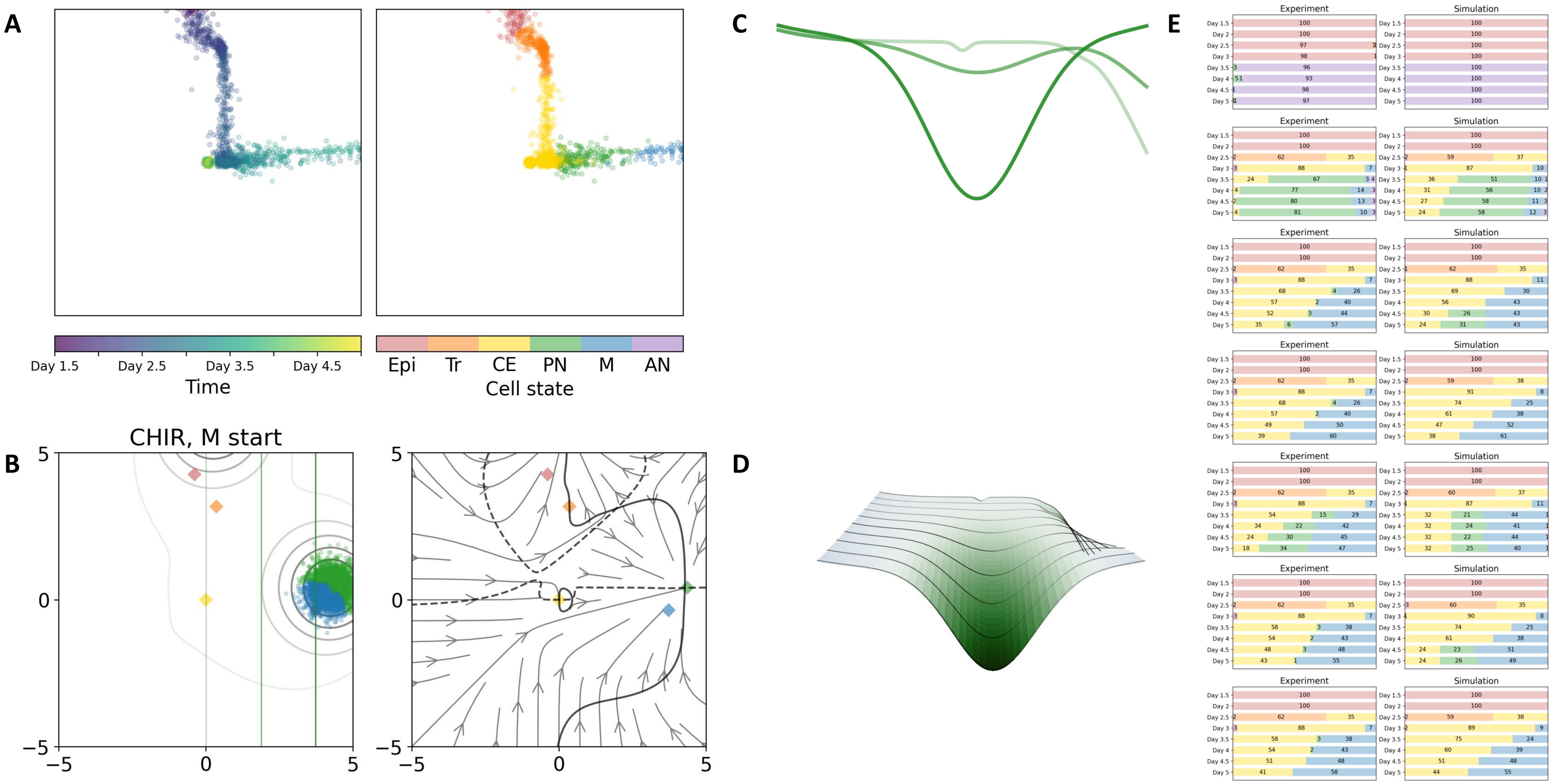

### 43 — Timecode: 20250602-012601

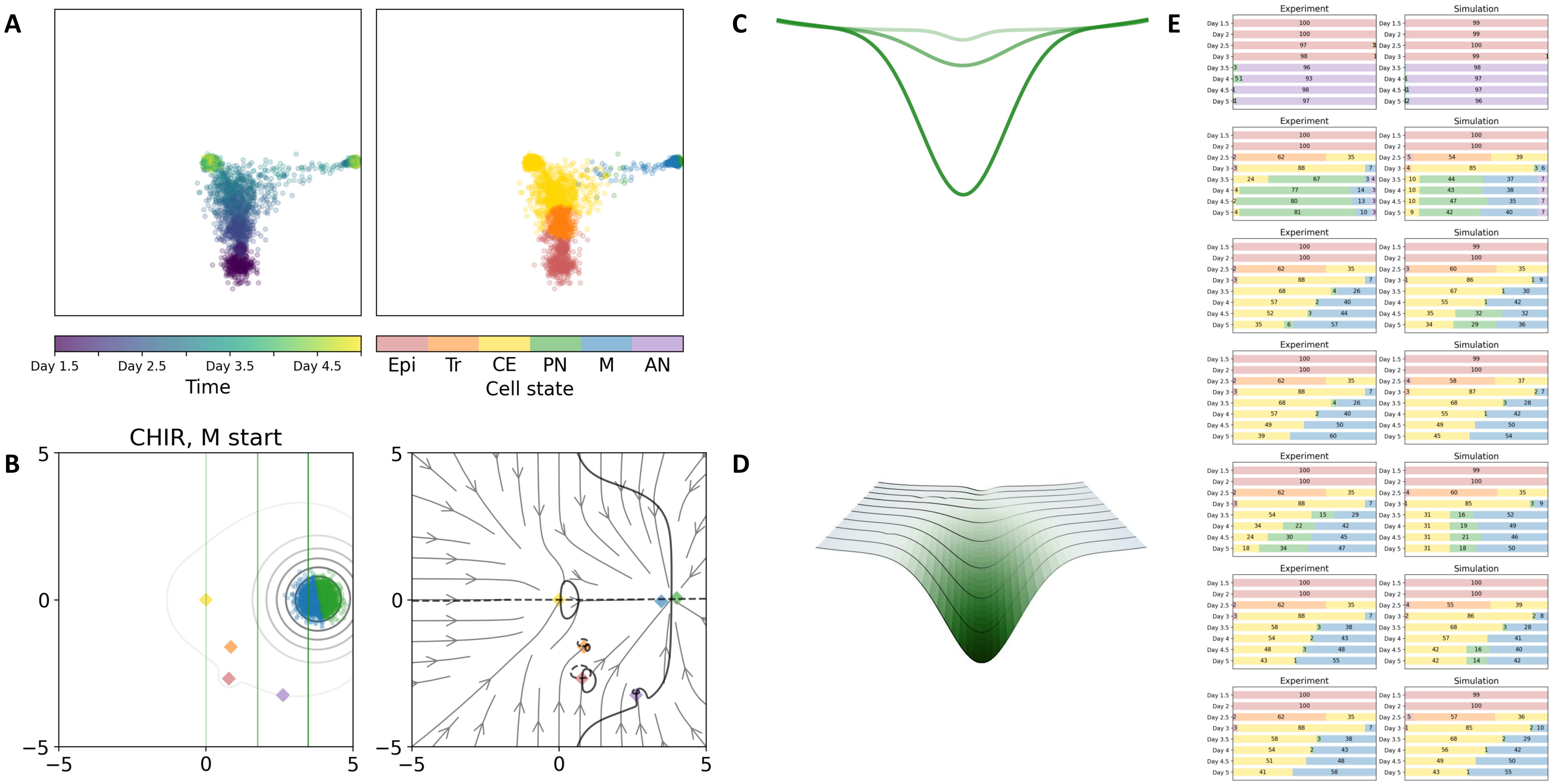

50 — Timecode: 20250602-071802

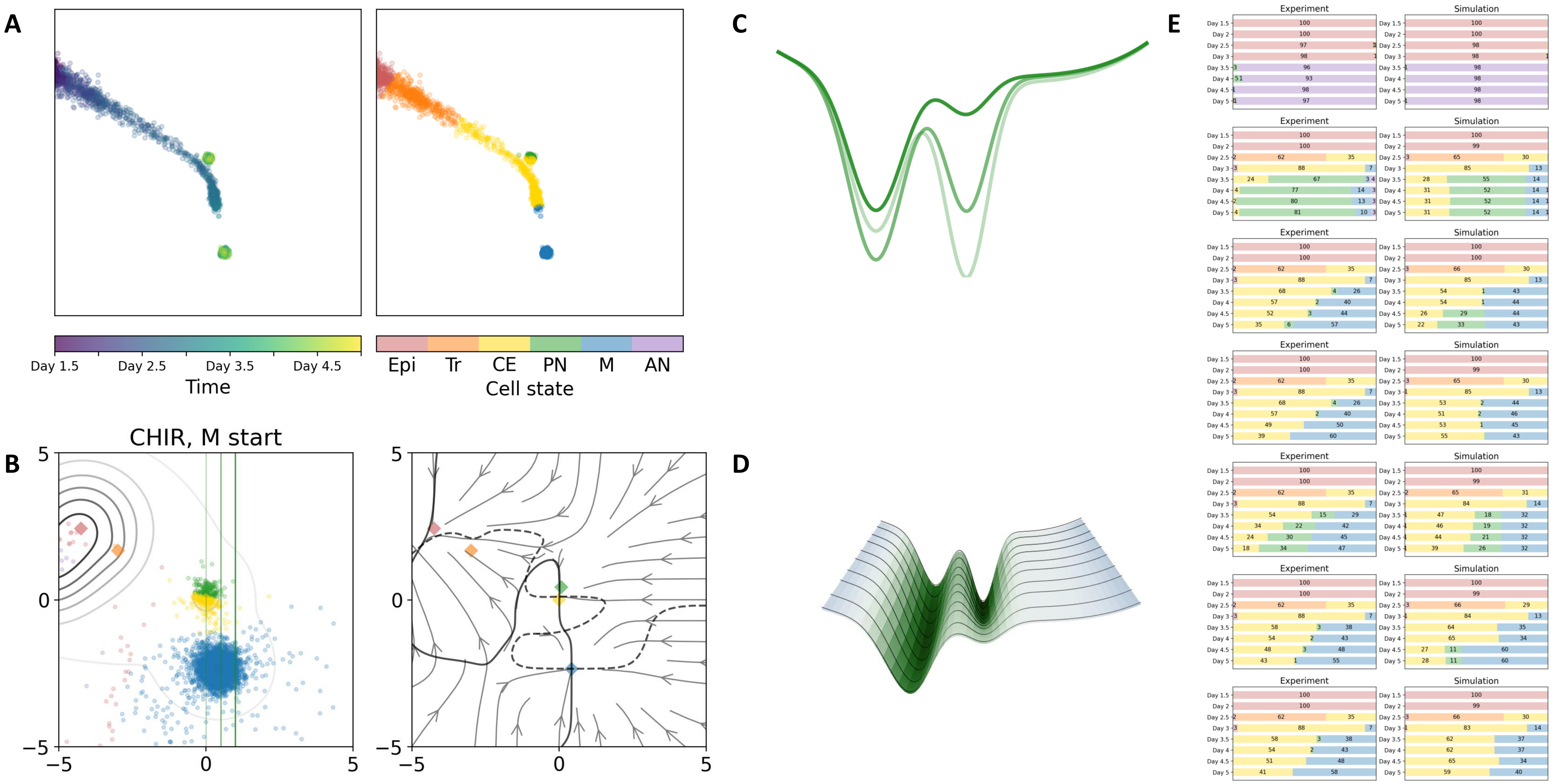

### 44 — Timecode: 20250602-020156

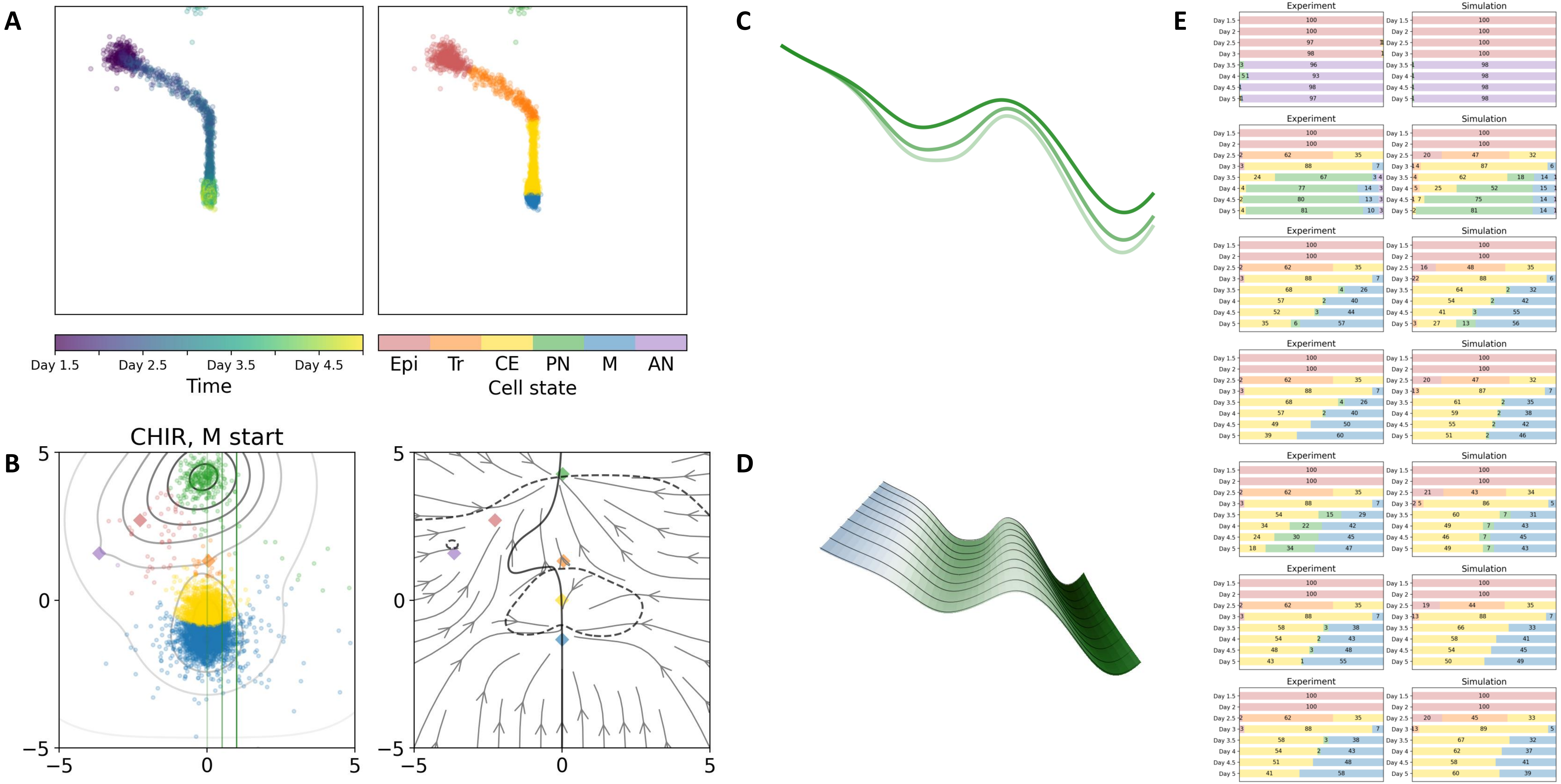
