## Supplementary figures and images for "Generative epigenetic landscapes map the topology and topography of cell fates"

### Supplementary Movie 1

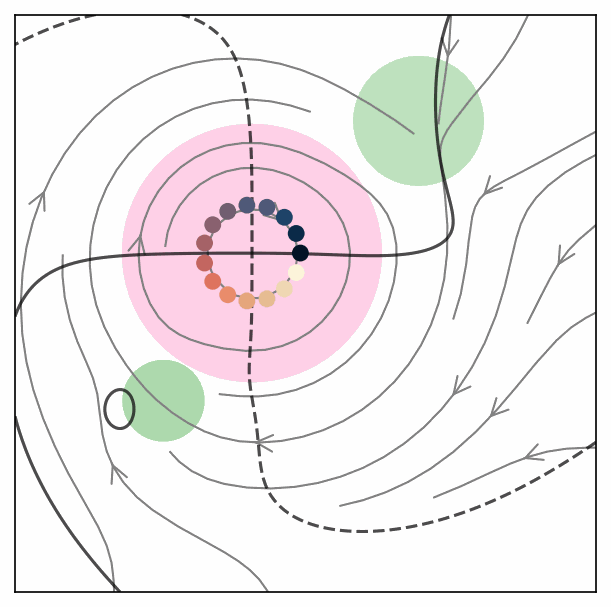

### Supplementary Movie 2

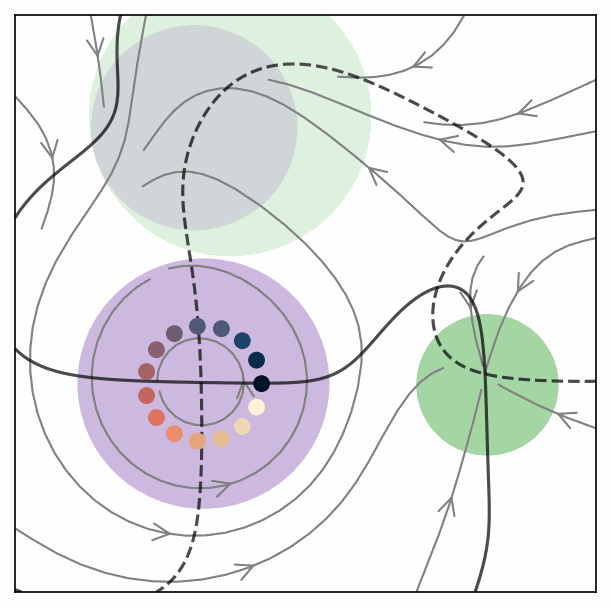

### Supplementary Movie 3

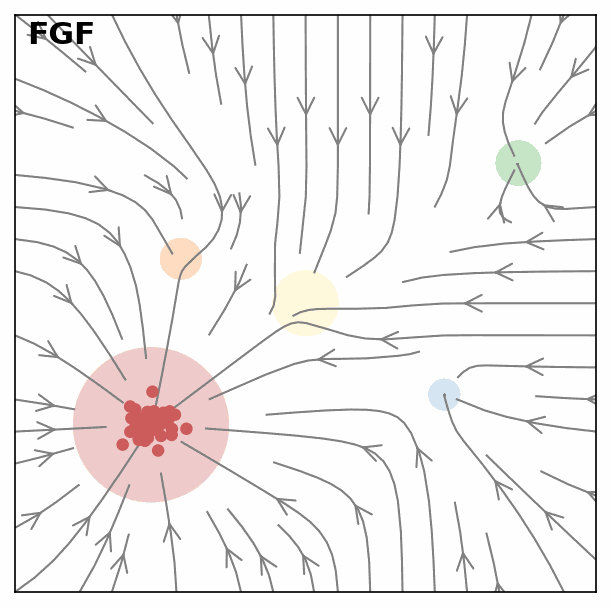

### Supplementary Movie 4

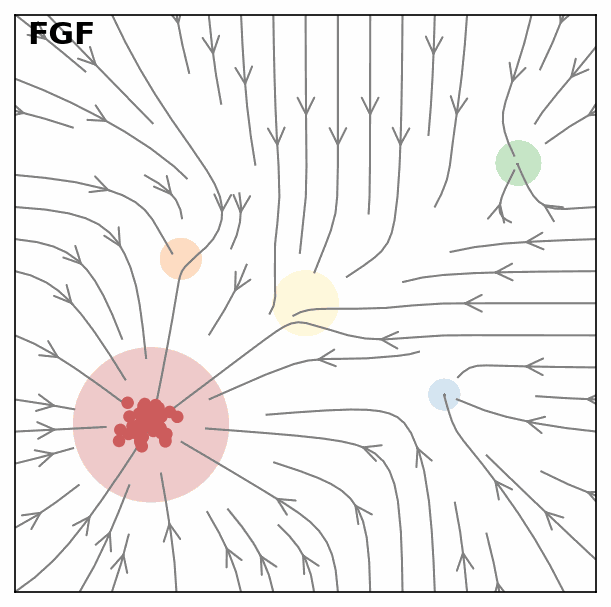

### Supplementary Movie 5

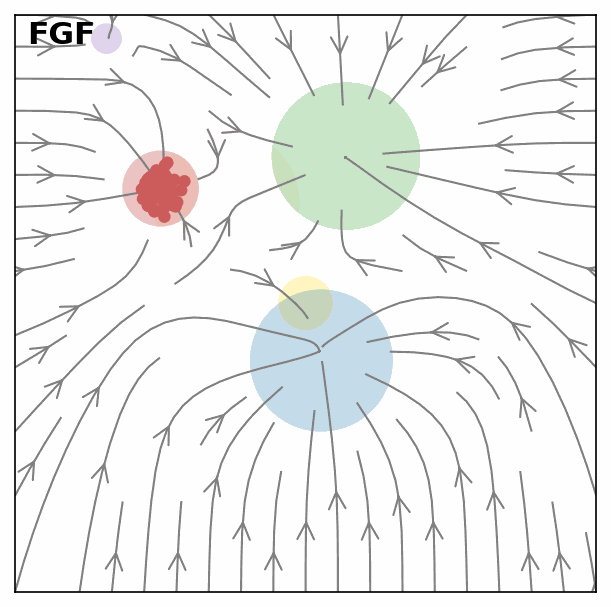

### Supplementary Movie 6

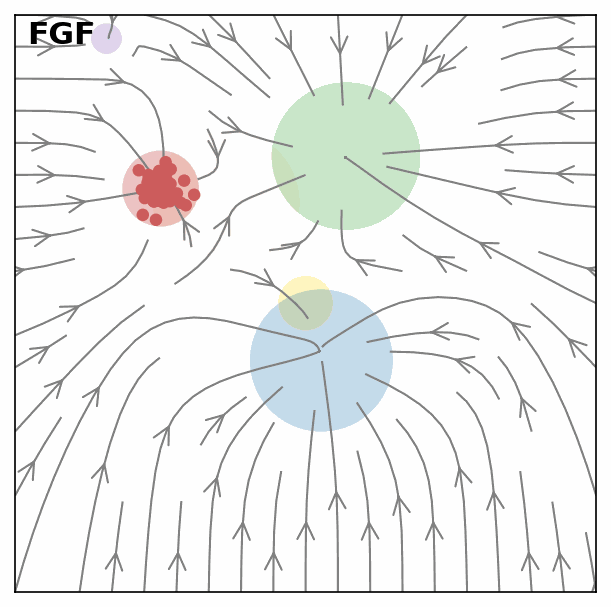
